# Efficient CRISPR genome editing and integrative genomic analyses reveal the mosaicism of Cas-induced mutations and pleiotropic effects of *scarlet* gene in an emerging model system

**DOI:** 10.1101/2024.01.29.577787

**Authors:** Sen Xu, Swatantra Neupane, Hongjun Wang, Thinh Phu Pham, Marelize Snyman, Trung V. Huynh, Li Wang

## Abstract

Despite the revolutionary impacts of CRISPR-Cas gene editing systems, the effective and widespread use of CRISPR technologies in emerging model organisms still faces significant challenges. These include the inefficiency in generating heritable mutations at the organismal level, limited knowledge about the genomic consequences of gene editing, and an inadequate understanding of the inheritance patterns of CRISPR-Cas-induced mutations. This study addresses these issues by 1) developing an efficient microinjection delivery method for CRISPR editing in the microcrustacean *Daphnia pulex*; 2) assessing the editing efficiency of Cas9 and Cas12a nucleases, examining mutation inheritance patterns, and analyzing the local and global mutation spectrum in the *scarlet* mutants; and 3) investigating the transcriptomes of *scarlet* mutants to understand the pleiotropic effects of *scarlet* underlying their swimming behavior changes. Our reengineered CRISPR microinjection method results in efficient biallelic editing with both nucleases. While indels are dominant in Cas-induced mutations, a few on-site large deletions (>1kb) are observed, most likely caused by microhomology-mediated end joining repair. Knock-in of a stop codon cassette to the *scarlet* locus was successful, despite complex induced mutations surrounding the target site. Moreover, extensive germline mosaicism exists in some mutants, which unexpectedly produce different phenotypes/genotypes in their asexual progenies. Lastly, our transcriptomic analyses unveil significant gene expression changes associated with scarlet knock-out and altered swimming behavior in mutants, including several genes (e.g., NMDA1, ABAT, CNTNAP2) involved in human neurodegenerative diseases. This study expands our understanding of the dynamics of gene editing in the tractable model organism *Daphnia* and highlights its promising potential as a neurological disease model.

## Introduction

CRISPR-mediated gene editing systems (Jinek et al. 2012; Cong et al. 2013; Mali et al. 2013) have become a primary tool for introducing DNA sequence modifications in target nuclear genomic regions because of their simplicity and ease of use. When fused with a guide RNA (gRNA), CRISPR nucleases can cause DNA double-strand breaks (DSBs) at a target sequence location that is complementary to the gRNA sequence. Without a DNA repair template, the DSBs can be repaired by the error-prone non-homologous end joining (NHEJ) pathway, resulting in indels and disruption of gene function (Hefferin and Tomkinson 2005; Rodgers and McVey 2016). When a homologous DNA template is provided, DSBs can be repaired through homology-directed recombination (HDR) by template-directed DNA synthesis to bridge the gap across DSBs (Liang et al. 1998; Sekelsky 2017). These features of the CRISPR editing system thus offer flexible control of the genomic locations and outcomes of the genetic modifications.

More importantly, CRISPR-Cas technologies (Jinek et al. 2012; Cong et al. 2013; Mali et al. 2013) have democratized the genetic and genomic research landscape, providing an important means of genetic engineering to emerging model systems that traditionally lack tools for genetic manipulation. The past few years have witnessed the successful implementation of CRISPR-mediated gene editing in organisms such as squids (Crawford et al. 2020), raider ants (Trible et al. 2017), black-legged tick (Sharma et al. 2022), cockroaches (Shirai et al. 2022) and lizards (Rasys et al. 2019), to name a few. Nonetheless, efficiently generating heritable biallelic mutations remains one of the most significant challenges involved in implementing CRISPR-Cas9 gene editing in emerging model systems.

In this study, we present a highly efficient microinjection-based method for generating heritable, biallelic knock-out mutations and evaluate the efficiency for knock-in mutations using the CRISPR-Cas system in the freshwater microcrustacean *Daphnia pulex*. *Daphnia* has been a model system for ecology, evolution, and toxicology for several decades (Miner et al. 2012; Ebert 2022). As the first whole-genome sequenced crustacean species (Colbourne et al. 2011), *D. pulex* has become an important genomics model system for gene-environment interaction (Altshuler et al. 2011), epigenetics (Harris et al. 2012), and evolutionary genomics (Lynch et al 2017). There is also growing interest in using *Daphnia* as a model in studying the evolution of development because *Daphnia* represents an important phylogenetic lineage in invertebrate evolution (Rivera et al. 2010; Mahato et al. 2014; Bruce and Patel 2022). Therefore, an efficient gene editing method would be invaluable for the genetic toolkit of *Daphnia* to unleash its full potential as an emerging model system.

Microinjection into embryos has been a major means of genetic manipulation in *Daphnia*. Under benign environmental conditions female *Daphnia* reproduce asexually through the production of ameiotic diploid embryos that directly develop into neonates in 2-3 days, whereas in stressful environments female *Daphnia* produces haploid eggs and mate with males to produce diploid dormant embryos (**Figure 1**). The asexual reproductive stage provides an excellent platform for microinjection-based genomic engineering. Many asexual embryos can be easily collected for injection from females of the same genotype. After microinjection, the injected embryos can quickly hatch into neonates (G_0_ generation) in 2-3 days, and G_0_ individuals can have asexual progenies (G_1_ generation) in ∼7 days, which guarantees a fast turn-around time for phenotypically and genotypically identifying G_0_ and G_1_ mutants. Furthermore, the asexual reproduction mode allows the long-term preservation of stable mutant genotypes with low maintenance efforts.

**Figure 1.**
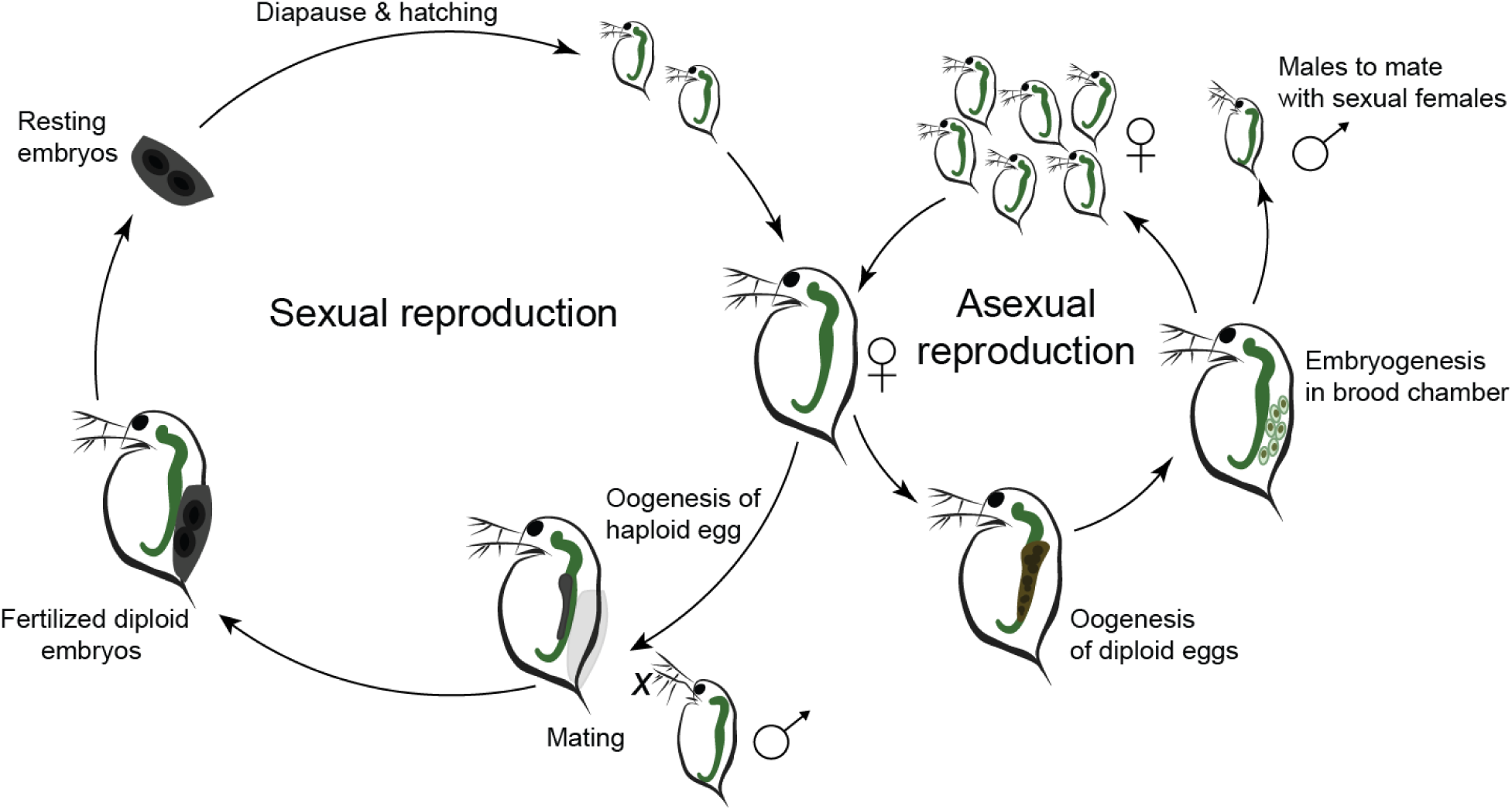
The cyclically parthenogenetic life cycle in *Daphnia*.

Building on these advantages, microinjection techniques have been developed for *Daphnia* to deliver biomolecules into the asexually produced embryos to achieve gene knockdown through RNAi (Kato et al. 2011; Hiruta et al. 2013), protein tagging (Kato et al. 2012), gene knock-out using CRISPR-Cas9 (Ismail et al. 2018), and knock-in using TALEN (Nakanishi et al. 2016). The successful development of these techniques has empowered important discoveries at the molecular level such as the molecular mechanisms of environmental sex determination in *Daphnia magna* (Kato et al. 2018).

However, it should be noted that most of these previous efforts focused on the species *Daphnia magna*, a species has diverged ∼200-million years ago from the focal species of this study *D. pulex* (Colbourne and Hebert 1996). *D. magna* has a substantially larger body size and larger embryo size compared to *D. pulex* (Toyota et al. 2016). Modified microinjection protocols have therefore been developed for *D. pulex* to deal with the technical challenges associated with these variations, for example, internal osmotic pressure in the embryos (Hiruta et al. 2013). However, no efforts have aimed at establishing a microinjection procedure for creating heritable biallelic knock-out and knock-in genotypes using CRISPR-Cas in *D. pulex*. Although knocking out the Hox gene *distalless* was successful in *D. pulex* using CRISPR-Cas9 (Hiruta et al. 2018), no biallelic knock-out lines were created because complete knockout of *distalless* is lethal.

More importantly, given that CRISPR-Cas system can cause spurious off-target mutations or on-target complex mutations in various model systems (Fu et al. 2013; Aryal et al. 2018; Höijer et al. 2022), it is imperative to systematically evaluate the mutations induced by CRISPR-Cas in the *Daphnia* system. However, such studies are still lacking to date.

To create heritable biallelic mutations in an efficient manner, it is critical to accurately deliver RNPs (ribonucleotide proteins - Cas9 fused with gRNAs) or plasmids into the nucleus of an embryo at the one-cell stage. In the hope of accomplishing this at a high efficiency, we first consider the timing of key events in the development of asexually produced embryos in *D. pulex*. During the asexual reproduction in female *Daphnia*, oocytes go through a modified meiosis (i.e., ameiosis) to produce chromosomally unreduced diploid embryos. In ameiosis, the original meiosis I is modified, resulting in suppressed recombination and no cytokinesis, while meiosis II remains normal and produces a polar body and a diploid embryo in the end (Hiruta et al. 2010). These asexually produced embryos can directly develop into neonates in the female’s brood chamber without fertilization.

The cytology of ameiosis in *Daphnia* has been carefully examined (Ojima 1958; Zaffagnini and Sabelli 1972; Hiruta et al. 2010). The ameiotic division begins with the breakdown of germinal vesicle while the eggs are still in the ovary of females, which also coincides with the timing of female molting. Ovulation (i.e., embryos moving into brood chamber from ovary) begins approximately 10-15 minutes after molting. Upon entering the brood chamber, the egg cell enters anaphase I, with chromosomes staying near the periphery of embryo (Ojima 1958; Zaffagnini and Sabelli 1972). The ameiotic division proceeds to anaphase II in approximately 10 minutes post ovulation and the entire division process is completed 20 minutes post-ovulation with polar body emission (Hiruta et al. 2010). At this point the chromosomes move to the deeper part of the embryo, the nucleus membrane re-emerges, and the first cleavage division is finished around 20-60 minutes post-ovulation (Ojima 1958; Hiruta et al. 2010).

Considering this timeline of key events, we suggest that the first 10 minutes post-ovulation (approximately between anaphase I to anaphase II) provides an optimal time window for microinjecting RNPs for introducing biallelic heritable modifications. The oocyte remains in the one-cell stage at this interval, during which the delivered RNPs would have opportunities to bind to chromosomes once the target loci become accessible (e.g., chromosomes exist in a less condensed state) during or after the ameiotic division. Also, microinjecting at an early timepoint is critical for the successful hatching of injected embryos because *Daphnia* embryos rapidly lose their membrane elasticity once ovulated and only early embryos with elastic membranes can sustain the damages caused by microinjection (Kato et al. 2011).

In addition to knowing when to deliver the RNPs alone into the embryo, understanding where in the embryo to deliver the RNPs is also critical for successful gene editing. The small size of the asexual embryos (with a diameter ∼50-100 μm) in *D. pulex* and the presence of massive amount of egg yolk and fat droplets in the embryos make it infeasible to locate the whereabouts of the nucleus or chromosomes during oogenesis under a typical stereomicroscope used for microinjection. Considering that the chromosomes undergo movement from a peripheral spot near the embryo membrane to a more central part of the embryo after the ameiotic division (Ojima 1958; Hiruta et al. 2010), an effective microinjection strategy would be to deliver a concentrated dose of RNPs close to the center of the embryo so that the RNPs can rapidly spread within the embryo to maximize possibilities of binding the targeted chromosomal loci.

Incorporating these considerations, we have developed a set of optimal practices for microinjection experiments for CRISPR-Cas genomic editing in *D. pulex* (**Figure 2**). In this study, we test the efficiency of Cas9 and Cas12a nucleases for generating a heritable biallelic *scarlet* gene knock-out. Also, we use Cas9 to create knock-in alleles at the *scarlet* gene. The SCARLET protein is responsible for transporting tryptophan, precursors of eye pigment (Ewart et al. 1994). Therefore, disruption of the *scarlet* gene can result in clear-eyed mutant daphniids that can be readily distinguished from the wild-type black-eyed individuals (Ismail et al. 2018). We also introduce a few other innovative modifications (e.g., microinjection needles, injection stage) to the existing microinjection system of *Daphnia* to substantially increase its efficiency.

**Figure 2.**
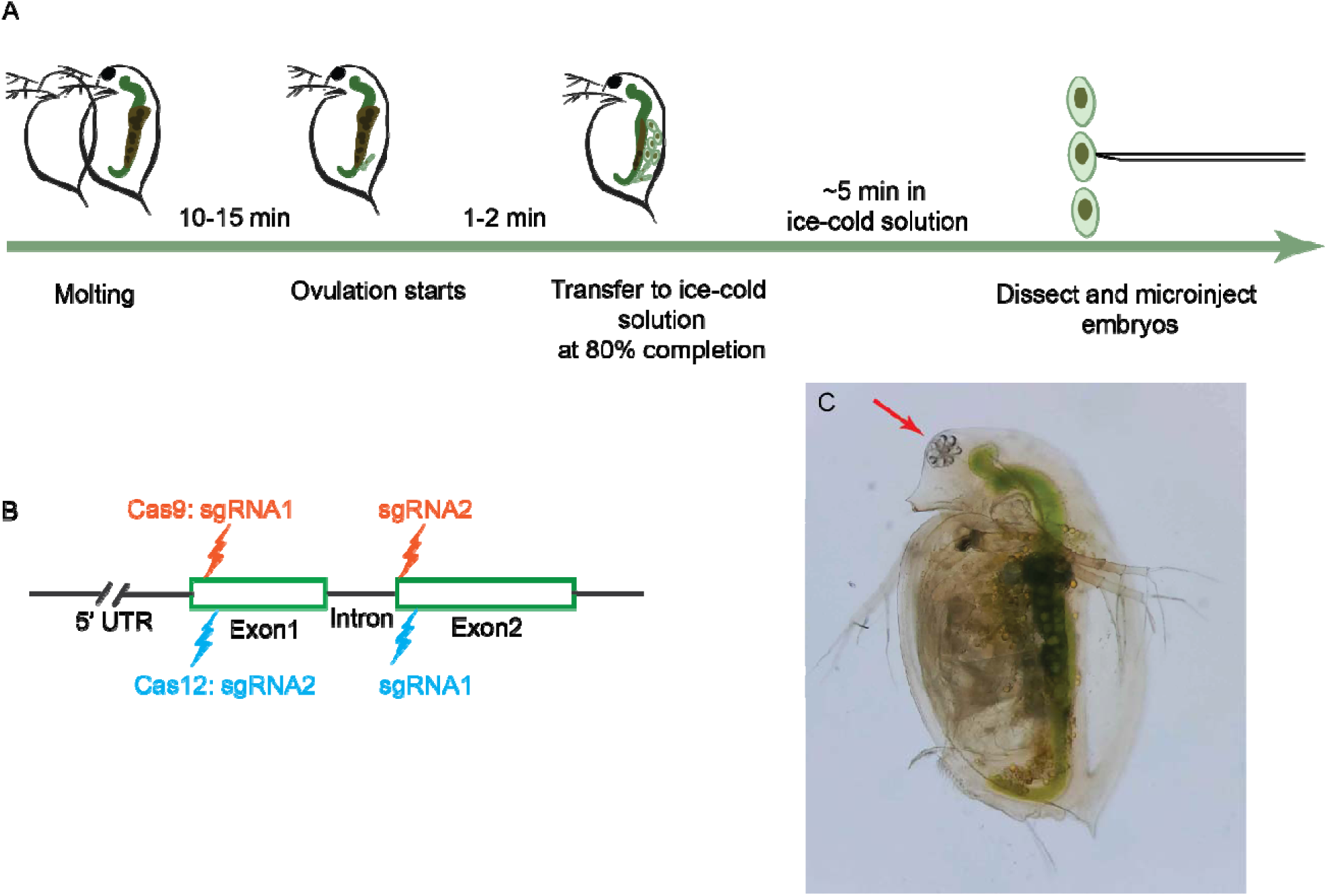
(A) The microinjection workflow for CRISPR-Cas9/Cas12 RNP. (B) the schematic locations for the guide RNA targets for Cas9 and Cas12 gene editing. (C) The clear-eye phenotype (red arrow) caused by the knock-out of scarlet gene.

Furthermore, we analyzed the whole-genome DNA sequences of the knock-out and knock-in mutants to assess the potential of off-target modifications and on-target mutation accuracy in the *D. pulex* genome. Lastly, because knocking out ABC transporters including the *scarlet* gene and *white* gene have pleiotropic impacts on the levels of biogenic amines in the brain (Borycz et al. 2008), male courtship behavior (Anaka et al. 2008), and cyclic GMP transportation (Evans et al. 2008), we examined the altered swimming behavior of *scarlet* mutants and performed RNA-seq experiments to investigate its possible causes and to understand the pleiotropic effects of the *scarlet* gene on genome-wide transcriptomic abundance.

## Materials and Methods

### Experimental animals

We maintained a healthy culture of 2-3-week-old *Daphnia* females that were all asexually derived from a single, natural *Daphnia* isolate EB1 (Eloise Butler, Minnesota). We kept these animals in artificial lake water COMBO (Kilham et al., 1998) at 25 °C and under a 16:8 (light:dark hours) photoperiod. Because we needed asexually reproducing females for collecting asexual embryos, the animals were fed with the green algae *Scenedesmus obliquus* every day and the newly born babies were removed every other day to prevent overcrowding that can trigger *Daphnia* to switch to sexual reproduction.

### Microinjection equipment

We used Eppendorf FemtoJet 4i microinjector and Injectman 4 micromanipulator to perform microinjection on *Daphnia* embryos under a Nikon SMZ800N dissection microscope. We prepared microinjection needles using aluminosilicate glass capillaries (catalog no. AF100-64-10, Sutter Instrument). We chose the aluminosilicate glass rather than regular borosilicate glass because it penetrates the chorion and membrane of *Daphnia* embryos at high efficiency and incurs little clogging at a fine tip size. Microinjection needles were pulled to have a final specification of ∼1.5-μm tip size and ∼7-mm taper length on a P-1000 needle puller (Sutter Instrument), using the following pulling parameters: heat 535 (ramp test value 525 + 10), pull 65, velocity 70, time 200, and pressure 250. The pulled needles were beveled on a BV-10 micropipette beveler (Sutter instrument) with a fine 104D grinding plate (Sutter Instrument) covered by a thin layer of soap water to forge a 30-degree bevel at the tip. The beveled needles were then immediately cleaned using 100% ethanol to remove contaminating debris introduced during pulling and beveling.

### CRISPR-Cas reagents

For the knock-out experiments at the *scarlet* locus using Cas9, we designed one crRNA (**Supplementary Table S1**) targeting the exon 1 and one crRNA for exon 2 using the Design Custom gRNA tool from IDT (Integrated DNA Technologies) based on the DNA sequence of the EB1 isolate (**Figure 2B**). These sgRNAs were chemically synthesized (Alt-R^TM^ custom sgRNAs, IDT). To prepare the RNPs, equal molar amount of each sgRNA and the tracrRNA (Alt-R^TM^ crRNA, IDT) was mixed and incubated at 95 °C for 5 min and cooled to room temperature to form the guide RNA. The guide RNA was subsequently mixed with Cas9 enzyme (catalog no. 1081058, IDT) and was incubated at room temperature for 15 minutes. We co-injected two different RNPs into embryos, with each sgRNA at a concentration of 125 ng/µl and Cas9 enzyme at 600 ng/µl.

For knocking out *scarlet* using Cas12a nuclease (Cpf1), we designed one crRNA targeting a 21-nucleotide sequence for the exon 1 and exon 2 each (**Figure 2B, Supplementary Table S1**). To prepare RNP, the Alt-R^TM^ A.s. Cas12a nuclease V3 (catalog no. 1081068, IDT) was fused with crRNA at equal molar amounts at room temperature. We co-injected two different RNPs into embryos, with each sgRNA at a concentration of 125 ng/µl and Cas12a nuclease at 600 ng/µl.

For the knock-in experiment at the *scarlet* locus using Cas9 nuclease, we designed a HDR template for repairing the double-strand break at the crRNA2 target site (**Supplementary Table S1**). This HDR template was chemically synthesized ssDNA (IDT), containing a stop codon cassette flanked by homology arms of 40 bp on each end. If successfully inserted at the target locus, it would disrupt the translation of the scarlet gene because the stop codon cassette presents stop codons in all the six possible reading frames.

### Embryo collection for microinjection experiments

For each microinjection experiment, we screened our animal culture to select a large number (∼100-150) of females that showed inflated dark ovary, indicating they would likely molt and ovulate in a few hours. Each of the selected animals was placed in a drop of COMBO artificial lake water containing 60 mM sucrose. These animals were regularly checked for signs of molting. The molted animals were closely monitored for signs of ovulation (i.e., oocytes starting to enter the brood chamber), which usually occurs 10-15 minutes after molting (**Figure 2**). Once a female started ovulation and ∼80% of embryos entered the brood chamber (**Figure 2**), we transferred this female to an ice-cold (∼1.5 °C) solution of COMBO with 60 mM sucrose.

We let the female stay in the ice-cold solution for approximately 5 minutes and then dissected the embryos out. We chose the 5-minute wait time because the oocytes should still be undergoing ameiotic division (see Introduction) while the cold temperature slows down the activities of cellular machinery.

The dissection was performed on the bottom surface of a small petri dish flipped upside down (60 mm x 15 mm, catalog no. FB0875713A, Fisher Scientific). After we removed the body of the daphniid, we aligned the embryos against the slightly raised edge of the petri dish. We also removed the excessive solution surrounding the embryos, leaving the embryos submerged under a thin layer of solution.

### Microinjection

We backloaded a microinjection needle with 1 μl of RNP using a microloader tip (Eppendorf, cat no. 930001007). We aimed to inject 1-2 nl of RNP into the embryo. With each needle being different, the injection pressure was generally between 100 and 220 hPa, whereas the background pressure was between 100 and 200 hPa, with an injection time of 0.8 second.

Once the embryos were ready for injection, we immediately performed the injection procedure **(Figure 2**). We injected the RNPs near the center of the embryo. Once the injection was completed, we added COMBO with 60 mM sucrose to the embryos and left the embryos in this condition for ∼30 minutes at room temperature (Cas9) or for 2 hours at 33°C (Cas12a). Then we transferred the injected embryos to a 24-well plate containing COMBO artificial lake water and cultured the embryos at 25°C. Embryos usually hatched into neonates in ∼48 hours at this temperature.

### Mutant screening

Among the hatched neonates from the injected embryos (G_0_ individuals), we searched for individuals with a clear eye or an eye of partially missing black pigment (i.e., mosaic individuals) as tentative knock-out and knock-in mutants. We kept these G_0_ neonates and examined their progenies. If all the asexually produced G_1_ offspring of a G_0_ mutant were clear-eyed, we concluded that the G_0_ individual carried biallelic knock-out mutations at the scarlet locus. We established a clonal mutant line using the G_0_ individual and its asexual progenies.

For the mosaic individuals, we examined the eye phenotype in the different broods of G_0_ individuals. We recorded the number of clear-eye and black-eye individuals and established mutant lines using a single clear-eye individual from the same or different broods.

### Whole-genome sequencing and mutation identification

To investigate whether off-target mutations occur in the knock-out and knock-in mutants, we performed whole-genome sequencing on the asexually produced offspring of each established mutant line plus the wildtype. DNA of mutant lines was extracted from pooled samples of 3^rd^ and 4^th^ generation offspring using a CTAB protocol (Wang and Xu 2021). The DNA sequencing library was prepared by BGI America or by us, and the genome sequencing was performed on a DNB sequencing or Illumina NovaSeq 6000 platform with 150-bp paired-end reads. Each mutant line was sequenced at ∼30x coverage per nucleotide site. Raw reads are available at NCBI SRA (Sequence Read Archive) under project PRJNA1055485.

We aligned the raw reads of each mutant/wildtype line to the *D. pulex* PA42 reference assembly 3.0 (Ye et al. 2017) using the Burrows-Wheeler Alignment Tool BWA-MEM version 0.7.17 (Li and Durbin 2010). We removed reads mapped to multiple locations in the genome and retained only uniquely mapped reads for identifying mutations. We generated calls of single nucleotide variants and indels using the mpileup and call functions of BCFtools (Danecek et al. 2021) for all mutant lines in a single VCF file. Default parameters were used for BCFtools mpileup and call functions, with the addition of the following FORMAT and INFO tags to the VCF file: AD (allelic depth), DP (number of high-quality bases), ADF (allelic depth on forward strand) and ADR (allelic depth on reverse strand). We retained only biallelic single nucleotide polymorphism sites (SNPs) with a quality score (QUAL) >= 20, sequencing depth (DP) >= 10, and a distance >= 50 bp from an indel in each mutant line.

A custom Python script was used to identify mutations in each mutant line using a consensus method. For each SNP site, we established the consensus wildtype genotype call using a majority rule: with a total of N samples in a VCF file, the consensus genotype of a site needs to be supported by at least N-1 samples. If a mutant line shows a genotype different from the consensus genotype, a tentative mutation is identified.

These tentative mutations must meet two criteria to enter the final pool of mutations. First, a mutant allele had to be supported by at least two forward and two reverse reads to avoid false positives due to sequencing errors. Second, a mutant genotype was recognized only when it is a heterozygous genotype derived from a homozygous wildtype genotype. This criterion is to avoid false positives caused by allele dropout due to insufficient sequence coverage or artifacts in library construction at heterozygous sites. This computation pipeline was experimentally verified in a previous study with a false positive rate <0.05 (Snyman et al. 2021).

Furthermore, we examined whether any of the identified mutations in the mutant lines occur within a 100-bp vicinity of any nucleotide sequence that is a blast hit with E value > 0.01 to the gRNA target sequences.

### Structural variation detection

To understand whether the mutants harbored any off-target structural variations caused by Cas nucleases, we used the SV caller Manta (Chen et al. 2016) to identify SVs in all the *scarlet* mutants. Due to the limited power of short-read data in detecting SVs such as inversions, we restricted our analysis to large indels and duplications. In addition to the mutants, we performed genomic sequencing on the wildtype strain. To identify newly arising SVs in the wildtype genomic background, we performed paired analysis between the wildtype and each mutant, where the wildtype served as the control and the mutant represented treatment. All the SV calls excluded imprecise predictions and any variants with a score below 30, thereby mitigating the risk of false positives.

### Behavioral and life-history assay of scarlet mutants

We examined whether the scarlet mutant females (KO2 and KO3) display excessive spinning swimming behavior compared to the wildtype, as reported in the *scarlet* mutant of another *Daphnia* species *D. magna* (Ismail et al. 2021). As we found the wildtype females frequently spins when they try to release babies from their brood pouch, we examined mutant and wildtype females that were 1-day old to 5-day old to avoid this confounding effect. Animals were placed in a 20ml scintillation vial with COMBO artificial lake water under an LED light. We started observing them after 30 minutes of acclimation. We counted the number of spins per minute within a 1-hour window.

### RNA-seq data collection

Transcriptomic sequencing was performed with 2- or 3-day old neonates of seven scarlet mutant lines (KO1, KO2, K03, K04, KI2, KI3, and KI4) and the wildtype to understand how the scarlet knock-out genotype reshaped genome-wide transcript abundance. Three replicate RNA-seq libraries were sequenced for each sample on an Illumina NovaSeq 6000 platform with 150-bp paired-end reads. The raw RNA sequencing data for this project can be found at NCBI SRA under PRJNA1060702. Additional notes about details of our RNA-seq experiments and analyses are available in the Supplementary Materials.

### Differential expression analysis

We examined the raw read quality using FastQC (https://www.bioinformatics.babraham.ac.uk/projects/fastqc/). Adapter trimming and quality filtering were completed using Trimmomatic v.0.39 (Bolger et al. 2014). Trimmed reads were mapped to the *Daphnia pulex* reference genome PA42 3.0 (Ye et al. 2017) using STAR aligner (Dobin et al. 2013) with default parameters. Reads mapping to multiple locations were removed using SAMtools (Li et al. 2009), and the program featureCounts (Liao et al. 2014) was used to obtain the raw read counts for each sample. Differential expression analysis was performed using DESeq2 v.1.34.0 (Love et al. 2014) using the Wald negative binomial test. To obtain an overview of the transcriptomic differences between all *scarlet* mutants and the wildtype, we pooled and compared all mutants against the control samples. Also, we compared the transcriptome of each mutant line with that of the wildtype for differential expression. We adjusted p-values for multiple testing using the Benjamini-Hochberg method as implemented in DEseq2. Genes were considered differentially expressed if they had an adjusted p-value of < 0.05 and greater than a 1.5-fold change. Our scripts are available at https://github.com/Marelize007/Scarlet.

### Co-expression analysis

To identify co-expressed gene modules and to explore the association between gene networks and the scarlet phenotype, we performed Weighted Gene Co-expression Network Analysis (WGCNA, Langfelder and Horvath 2008) to generate a signed co-expression network using the variance-stabilized read counts for the scarlet mutants and wild-type. For computational efficiency, we restricted the analysis to the top 50% of the most variable genes (n=7093). Clusters of highly co-expressed genes were identified by constructing the topological overlap matrices, a measure of interconnection between genes, with a soft cut-off threshold of 14 using the blockwiseModule function (see Supplementary Materials for additional technical details).

Module eigengenes were calculated using the moduleEigengene function. Module eigengenes can be defined as the most representative gene within a module and allow us to study how related the modules are and the correlation between modules and phenotypic traits. An eigengene heatmap **(Supplementary Figure S1)** was constructed to visualize the correlation between modules and the *scarlet* knock-out phenotype.

Lastly, we constructed a network of differentially expressed genes in the red module to visualize the correlations between them **(Supplementary Figure S2)**. Negative correlation edges were colored black and positive correlation edges were colored red. Only edges with a Pearson correlation coefficient greater than 0.9 were kept. The size of the vertices was scaled in proportion to the level of expression of each gene. The genes were further clustered into communities based on edge betweenness.

### Functional enrichment analysis of differentially expressed genes

We performed GO term enrichment analysis using the R package topGO (Alexa and Rahnenfuhrer 2019) to investigate the biological relevance of differentially expressed genes. The default algorithm, weight01, was used along with the Fisher’s exact test.

Furthermore, to detect KEGG pathways enriched for differentially expressed genes, the GHOSTX program (Moriya et al. 2007) was used to query all the annotated (18,440) genes from the *D. pulex* PA42 transcriptome (Ye et al. 2017) in the KEGG Automatic Annotation Server (KAAS). A total of 10,135 genes were assigned a KO (KEGG ortholog) number, of which 6,282 were mapped to KEGG pathways. Enriched pathways were identified through hypergeometric tests with Holm-Bonferroni correction in a custom script (https://github.com/Marelize007/Scarlet). KEGG pathways with a p-value < 0.05 were considered significantly enriched with differentially expressed genes.

## Results

### Optimized microinjection methods

Our microinjection method featured a few notable improvements, empowering fast and efficient microinjection with *Daphnia*. Our needle pulling and beveling procedure consistently generated injection needles with a tip size of 1-2 µm and 5-6 mm taper. At such fine tip size, the beveled needle had substantially fewer clogging issues than non-beveled ones, whereas aluminosilicate-glass needles allowed easier penetration into the embryos than borosilicate-glass needles. Moreover, dissecting and injecting embryos on the same petri dish stage produced an increased number of intact embryos for injection, compared to a procedure involving transferring the newly ovulated embryos (extremely fragile with an irregular shape) from a dissection stage to the injection stage (which would inevitably damage some embryos). To clearly demonstrate our microinjection procedure, a video is available through this link (https://youtu.be/z1Dc0vTAj8A).

### Knock-out and knock-in efficiency at the scarlet locus

Using our optimized microinjection procedure, we were able to consistently perform successful microinjection of Cas9/Cas12a RNPs into the asexual embryos of *Daphnia* and achieve a hatching rate of 24-59% for the injected embryos (**Table 1 and Table 2**). In *scarlet* CRISPR-Cas9 knock-out experiments, the success rates for generating the clear-eye phenotype in G_0_ individuals (**Figure 2C**) ranged from 1% to 15.6% with a median of 3.85%, whereas in the knock-in experiments the clear-eye phenotype rate was between 1.1% and 3.4% (**Table 1**). Moreover, we identified mosaic mutants among G_0_s that have reduced black pigments in the eyes (**Table 1**). For the Cas12a knock-out experiments, the success rates for generating clear-eyed G_0_ individuals were between 0 and 16%, whereas the rate for mosaic individuals ranged between 0 and 19%.

**Table 1.** Summary of CRISPR-Cas9 knock-out (KO) and knock-in (KI) experiments for the scarlet gene.

| Experiment | No. of injected embryos | No. of hatching embryos (percentage) | No. of clear-eye Phenotype (percentage) | No. of mosaic phenotype (percentage) |
| --- | --- | --- | --- | --- |
| KO | 80 | 28 (35.0%) | 1 (3.6%) | - |
| KO | 176 | 45 (25.6%) | 7 (15.6%) | - |
| KO | 147 | 67 (45.6%) | 1 (1.5%) | - |
| KO | 63 | 24 (38.1%) | 1 (4.1%) | - |
| KO | 107 | 35 (32.7%) | 2 (5.7%) | 1 (2.9%) |
| KO | 245 | 100 (40.8%) | 1 (1.0%) | - |
| KI | 140 | 80 (57.1%) | 1 (1.3%) | - |
| KI | 200 | 117 (58.5%) | 4 (3.4%) | 3 (2.6%) |
| KI | 235 | 130 (55.3%) | 2 (1.5%) | - |
| KI | 204 | 92 (45.1%) | 1 (1.1%) | - |

**Table 2.** Summary of CRISPR-Cas12 experiments for the scarlet gene.

| Experiment | No. of injected embryos | No. of hatching embryos (percentage) | No. of clear-eye Phenotype (percentage) | No. of mosaic phenotype (percentage) |
| --- | --- | --- | --- | --- |
| KO | 59 | 28 (47.5%) | 0 | 2 (7.1%) |
| KO | 75 | 28 (37.3%) | 1 (3.6%) | 2 (7.1%) |
| KO | 131 | 31 (23.7%) | 1 (3.2%) | - |
| KO | 93 | 25 (26.7%) | 4 (16%) | - |
| KO | 57 | 21 (36.8%) | 1 (4.7%) | 4 (19.0%) |
| KO | 69 | 17 (24.6%) | 0 | 2 (11.8%) |

From our knock-in experiments, two out of eight clear-eyed G_0_s were found to have successful insertion of the stop codon cassette at the scarlet locus. However, the insertion of the stop codon was accompanied by complex modifications at the target site (see below).

### Transmission pattern of scarlet knock-out genotypes

For all the established clear-eyed and mosaic G_0_ individuals, we examined the eye phenotype of their asexual offspring from different broods (G_1_ individuals). For all the clear-eyed G_0_s from Cas9 knock-out/knock-in experiments, all the G_1_s from at least three different broods showed the clear-eye phenotype, indicating heritable biallelic modifications at the scarlet locus. Moreover, we examined the asexual G_2_ offspring from a random set of clear-eyed G_1_s, and all the G2s showed the same phenotype as their mothers, as expected under asexual reproduction.

For the Cas9-produced mutants, we occasionally found a mosaic individual produced only black-eyed offspring, indicating somatic mutation at the *scarlet* locus. Nonetheless, most mosaic individuals produced clear-eyed offspring, sometimes along with black-eyed siblings. We noted that no offspring from mosaic individuals exhibited the mosaic phenotype. Some mosaic individuals produced only clear-eyed offspring, while others interestingly produced both clear-eyed and black-eyed offspring in the same or different broods (**Figure 3 and Supplementary Table S2**). This observation led us to hypothesize that the asexual offspring of *Daphnia* originated from different oogonia, some of which were genetically modified by CRISPR-Cas9 and others were not. This also raised the possibility that the genetically modified oogonia may carry different genetic modifications at the scarlet locus, i.e., mosaicism in the germline cells.

**Figure 3.**
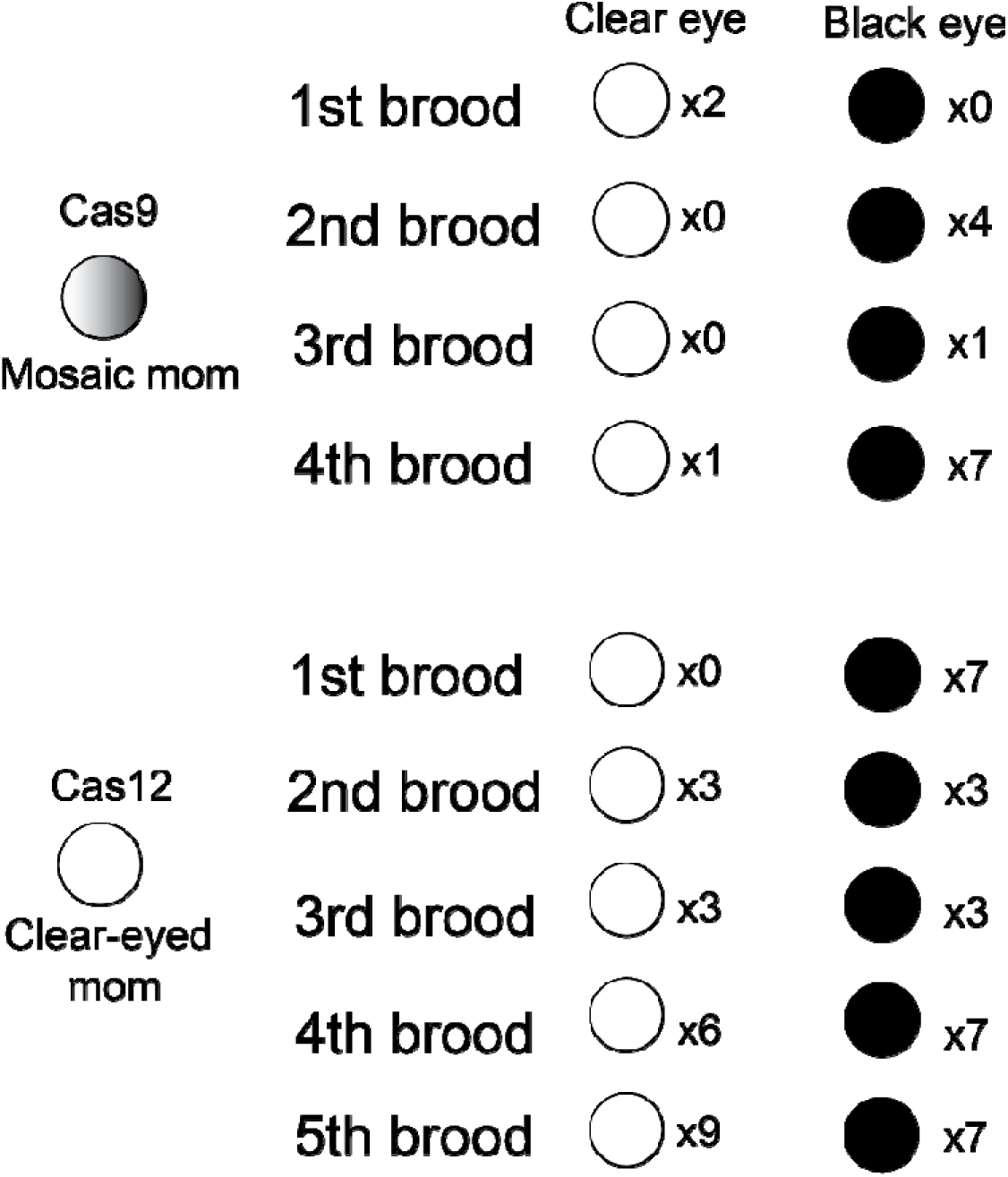
Examples of transmission pattern of scarlet knock-out genotype.

On the other hand, we saw an increase of mosaic G_0_ individuals (5 out of 8) from Cas12a knock-out experiments. These mosaic individuals produced only black-eyed G_1_s in all the examined broods during their life span (**Table S3**). This observation suggests increased somatic knock-out of *scarlet* with Cas12a nuclease under our experimental conditions. However, germline knock-out did occur in Cas12a experiments. Some mosaic G_0_ individuals and a clear-eyed individual produced mixed broods (**Figure 3 and Supplementary Table S3**), strongly indicating germline mosaicism in the G_0_s.

### Knock-out genotypes induced by Cas9 nuclease

We used the whole-genome short-read sequences of the mutants to analyze the Cas9-induced mutations at the *scarlet* locus. In our analyses we were able to phase the two haplotypes of the scarlet locus and identify the haplotype-specific mutations. Detailed mutation sequence alignment is available in **Supplementary File 1**. Our analyses showed that, although RNPs targeting two different sites of the *scarlet* locus were co-injected, only one knock-out mutant (KO4) contained a homozygous segmental deletion (234 bp) spanning the two target sites (**Figure 4A**), indicating the simultaneous occurrence of DNA double-strand breaks induced by CRISPR-Cas9 at these two target sites in the single-cell stage. Many of the other knock-out mutants showed small indels at the target sites. For example, at target site 2 a 13-bp insertion on allele 1 and a 2-bp insertion and a 3-bp substitution on allele 2 were identified in the mutant KO1, most likely because of NHEJ repair (**Figure 4B**). Across all the mutants the lengths of insertions ranged between 5 and 29 bp, whereas that of deletions varied from 1 to 16 bp (**Supplementary Table S4**).

**Figure 4.**
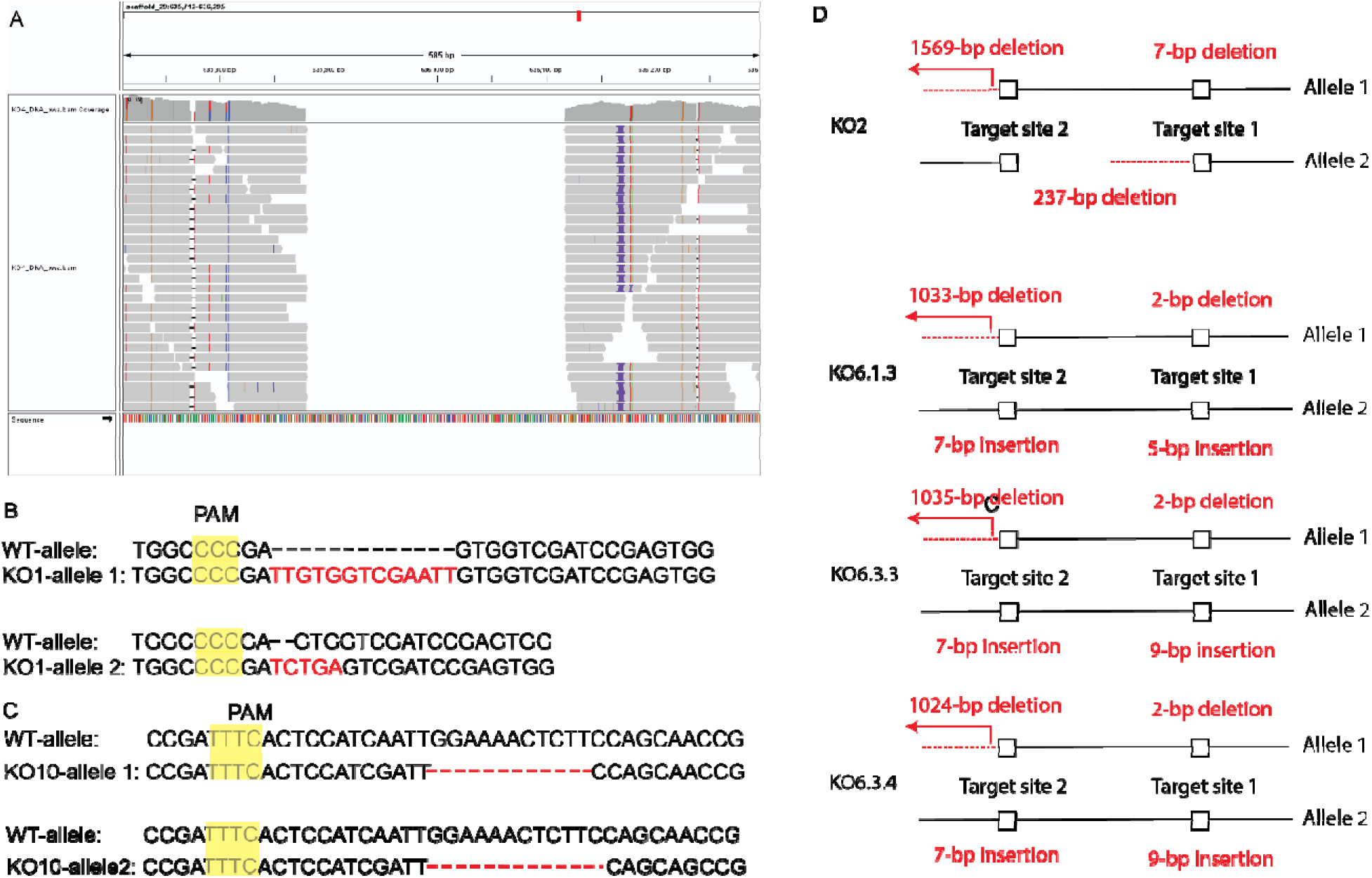
Examples of Cas9/Cas12 induced mutations. (A) Short-sequence alignment around the homozygous 234-bp segmental deletion in mutant KO4. Segmental deletion is inferred because of the absence of reads (denoted by grey bars). (B) Cas9-induced allele-specific mutations (red) at target site 2 in mutant KO1. (C) Cas12-induced allele-specific mutations (red) at target site 2 in mutant KO10. (D) Mutants with unexpected, on-target hemizygous deletions.

We noted that the modifications of two alleles at the same target sites were different in all mutants (**Supplementary Table S4**). For example, two alleles could have indels of different sizes, or one allele had an insertion whereas the other allele had a deletion, indicating the DNA repair outcome was often independent between the two alleles. Intriguingly, although induced mutations occurred at both target sites, no segmental deletions were generated in most of these knock-out mutants. This was most likely because the induction of DNA double-strand breaks at different target sites occurred in a sequential manner rather than simultaneously during cell division.

We identified a few mutants that carried unexpected, large deletions (>50bp) at the *scarlet* locus (**Figure 4D**). For example, in mutant KO2 allele 2 had a segmental deletion of 237 bp spanning the two target sites, whereas allele 1 had a 1569-bp deletion upstream of the target site 2 (**Figure 4D**). In the mutants KO6.1.3, KO6.3.3, and KO6.3.4, allele 1 had a 1033-bp, 1035-bp, and 1024-bp deletion upstream of the target site 2, respectively (**Figure 4D**). It seems allele 1 at target site 2 was more vulnerable to large deletion than allele 2, as no large deletions were found at allele 2 in any mutants.

The presence of both black-eyed and clear-eyed neonates in the asexual broods of the mosaic individuals led us to hypothesize that the asexually produced neonates of the same mother could be derived from primary oocytes with different or no genetic modifications caused by CRISPR-Cas9 (i.e., mosaicism in the germline cells). To test this hypothesis, we examined the genotypes of the clear-eyed neonates from the same or different broods of the same mosaic mother (KO5 and KO6). Within the same brood, neonates in the first brood of KO5 (KO5.1.1 and KO5.1.2) and the first three broods of KO6 (first brood-KO6.1.1, KO6.1.3, KO6.1.4; second-KO6.2.1, KO6.2.2; third-KO6.3.2, KO6.3.3, KO6.3.4) all carried different genotypes, whereas the fifth brood of KO6 (KO6.5.1, KO6.5.2, KO6.5.3) had the same genotype for all neonates (**Supplementary Table S4**). These observations strongly supported our hypothesis. Across the multiple broods of KO6, we observed the presence of different knock-out genotypes, substantiating the notion of mosaicism in germline cells. Also, the same genotype appeared in different broods of the KO6 individual (KO6.3.2 and KO6.4.1), suggesting these neonates were likely derived from the same primary oocyte cells or different primary oocytes of the same genetic modification.

### Knock-out genotypes induced by Cas12a

Compared to Cas9-induced mutants, our collection of Cas12a-induced knock-out mutants showed distinct patterns of genetic modification at the two target sites in *scarlet*. We detected only deletions at the target sites, with their sizes ranging from 1 to 73 bp (**Supplementary Table S4**). Moreover, in three of the four mutants more than one allele-specific target site remained unmodified, whereas for Cas9-induced mutants only one (KO1) out of 18 mutants had a single site unmodified (**Supplementary Table S3, S4**). For detailed mutation sequence alignment, see Supplementary File 1.

### Knock-in genotypes at the scarlet locus

Out of the five mutants from our knock-in experiment at target site 2 (mutation sequence alignment available in Supplementary File 2), we identified two clear-eyed mutant lines (KI4, KI6) with complete knock-in of the stop-codon cassette in allele 1 based on our reconstruction of the haplotypes with short-read data. The allele 2 in both KI4 and KI6 harbored a 7-bp insertion at the target site (**Figure 5**). However, the identified knock-in events were accompanied by complex local genomic rearrangements in both mutants. In KI4, the insertion of a 240-bp segment containing the stop-codon cassette with multiple duplicated copies of homology arms occurred at the location of a 635-bp deletion covering the majority of the target site, resulting in a net loss of 395 bp (**Figure 5**). In KI6, two versions of allele 1 with knock-in insertion were observed. In retrospect, this was most likely due to germline mosaicism and complicated our reconstruction of the insertion haplotypes. Our short-read genomic data allowed us to completely reconstruct only one haplotype (KI6-hap1), leaving the other one partially resolved (KI6-hap2). In KI6-hap1 (**Figure 5**), a 116-bp segment containing a stop-codon cassette and a homology arm was inserted to replace a 635-bp segment (the same deletion tract observed in KI4). For KI6-hap2, an insertion > 127bp containing multiple duplicated copies of one homology arm occurred at the target site, accompanied by the deletion of original sequence of unknown length.

**Figure 5.**
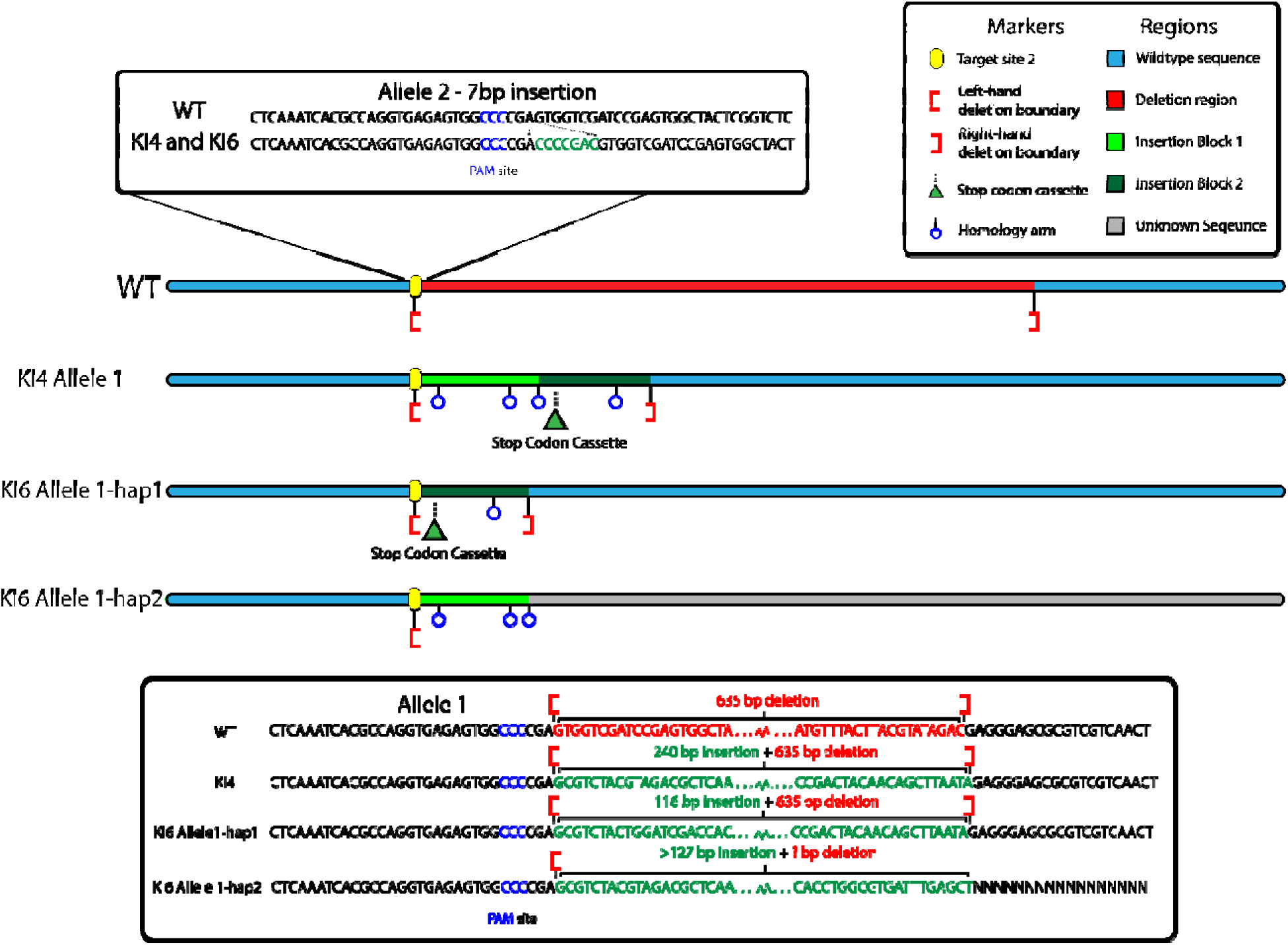
Illustration of the genomic rearrangements accompanied the knock-in of stop codon cassette in the scarlet locus in two mutant lines, KI4 and KI6.

### Genome-wide mutation rate in scarlet mutants

To determine whether the *scarlet* mutants showed an elevated level of base-substitution mutations in comparison to the wildtype and whether any mutations were due to the mistarget effect of Cas9/Cas12a, we examined the genomic DNA sequences of all *scarlet* mutant lines derived from their 3^rd^/4^th^ generation offspring. Using a stringent germline mutation detection procedure that was experimentally tested with a false discovery rate < 5% (Snyman et al. 2021), we identified base substitutions in 7 mutant lines with 1-3 base substitutions in each line, whereas the other 17 mutant lines (including the wildtype) had no base substitutions (**Supplementary Table S7**). None of the base substitutions occurred within or near the possible mistarget sites of Cas9/Cas12a (with 1 or 2 mismatches to the target sites). The base substitution rate was on the order of 10^-9^ per site per generation, on par with the spontaneous mutation rate in *Daphnia* (Keith et al. 2016; Flynn et al. 2017), indicating no elevation of base substitution rate caused by gene editing.

Similarly, our analysis of structural variations (SVs) did not reveal an elevated number of large indels and duplications. In 20 out of 28 mutant lines, no SVs were detected. Only one monoallelic insertion event was detected in one mutant (KI6), whereas six hemizygous deletion events were identified with lengths ranging from 54 to 23,418 bp (**Supplementary Table S8**) in five mutants. Notably, a 23418-bp hemizygous deletion occurred in KO9. Moreover, two duplication events were found (297bp and 1735bp). None of these events occurred in genic regions except for a deletion event in mutant KO5.1.1 overlapping part of gene3461 (kinesin-related protein 12 signal transduction). Considering that large-scale hemizygous deletion, insertion and duplications (Xu et al. 2011; Keith et al. 2016) occurs at high rates in the asexual reproduction in *Daphnia* (on the order of 10^-5^/bp/generation), the observed deletion rates (ranging from 1.12 x10^-7^ to 4.86 x 10^-5^ /bp/generation), insertion rate (3.07x10^-7^ /bp/generation), and duplication rates (from 6.17 x 10^-7^ to 3.6 x 10^-6^) were below or on par with the spontaneous rates (**Supplementary Table S9**).

### Spinning behavior in scarlet mutants

Our daily observations of the wildtype vs *scarlet* mutants (day 1 to day 5 post birth) revealed that while the wildtype swam up and down with small hops and rarely spun themselves, the spinning behavior exacerbated as the mutant neonates grew up (**Figure 6A**). The movement of mutant female neonates was a combination of normal movement interspersed by episodes of fast spinning, with the spins on day 1 averaging 7.5 in one minute increasing to 63.1 in one minute on day 5. We also observed that some mutants tend to perch at the bottom of the vial on their abdomen or back after fast spinning, which was rarely observed in the wildtype.

**Figure 6.**
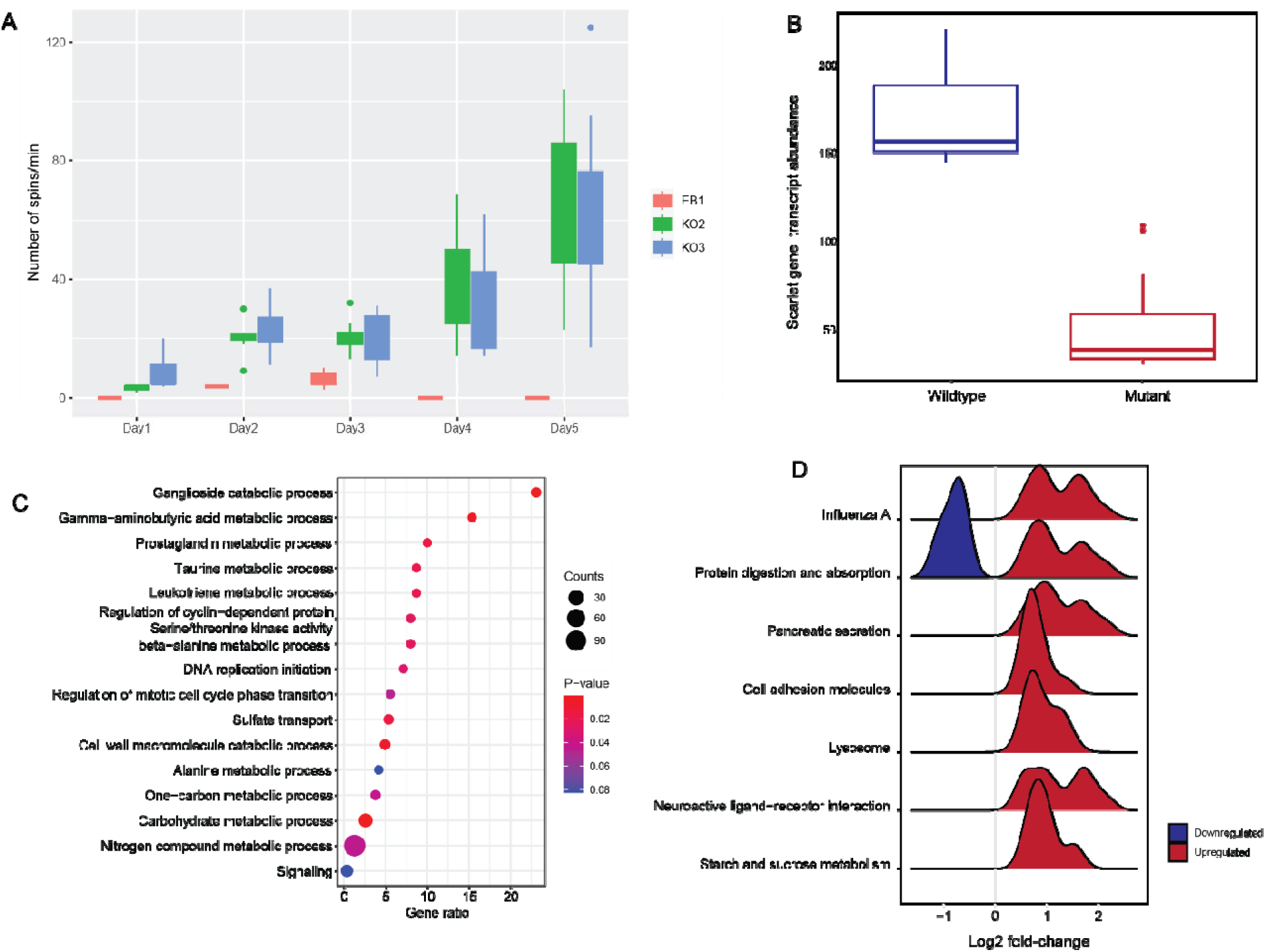
(A) The number of spins observed in scarlet mutants KO2 and KO3 in the 8 days post hatching. (B) The transcript abundance of scarlet in wildtype and mutants. (C) Top ranked GO terms from the GO enrichment analysis. Gene ratio is calculated as counts divided by the expected number of genes. (D) The Log2fold distribution of differentially expressed genes in the significantly enriched KEGG pathways. The height of peak represents the number of differentially expressed genes.

### Differential gene expression

We performed RNA-seq analyses with three biological replicates of seven *scarlet* mutants each and the wildtype, totaling 24 samples. An average of 20.9 million (SD=3.5 million) reads were sequenced per sample. Approximately 99% of the raw reads passed our quality control and trimming. On average 78% (SD=4.6%) of the retained reads uniquely mapped to the *D. pulex* reference genome and were used for downstream analyses (**Table S10**).

Our pooled differential expression analysis contrasting all mutants’ replicates against the wildtype revealed 328 significantly upregulated genes and 112 genes downregulated in the scarlet mutant lines compared to the wildtype (**Supplementary Table S11, Supplementary File 3**). The number of upregulated genes was significantly higher than downregulated genes (chi-square test p<0.0001). Among these genes, the *scarlet* gene was significantly downregulated in the mutants (**Figure 6B**). GO term enrichment analysis showed that various metabolic and catabolic processes comprised most of the top-ranked GO terms (**Figure 6C, Supplementary Table S12**), with a notable exception of cyclin-dependent kinase serine/threonine kinase activity, sulfate transport, DNA replication initiation, and signaling. This indicated that the pleiotropic effects of scarlet knock-out reached beyond the metabolic processes.

Through the KEGG pathway enrichment analysis of differentially expressed genes, we found that the significantly enriched pathways contained Protein digestion and absorption, Pancreatic secretion, Influenza A, Lysosome, Neuroactive ligand-receptor interaction, Cell adhesion molecules, and Starch and sucrose metabolism (**Figure 6D**, **Supplementary Table S12**). Nearly all the significantly enriched pathways consisted of only up-regulated genes, with a mix of up- and down-regulated genes in Protein digestion and absorption (**Figure 6D**).

Considering the altered swimming behavior of *scarlet* mutants and neurotransmitter deficiency noted in the *scarlet* mutant of *D. magna* (Ismail et al. 2021), we further examined the KEGG pathways relevant for movement and neurological transmission, which were Neuroactive Ligand-receptor Interaction and Cell Adhesion Molecules. In Neuroactive Ligand-receptor Interaction, differentially expressed genes included multiple paralogs of trypsin genes, neuropeptides capa receptor-like gene, and glutamate receptor ionotropic NMDA1. It should be noted that NMDA1 is involved in neurodevelopmental disorders with movement abnormalities. For Cell Adhesion Molecules, eight paralogs of contactin-associated protein-like 2 (CNTNAP2) were upregulated. CNTNAP2 has been found to be associated with multiple neurological disorders including Autism (Alarcon et al. 2008) and Pitt-Hopkins syndrome (Peippo and Ignatius 2011).

Furthermore, we identified a few non-enriched KEGG pathways that contain highly differentially expressed genes. In the GABAergic synapse pathway, gene ABAT was significantly upregulated almost 4-fold in *scarlet* mutants. ABAT gene is known to be involved in human disease Encephalopathy with uncontrolled limb movements and exaggerated reflexes (Louro et al. 2016; Koenig et al. 2017). The abovementioned glutamate receptor ionotropic NMDA1 was 1.6-fold upregulated in *scarlet* mutants and is also associated with the amyotrophic lateral sclerosis (ALS) pathway.

In the individual analysis of each *scarlet* mutant in comparison to the wildtype, the number of significantly upregulated genes in mutants ranged from 453 to 1111, whereas that of significantly downregulated genes was between 250 to 711 (**Supplementary Table S11**). Moreover, we generated the consensus set of differentially expressed genes that were shared by all mutants (i.e., the same direction of expression change > 1.5 fold), consisting of 26 up-regulated and 5 down-regulated genes (**Supplementary Table S13**). Many genes on this list were involved in various metabolism pathways (**Supplementary Table S14**). More importantly, it corroborated several genes that emerged from the pooled analysis including ABAT, CNTNAP2, and a few paralogs of trypsin. It also drew our attention to a downregulated gene in the *scarlet* mutant, gene4054 (SLIT2), which is important for axon regeneration and axon guidance, two fundamental processes in the nervous systems (Curcio and Bradke 2018). We noted that another gene, gene3375 (SLIT3), which is involved in axon regeneration and axon guidance, did not enter the consensus list because in one mutant its fold change did not exceed 1.5. However, SLIT3 appeared to be co-regulated with scarlet in our co-expression analysis (see below).

### Gene modules and co-expression analysis

With our gene expression data, a total of 14 gene co-expression modules were obtained. For each module, module eigengenes were Spearman rank correlated with the *scarlet* knock-out phenotype. Among the 14 modules identified, six were significant in their correlations with the *scarlet* mutant phenotype (**Supplementary Figure S1**). The positively correlated modules (**Supplementary Figures S2-S6**) were magenta (ρ = 0.43, p = 0.04), red (ρ = 0.76, p = 1x10^-5^), yellow (ρ = 0.52, p = 0.009), and salmon (ρ = 0.6, p = 0.002) modules, while negatively correlated ones were: purple (ρ = -0.58, p = 0.003) and grey (ρ = -0.48, p = 0.02).

Out of these modules, the red module contained genes that were the most differentially expressed between the wild-type and *scarlet* knock-out mutant lines. A network was constructed of the differentially expressed genes in this module to visualize the correlations between them (**Supplementary Figure S2**). The scarlet gene was most positively co-expressed with gene CTRL (chymotrypsin-like protease CTRL-1), which functions as a protease and hydrolase, whereas CPA4 and SLIT3 are the most negatively co-expressed genes with scarlet. CPA4, or carboxypeptidase A4, functions as a protease that hydrolyzes peptide bonds at the carboxy-terminal end of a protein or peptide. SLIT3, slit homolog 3 protein, is a developmental protein which aids in the differentiation and development of the nervous system.

## Discussion

CRISPR gene editing has been successfully implemented in a growing number of emerging model eukaryotic organisms (e.g., squid, ant, tick) using various effective delivery methods (Trible et al. 2017; Xu et al. 2019; Crawford et al. 2020; Sharma et al. 2022). These efforts have paved the way for future functional genomic studies to examine genotype-phenotype relationship in unprecedentedly diverse organisms. Among these emerging systems, *Daphnia* has excellent potential for genomic functional studies, largely thanks to the wealth of knowledge accumulated from decades of research on their evolution/adaptation, ecology, toxicology, phenotypic plasticity, and response to environmental factors (Altshuler et al. 2011). Since the development of first-generation *Daphnia* genomic tools (Colbourne et al. 2011), researchers have identified a large number of candidate genes responsible for various biological processes, such as adaptation to freshwater salinization (Wersebe and Weider 2023), heavy metal contamination (Shaw et al. 2007), the origin of obligate parthenogenesis and cyclical parthenogenesis (Xu et al. 2015; Xu et al. 2022; Huynh et al. 2023; Snyman and Xu 2023), and adaptation to ecologically distinct habitats (Ye et al. 2023), which are ready to be further interrogated for functional insights.

As a major tool for functional studies, although microinjection-based CRISPR-Cas9 gene editing for *Daphnia pulex* has been established (Hiruta et al. 2018), this study addresses several technical aspects of the microinjection procedure that have not been fully optimized. Furthermore, through creating *scarlet* gene knock-out mutants, we evaluate the efficiency of creating heritable mutations with CRISPR-Cas9/Cas12a, the spectrum of on-target mutations, and potential off-target mutations. Lastly, as *scarlet* appears to be pleiotropic, likely involved in the production of histamine and other neurotransmitters (Ismail et al. 2021), we examine the swimming behavior of *scarlet* mutants and the associated transcriptomic profiles to investigate the pleiotropic effects of *scarlet* and the underlying genetic causes.

### Effectiveness of the microinjection procedure

In this study we have developed a robust and effective procedure for generating knock-out mutants in *D. pulex*. The microinjection procedures in model organisms such as *Drosophila* (Ringrose 2009), *Caenorhabditis elegans* (Evans 2006), *Anopheles* mosquitoes (Carballar-Lejarazú et al. 2021) inspired us to develop optimized fabrication for injection needles with upgraded glass capillary (i.e., aluminosilicate) and repurpose a flipped small Petri dish as the injection stage for *Daphnia* embryos. We directly inject RNPs instead of plasmids encoding Cas9/Cas12 and guide RNAs because RNPs can result in mutations at a much greater efficiency than injecting plasmids or mRNA encoding Cas enzymes (Kim et al. 2014; Hendel et al. 2015). Most importantly, based on the literature of the development of asexual *Daphnia* embryos (Ojima 1958; Zaffagnini and Sabelli 1972; Hiruta et al. 2010), we propose and investigate that approximately 10 minutes post ovulation (1-2 min ovulation time, 6 min in ice-cold medium, and 2-3 min injection time), while the embryos are still in the single-cell stage, provides an effective time window for inducing heritable biallelic modifications.

The results of our CRISPR-Cas9 knock-out experiments at the *scarlet* locus strongly supported this idea, with a success rate of 1-15% in generating clear-eyed G_0_ individuals (**Table 1**). More importantly, all the G_0_ individuals from the Cas9 experiments carry heritable biallelic mutations, as evidenced by the clear-eye phenotype in all their offspring, highlighting the effectiveness of our microinjection strategy.

Consistent with the results of Cas9 knock-out experiments, our microinjection experiments using the A.s. Cas12a nuclease also efficiently generated heritable biallelic mutations. This is the first successful implementation of CRISPR-Cas12a in *Daphnia* to the best of our knowledge. The addition of Cas12a to the *Daphnia* gene editing toolkit significantly expands the range of editable genomic regions beyond what can be achieved with Cas9 nuclease alone.

However, it is notable that our Cas12a knock-out experiments yielded a large portion of clear-eyed (5 out of 8) mutants due to somatic mutations (i.e., no clear-eyed asexual progenies). We offer a potential explanation for the lower rate of introducing heritable mutations compared to the Cas9 knock-out experiments. The A.s. Cas12a nuclease used in this study is temperature dependent and has low activity level below 30°C (Moreno-Mateos et al. 2017). As 30°C and above is outside the normal temperature range of *Daphnia pulex*, in our experiments we kept the Cas12a-injected embryos at 33°C for only two hours post injection and then transferred them to 25°C. The Cas12a nuclease was likely not fully active in the single-cell stage embryo during the 2-hour incubation at 33°C. Thus, the editing activity most likely took place after the one cell stage and affected only somatic tissues including the eye. A potential solution to increase the chance of germline modification is to incubate the injected embryos at the (near) optimal temperature of Cas12a (e.g., 37°C), the detrimental effects of which to the embryos have to be experimentally determined for *Daphnia*. Moreover, as more temperature tolerant versions of Cas12a nuclease have become available (e.g., Cas12a Ultra from IDT, which was unavailable at the time of this study), it will be beneficial for future studies to examine its gene editing efficiency at *Daphnia*-appropriate or out-of-range temperatures.

### Mosaicism in the germline cells and implications

One of the most interesting findings of this study is the mosaicism in the germline cells of *scarlet* G_0_ mutants, which informs us of the editing process in the embryos and the inheritance of edited alleles across generations. In general, genetic mosaicism resulting from CRISPR-Cas gene editing in human and mouse zygotes has been recognized as a consequence of the prolonged activity of Cas nucleases beyond the first embryo cleavage event (Davies 2019). Although we intended to create biallelic modification in the one-cell stage of the *Daphnia* embryos, in some cases the successful editing only occurred after the one-cell stage, affecting different tissues through independent editing events and resulting in different Cas-induced mutations.

It is evident from both our Cas9 and Cas12 experiments that the germline cells of G_0_ individuals (with mosaic scarlet phenotype) were differentially edited, which asexually produced mixed broods of clear-eyed and black-eyed progenies (**Figure 3, Table S2 and S3**). Even among the clear-eyed progenies of the same mosaic G_0_s, our genomic analyses unveil that their knock-out *scarlet* genotypes are different (**Table S4**). These observations also suggest that during the asexual reproduction cycle of *Daphnia*, oogonia going into one asexual brood are derived from different primary oocytes, rather than one primary oocyte giving rise to all the oogonia.

Genetic mosaicism due to gene editing is generally considered a potential risk for clinical applications (Davies 2019) and could confound downstream analyses of the mutants. Our genomic sequencing of the knock-in mutants was an example of the confounding effect of germline mosaicism. Without realizing mosaicism and assuming all asexual progenies of a female *Daphnia* were genetically identical, we pooled all the progenies of KI mutants to establish a “clonal” mutant line. The presence of more than 2 alleles at the *scarlet* locus in the genomic sequences of this “clonal” line strongly indicates the presence of germline mosaicism in the G_0_ individual. Nonetheless, germline mosaicism most likely does not exist in the G_1_ individuals as all their offspring (G_2_s) exhibit the same phenotype as the mother. There is also no reason to believe that Cas nucleases could be present in the G_1_s due to transgenerational passing down from the G_0_s.

Despite its confounding effects, we argue that germline mosaicism due to the prolonged activities of Cas nucleases can be advantageous for *Daphnia* gene editing experiments. This is because, even in the absence of editing during the one-cell embryo stage, the prolonged nuclease activities can increase the chances of generating heritable biallelic alterations in the germline cells. Furthermore, given the independent editing activities in germline cells, multiple knock-out and knock-in genotypes could appear in the G_1_ offspring, facilitating the production of desirable mutant genotypes.

On the other hand, germline mosaicism necessitates a thoughtful plan for *Daphnia* mutant screening experiments, especially for those focusing on genes with no readily visible phenotypes. We suggest that a G_0_ mom should be maintained through at least two or three broods. The first-brood individuals of the same mom can be sacrificed for genotyping at the target locus to identify the presence/absence of mutant alleles. For the matter of efficiency, the first brood can be pooled for DNA extraction, PCR amplification of target locus, and a T7 endonuclease assay (Parkinson and Lilley 1997) used to detect induced mutations. If mutant alleles are detected from the first brood of a G_0_ individual, this G_0_ is either a mosaic or pure mutant. Then each of their second-brood progenies can be used to establish clonal lines that can be individually genotyped to identify mutant lines. We caution that even if the G_0_ mom carries germline mutations, the first brood could contain no mutant individuals, thus misleading our conclusions. To mitigate this, one could expand the first round of genotyping to the first two broods to increase the chance of detecting mutant alleles.

### Knock-out genotypes and implications

Despite their error-prone nature, nonhomologous end joining (NHEJ) and microhomology-mediated end joining (MMEJ) are primary pathways for the repair of DNA double-strand breaks when homologous repair template is not available (Sfeir and Symington 2015). In our Cas9 and Cas12a knock-out experiments, two gRNAs targeting the scarlet locus were co-injected. Therefore, we initially expected to see a homozygous segmental deletion if DNA double-strand breaks occur simultaneously at the two target locations. In the event of only one of the locations experienced cleavage, we expected to see small indels at one location.

However, the genotypes of our knock-out mutants show more complicated DNA repair outcomes, revealing some under-appreciated aspects of the gene editing process in *Daphnia* embryos. In fact, clean segmental deletions between two target sites only occurred in two out of 18 knock-out Cas9 mutants (KO2, KO4, **Table S4**), suggesting that Cas-induced cleavage at the two target locations occurs rarely at the same time, which could be due to their differential accessibility to the binding of RNPs associated with local nucleosome occupancy or other factors.

The majority of Cas9 mutants show small indels at both of the target locations. The absence of segmental deletions spanning the two target sites strongly suggests that the DNA cleavage at these sites did not occur at the same time, most likely one after another. Consistently, in our Cas12a mutant KO10, only target site 2 had mutations, whereas site 1 had no mutation, possibly due to reduced activity of Cas12a in our experiments. Furthermore, the induced modifications on the two alleles are different, suggesting independent repair events. This is supported by the genotype of our Cas12 mutants KO7 and KO8, where only one allele of either target site 1 or target site 2 was modified, clearly pointing to the possibility that the DNA cleavage on the two alleles do not occur at the same time.

Furthermore, we find large deletions (>1000bp) at target site 2 in a few Cas9 mutants. This is not an uncommon DNA repair outcome for Cas9-induced cleavage and has been previously reported in *C. elegans*, mouse zygote and cultured cells (Shin et al. 2017; Adikusuma et al. 2018; Au et al. 2019; Davies 2019). As to the repair mechanism generating these unintended large deletions, MMEJ DNA repair and polymerase theta-mediated end joining has been proposed as a candidate mechanism (Owens et al. 2019; Schimmel et al. 2023). Although these large deletions do not disrupt any other coding regions in our *scarlet* mutants, it is crucial to consider the potential occurrence of large deletions when designing CRISPR target sites and genotyping mutants. This type of large deletion is hardly detectable through regular PCR. For example, the genotype of our KO2 sample was initially identified as a homozygous segmental deletion based on a regular PCR test. However, its genomic sequencing unveiled a segmental deletion on only one allele, whereas the other allele harbors a large deletion upstream of target site 2 that removes one primer landing site for our PCR assay, which results in the PCR amplification of only one allele with segmental deletion as observed in our initial genotyping.

### Knock-in efficiency

Homology-dependent repair (HDR) with CRISPR using a DNA donor template is notoriously inefficient in generating high-fidelity DNA modification (Riesenberg et al. 2023; Schimmel et al. 2023). This is primarily because HDR is inefficient in comparison to NHEJ and MMEJ, which results in the activation of latter pathways for the imprecise repair of DNA damages (Riesenberg et al. 2023). Although in the knock-in experiments we chose ssDNA as the donor template, which boosts HDR efficiency compared to plasmid-based templates, the results of our knock-in experiment still show inefficient HDR repair. We did not observe any precise editing as expected, with most mutants not incorporating the stop codon cassette in the repair template. In the two mutants (KI4, KI6) where incorporation occurred, the insertion of the stop codon cassette is accompanied by a large 635-bp deletion and other complex rearrangements at the target site. It is not clear which DNA repair pathway(s) are responsible for this mutagenic outcome. However, NHEJ-mediated knock-in (Maresca et al. 2013; Auer et al. 2014) seems capable of producing this kind of knock-in involving complex local rearrangement as seen in another *Daphnia* species *D. magna* (Kumagai et al. 2017).

Numerous strategies for improving the efficiency of HDR repair have been developed, including inhibiting key proteins of the NHEJ and MMEJ pathways (Riesenberg et al. 2023; Schimmel et al. 2023) and keeping a close proximity of the HDR repair template with the Cas components (Aird et al. 2018; Sharon et al. 2018). We note that the microinjection of small-molecule chemical inhibitors of NHEJ and MMEJ pathway (Schimmel et al. 2023) with RNP and HDR template into *Daphnia* embryos is worth further investigation because of its simplicity in implementation.

### Off-target mutations in scarlet mutants

Unintended mutagenesis caused by Cas nucleases at non-target genomic locations undermines the integrity of gene knock-out mutants. Our analyses of base-substitution and structural variation (SV) mutations in the *scarlet* mutant lines did not show excessive mutations in comparison to the wildtype, suggesting minimal risks of off-target mutagenesis of Cas9 and Cas12a in our experiments. Nonetheless, off-target risks need to be assessed in a case-by-case manner because multiple factors such as the uniqueness of target sites, nuclease concentrations, and the Cas nuclease variants used in specific experiments could jointly affect the occurrence of off-target mutations (Davies 2019). Before engaging in an extensive gene editing experiment, it is now possible to gain an empirical understanding of the off-target effects and genomic location of mistargets through *in vivo* or *in vitro* methods that combine the digestion of DNA by RNPs and high-throughput sequencing (Huang and Huang 2023)

### Daphnia as an emerging model for neurodegenerative behavior

Our behavioral assay of *scarlet* mutants shows the progression of spinning moves as individual daphniids grow up. The *scarlet* mutants in *D. magna* also show a similar progression pattern (Ismail et al. 2021). Interestingly, the spinning moves can be rescued by supplementing histamine to mutant neonates of *D. magna*, whereas adults’ spins are irreversible, suggesting a progressive neurodegenerative effect of the scarlet knock out mutation (Ismail et al. 2021). However, the rescuing effect of histamine still needs to be verified in *D. pulex*.

The progressive nature of the altered swimming behavior in *scarlet* mutants draws an interesting parallel with the worsening of symptoms in some human neurodegenerative diseases such as Amyotrophic Lateral Sclerosis (ALS) (Akcimen et al. 2023). Interestingly, the perturbed transcriptomes of our scarlet mutants offer insights into the potential mechanistic basis of this behavior. Several genes involved in human neurodegenerative diseases such as NMDA1, CNTNAP2, and ABAT are highly differentially expressed in the behavior-changing *scarlet* mutants compared to the wildtype.

These findings provide insight into the pleiotropic effects of the *scarlet* gene and open opportunities for further understanding the altered gene expression of critical disease-causing genes in relation to symptom progression. Nonetheless, our transcriptomic data is restricted to 2-3-day-old female neonates, leaving much to be explored regarding male’s behavioral and transcriptomic responses. As *Daphnia* is nearly transparent with the nervous system easily visible in the head region with a modern-day microscope, future studies can use single-cell RNA-seq or spatial transcriptomics to obtain precise neuron-specific transcriptomic profiles and can *in vivo* tag and track specific proteins across developmental stages in the context of disease progression. Moreover, the asexual clonal reproduction of *Daphnia* can provide an endless supply of experimental replicates of the same genetic background. Therefore, we suggest that the *Daphnia scarlet* mutants provide a powerful model system for understanding the genetic causes of neurological defects and associated behavioral aberrations from the perspective of faulty ABC transporter genes.

## Acknowledgments

We would like to thank J. Dittmer and P. Wright for their assistance with swimming behavior experiments. We also thank Michael Pfrender for his comments on an early draft of this manuscript. This work is supported by NIH grant R35GM133730, NSF CAREER grant MCB-2042490/2348390, NSF EDGE grant 2220695/2324639 to SX.

## Notes

### Competing Interest Statement

The authors have declared no competing interest.

