## Supplementary file 1 for "Efficient CRISPR genome editing and integrative genomic analyses reveal the mosaicism of Cas-induced mutations and pleiotropic effects of *scarlet* gene in an emerging model system"

# KO1

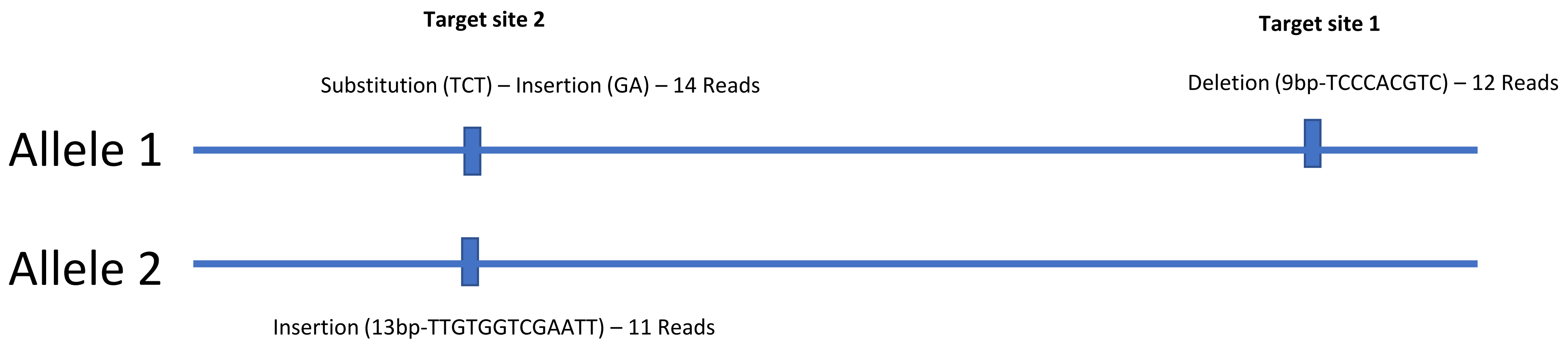
Target site 2
Target site 1
Substitution (TCT) – Insertion (GA) – 14 Reads
Deletion (9bp-TCCCACGTC) – 12 Reads
Allele 1
Allele 2
Insertion (13bp-TTGTGGTCGAATT) – 11 Reads

# KO1

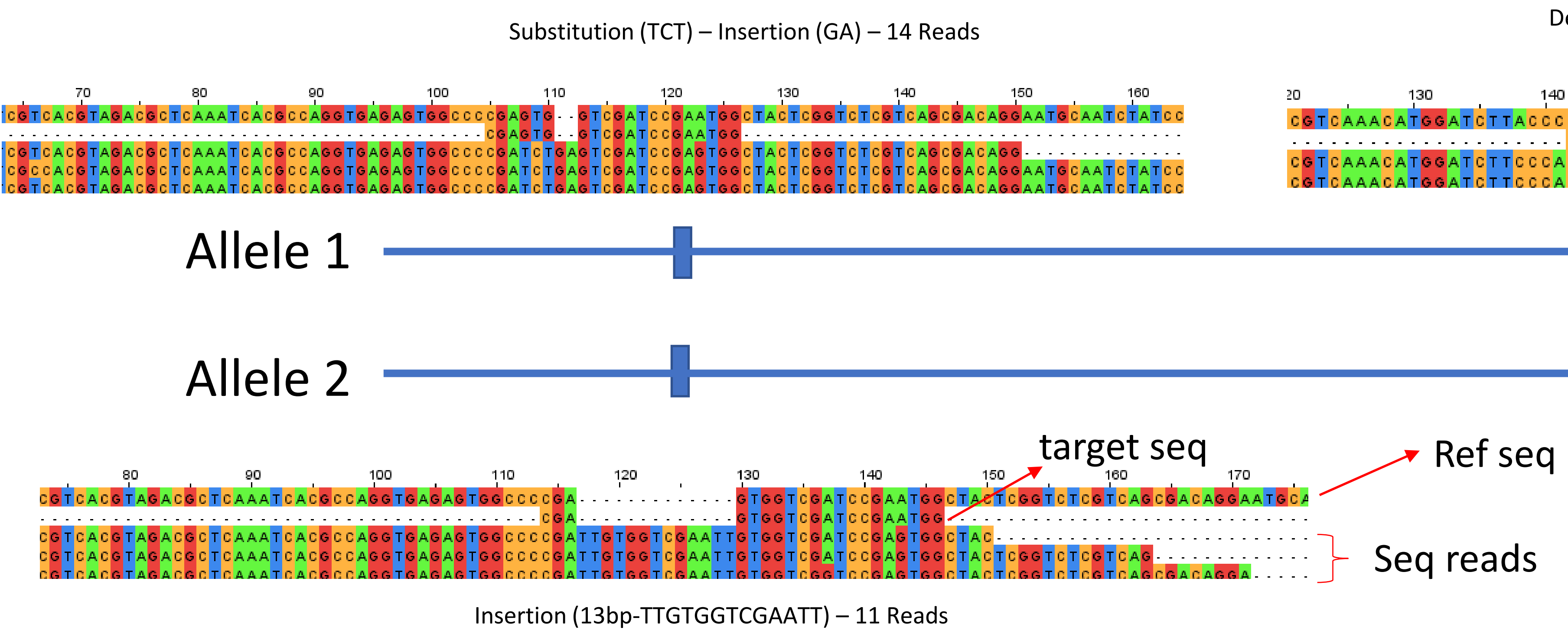


Note: In each alignment the top sequence is the reference sequence, the second top sequence is the target site sequence, and remaining sequences represent sequencing reads

# KO2

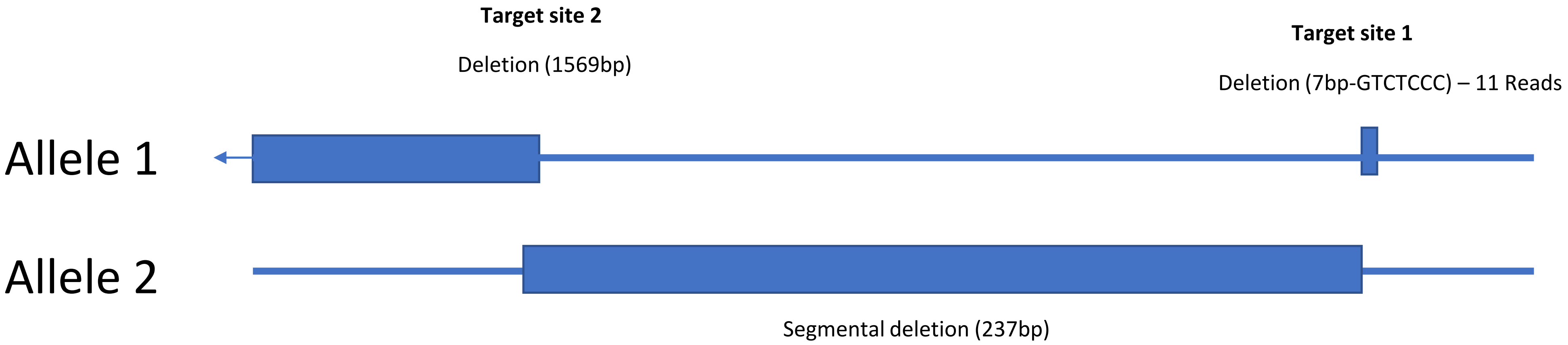
Target site 2
Deletion (1569bp)
Target site 1
Deletion (7bp-GTCTCCC) – 11 Reads
Allele 1
Allele 2
Segmental deletion (237bp)

# KO2

**Target site 1**

Deletion (7bp-GTCTCCC) – 11 Reads

**Target site 2**

1569 bp deletion supported by 8 split reads

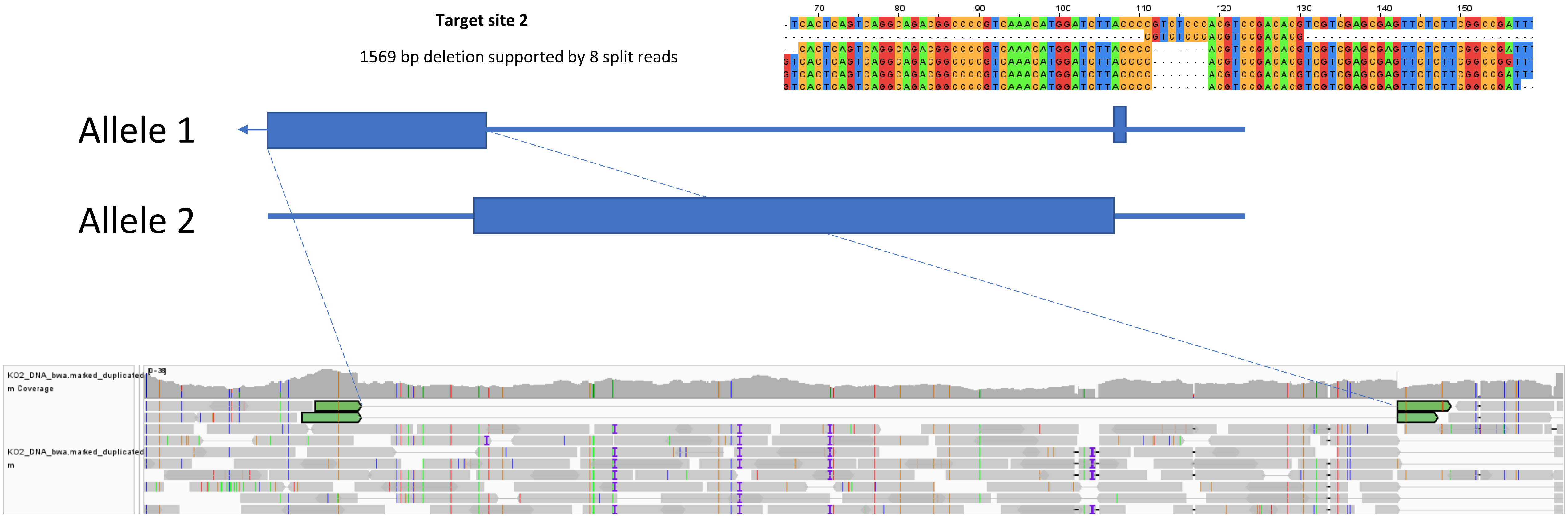

# KO2

**Target site 2**

Deletion (1569bp)

**Target site 1**

Deletion (7bp-GTCTCCC) – 11 Reads

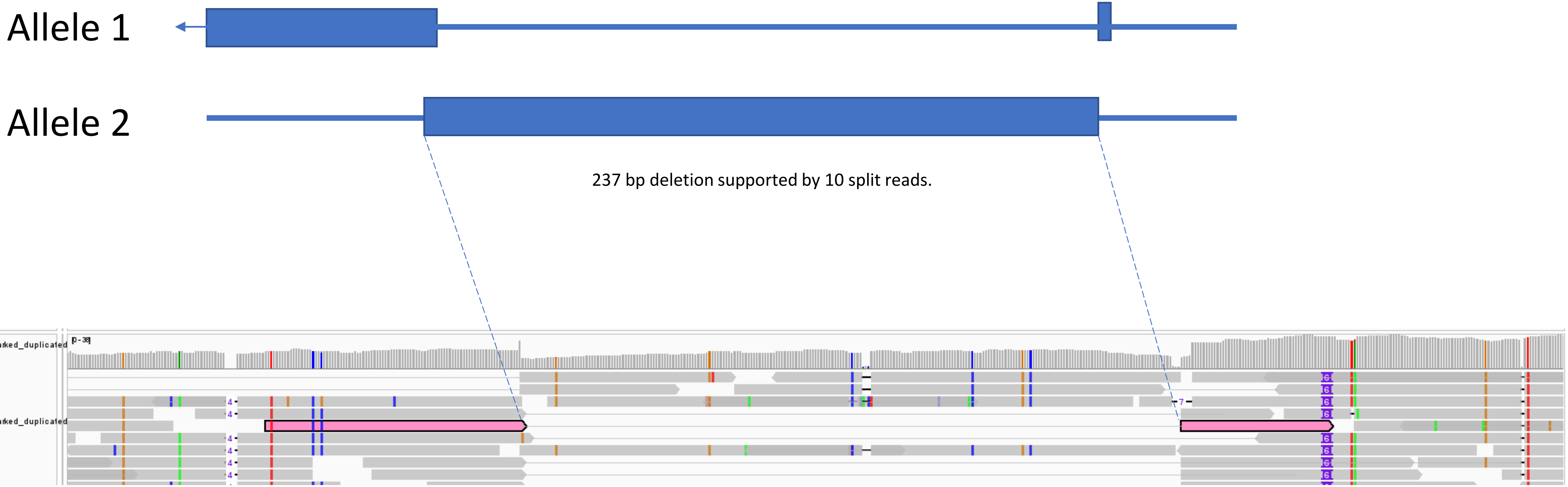

# KO3

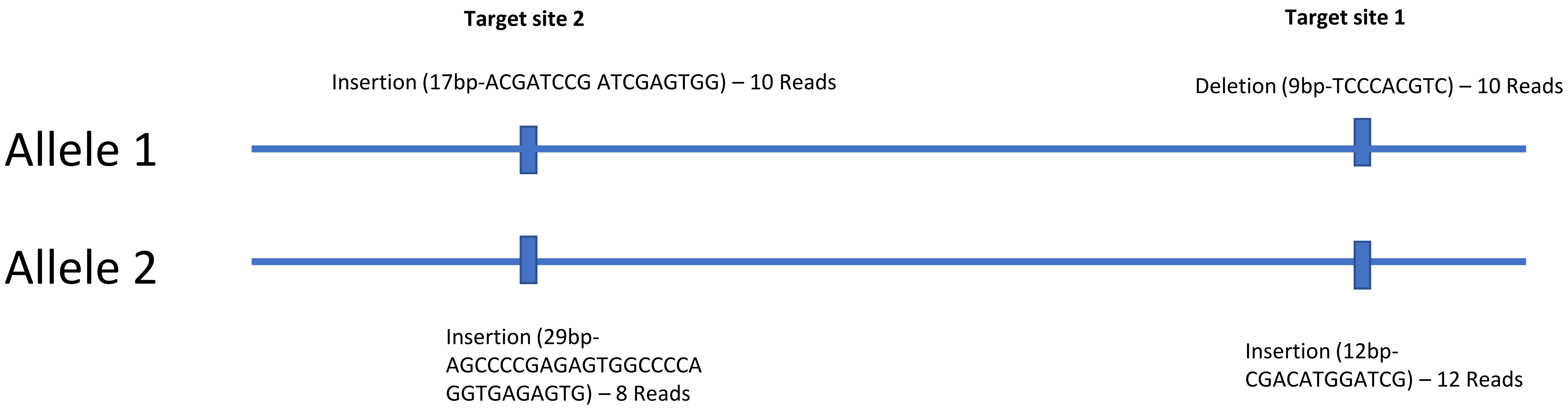
Target site 2
Target site 1
Insertion (17bp-ACGATCCG ATCGAGTGG) – 10 Reads
Deletion (9bp-TCCCACGTC) – 10 Reads
Allele 1
Allele 2
Insertion (29bp-AGCCCCGAGAGTGGCCCCA GGTGAGAGTG) – 8 Reads
Insertion (12bp-CGACATGGATCG) – 12 Reads

# KO3

**Target site 2**

Insertion (17bp-ACGATCCG ATCGAGTGG) – 10 Reads

**Target site 1**

Deletion (9bp-TCCCACGTC) – 10 Reads

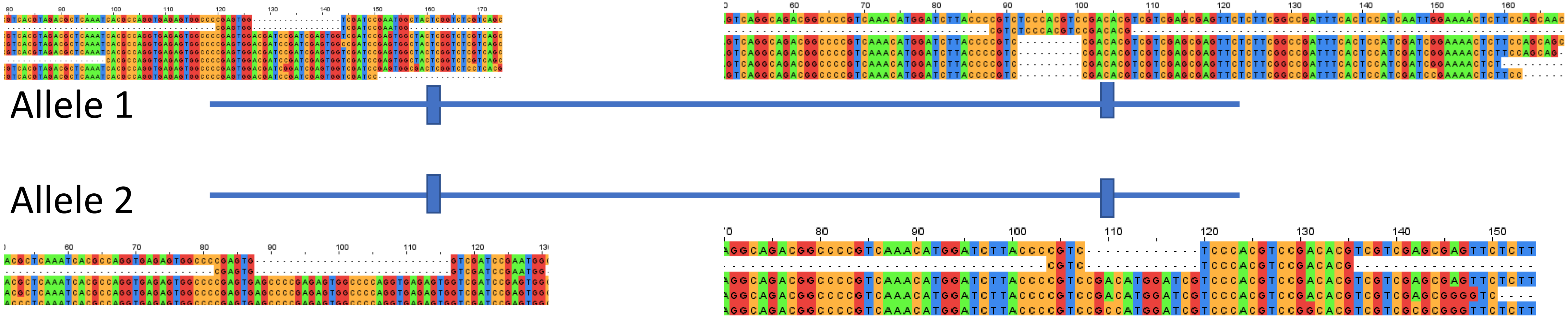


Insertion (29bp-AGCCCCGAGAGTGGCCCCA GGTGAGAGTG) – 8 Reads

Insertion (12bp-CGACATGGATCG) – 12 Reads

# KO4

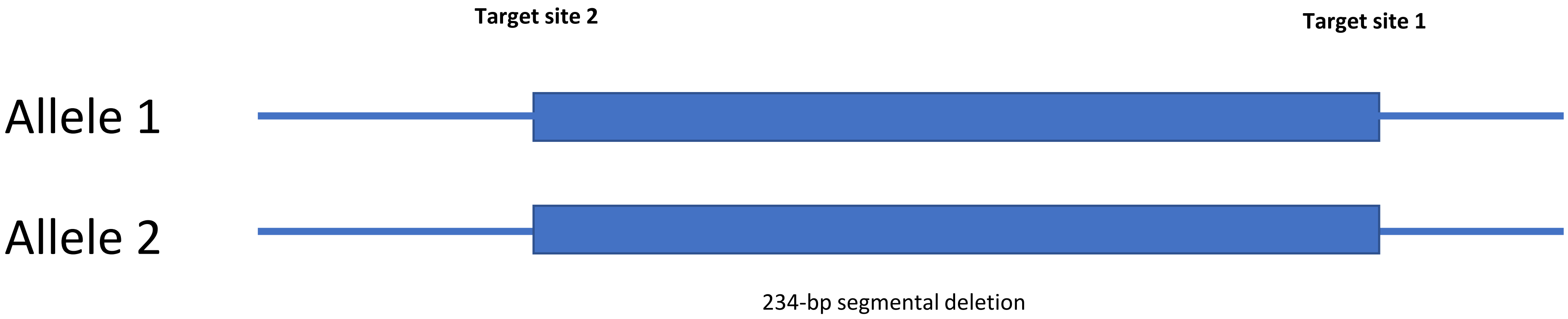
Target site 2
Target site 1
Allele 1
Allele 2
234-bp segmental deletion

# KO4

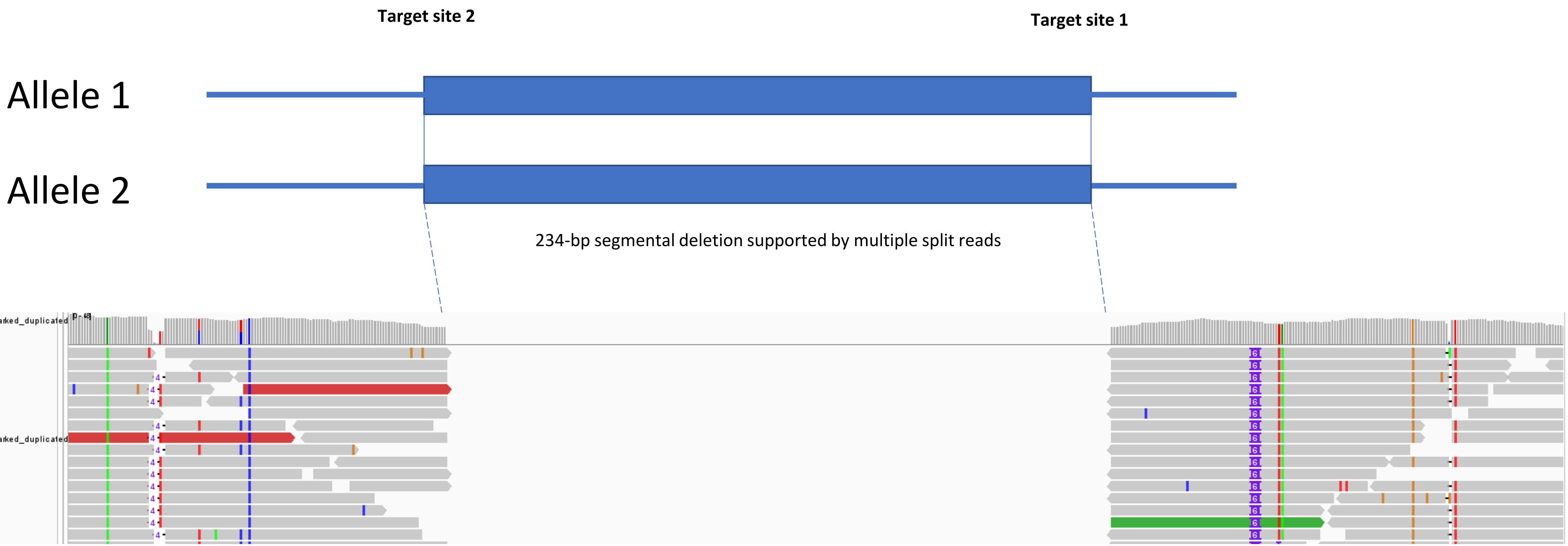
Target site 2
Target site 1
Allele 1
Allele 2
234-bp segmental deletion supported by multiple split reads
KO4_DNA_bwa.marked_duplicated
m Coverage
KO4_DNA_bwa.marked_duplicated
m

### KO5.1 broods

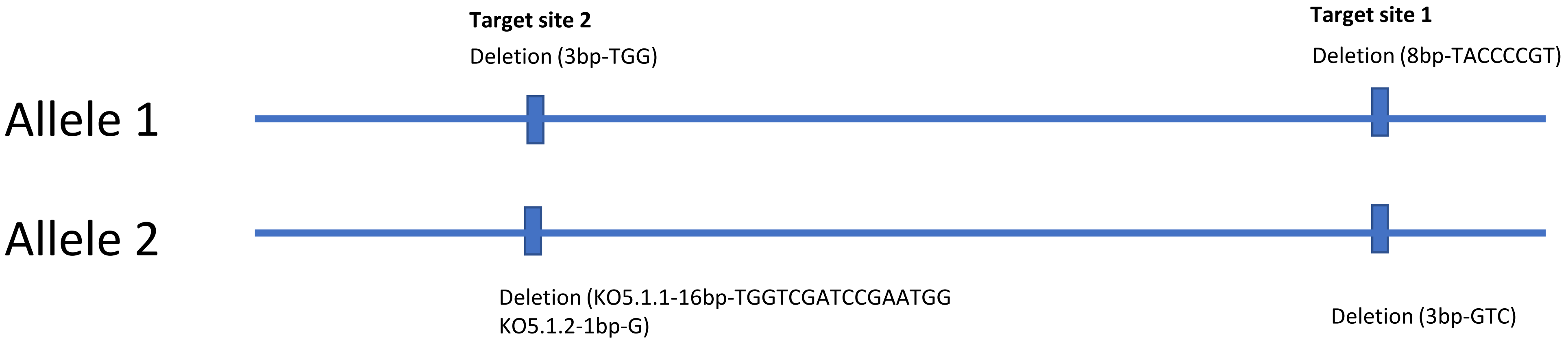
Target site 2
Deletion (3bp-TGG)
Target site 1
Deletion (8bp-TACCCCGT)
Allele 1
Allele 2
Deletion (KO5.1.1-16bp-TGGTCGATCCGAATGG
KO5.1.2-1bp-G)
Deletion (3bp-GTC)

# KO5.1.1

Deletion (3bp-TGG)-5 reads

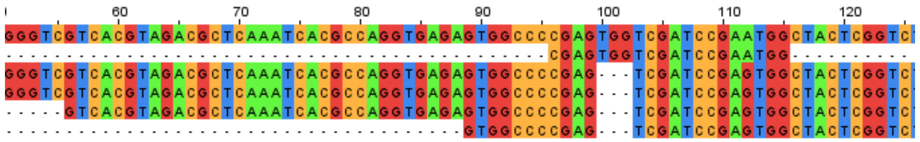

Deletion (8bp--TACCCCGT)-9 reads

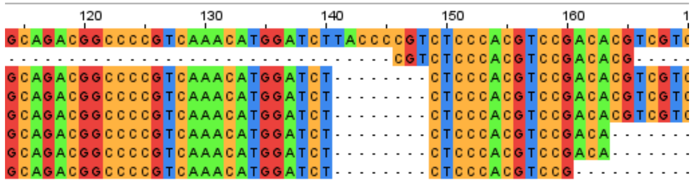

**Target site 2**

**Target site 1**

Allele 1

Allele 2

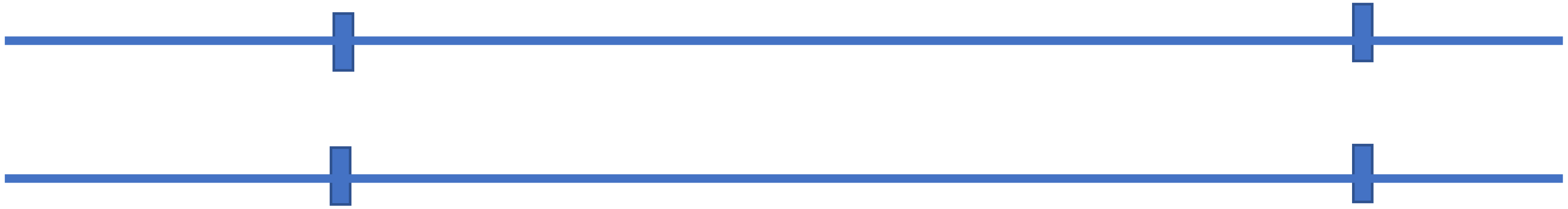

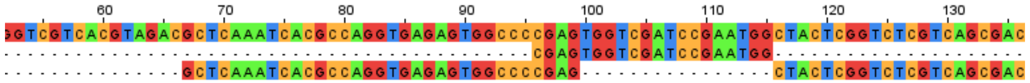

Deletion (16bp-TGGTCGATCCGAATGG)-1 read

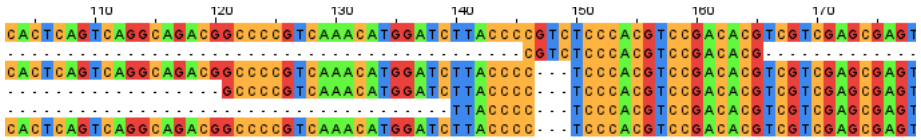

Deletion (3bp-GTC)-6 reads

# KO5.1.2

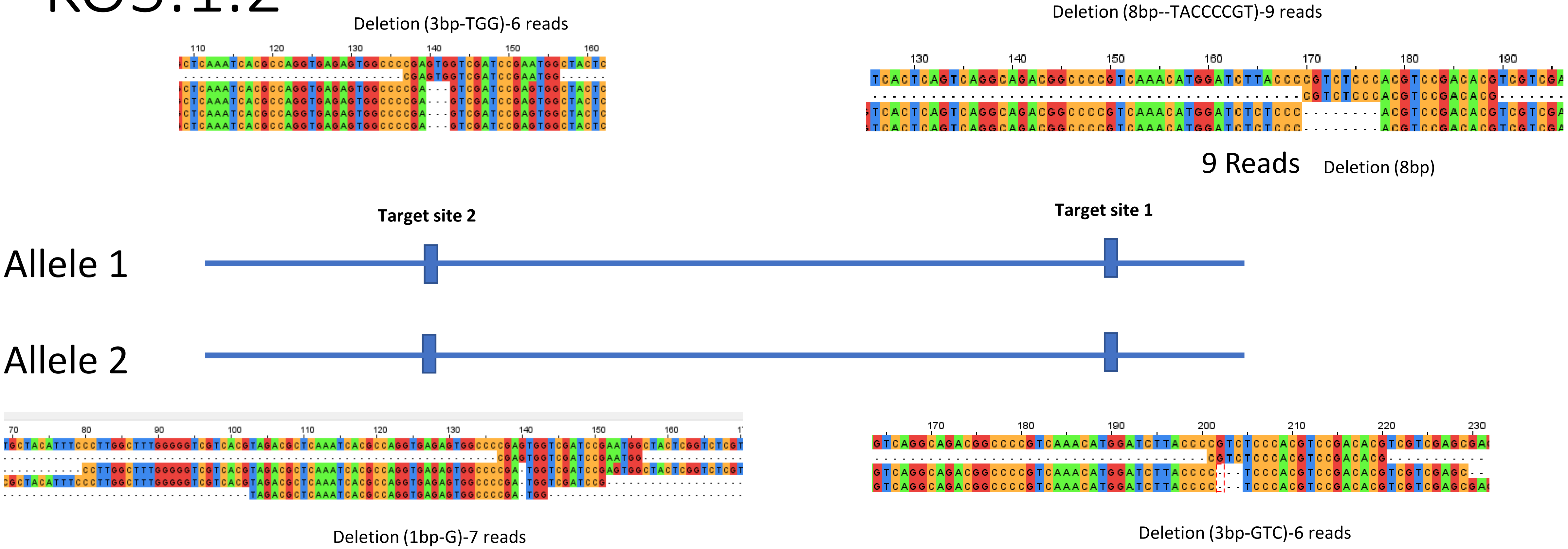
Deletion (3bp-TGG)-6 reads
Deletion (8bp--TACCCCGT)-9 reads
9 Reads
Deletion (8bp)
Target site 2
Target site 1
Allele 1
Allele 2
Deletion (1bp-G)-7 reads
Deletion (3bp-GTC)-6 reads

### KO6.1 broods

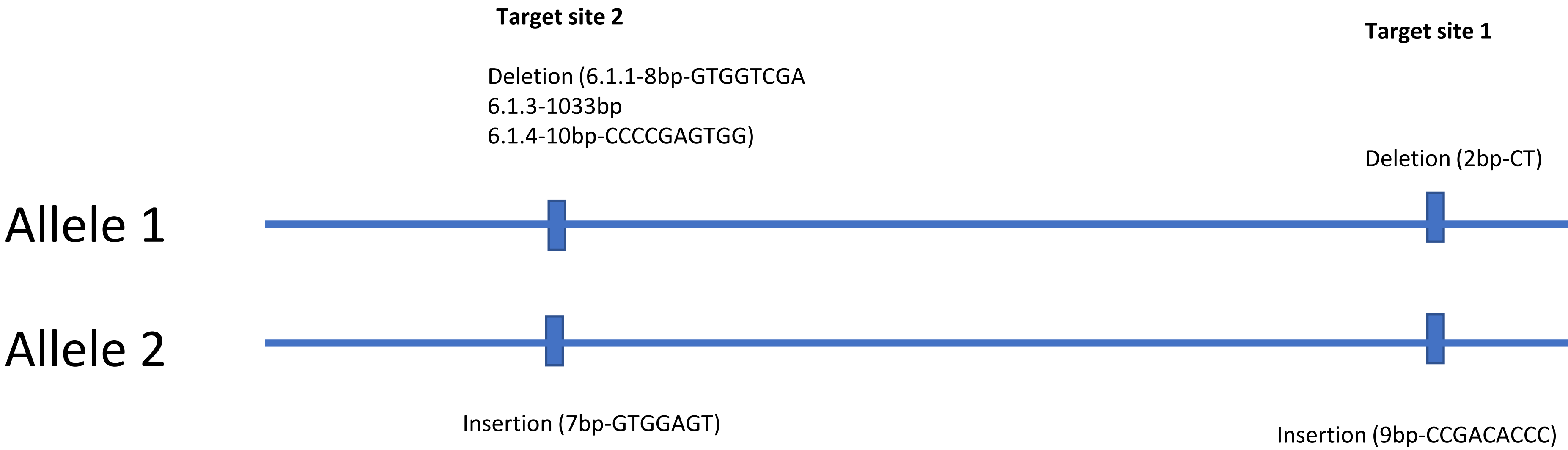
Target site 2
Deletion (6.1.1-8bp-GTGGTCGA
6.1.3-1033bp
6.1.4-10bp-CCCCGAGTGG)
Target site 1
Deletion (2bp-CT)
Allele 1
Allele 2
Insertion (7bp-GTGGAGT)
Insertion (9bp-CCGACACCC)

# KO6.1.1

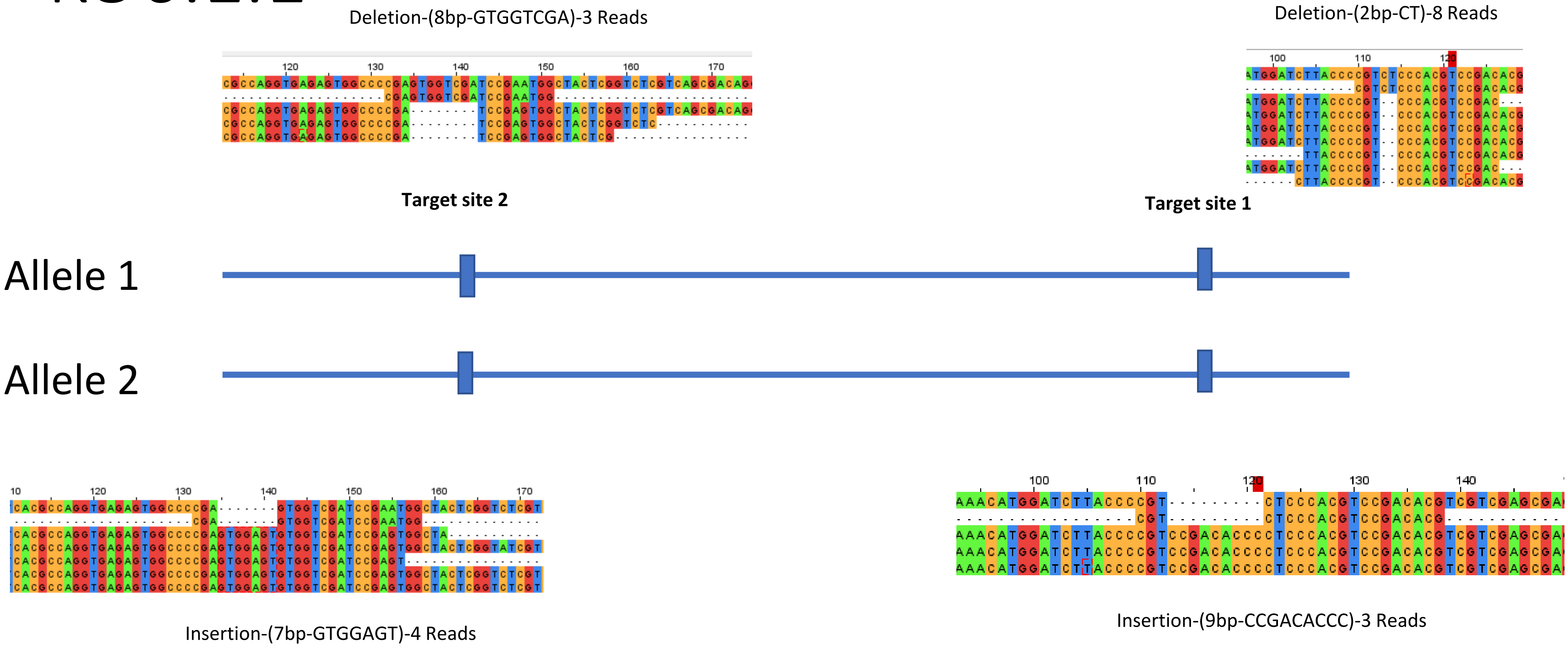

Deletion-(8bp-GTGGTCGA)-3 Reads
Deletion-(2bp-CT)-8 Reads
Target site 2
Target site 1
Allele 1
Allele 2
Insertion-(7bp-GTGGAGT)-4 Reads
Insertion-(9bp-CCGACACCC)-3 Reads

# KO6.1.3

deletion-(1033bp)-4 Reads

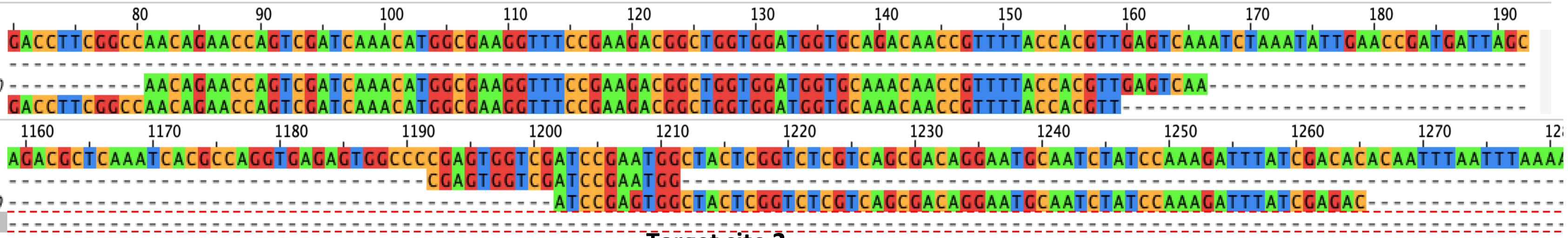

Target site 2

Deletion-(2bp-CT)-11Reads

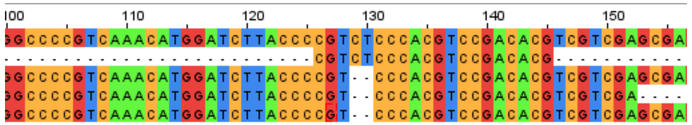

Target site 1

Allele 1

Allele 2

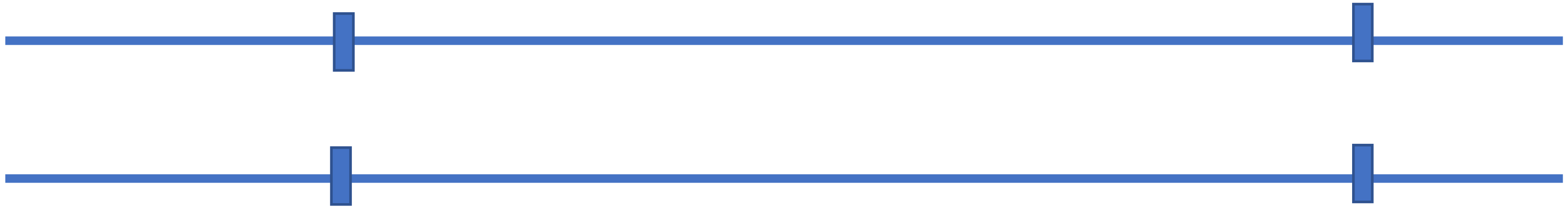

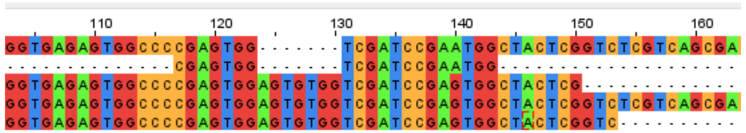

Insertion-(7bp-AGTGTGG)-4 Reads

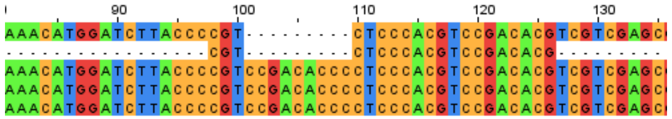

Insertion- (9bp-CCGACACCC) -5Reads

# KO6.1.4

Deletion-(10bp-CCCCGAGTGG)-3Reads

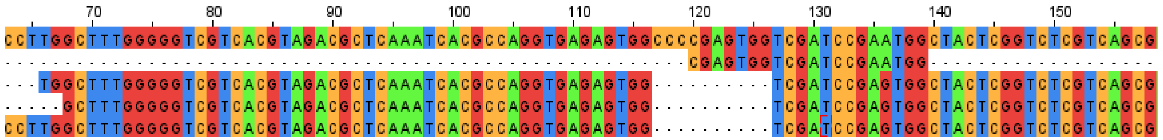

Deletion-(2bp-CT)-6Reads

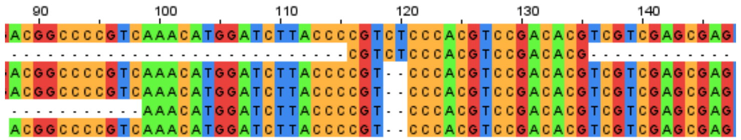

**Target site 2**

**Target site 1**

Allele 1

Allele 2

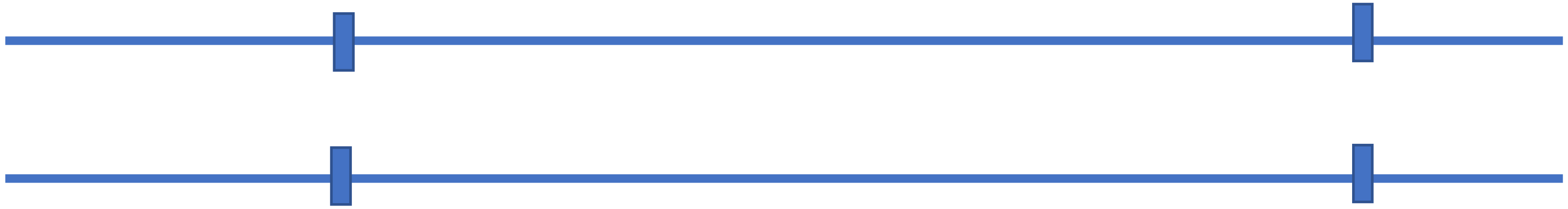

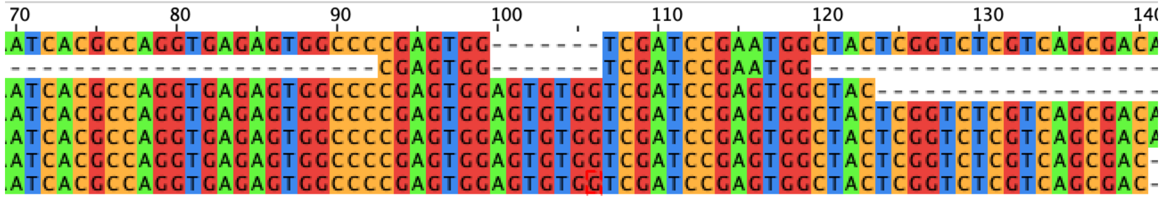

Insertion-(7bp-AGTGTGG)-11Reads

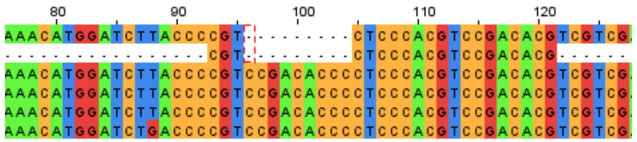

Insertion-(9bp-CCGACACCC)-6Reads

# KO6.2

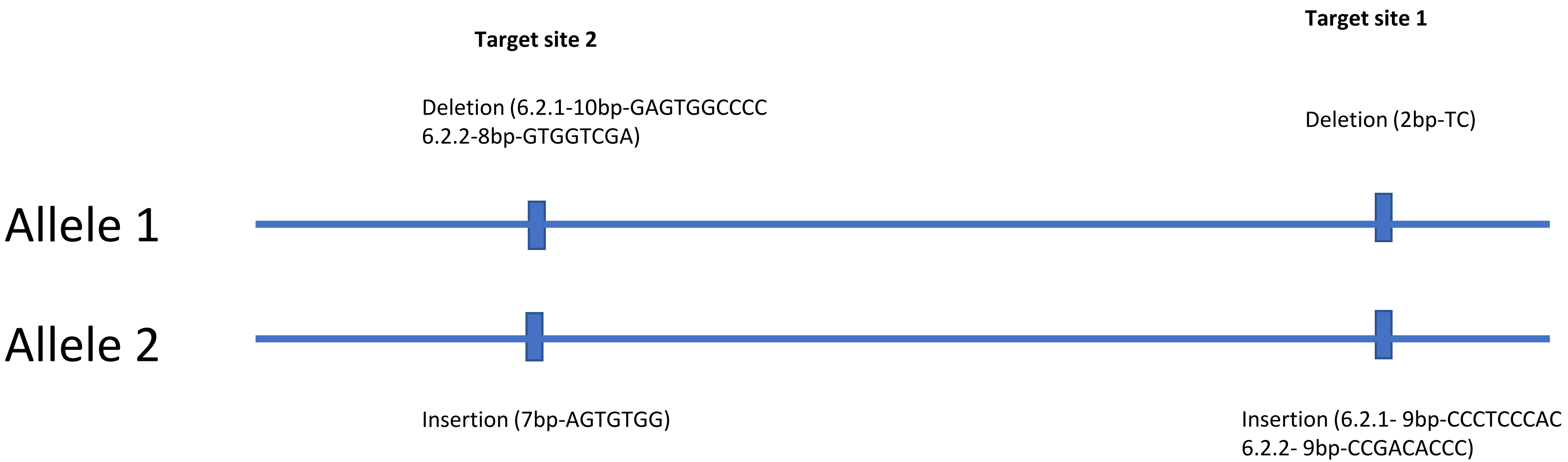
Target site 2
Target site 1
Deletion (6.2.1-10bp-GAGTGGCCCC
6.2.2-8bp-GTGGTCGA)
Deletion (2bp-TC)
Allele 1
Allele 2
Insertion (7bp-AGTGTGG)
Insertion (6.2.1- 9bp-CCCTCCCAC
6.2.2- 9bp-CCGACACCC)

# KO6.2.1

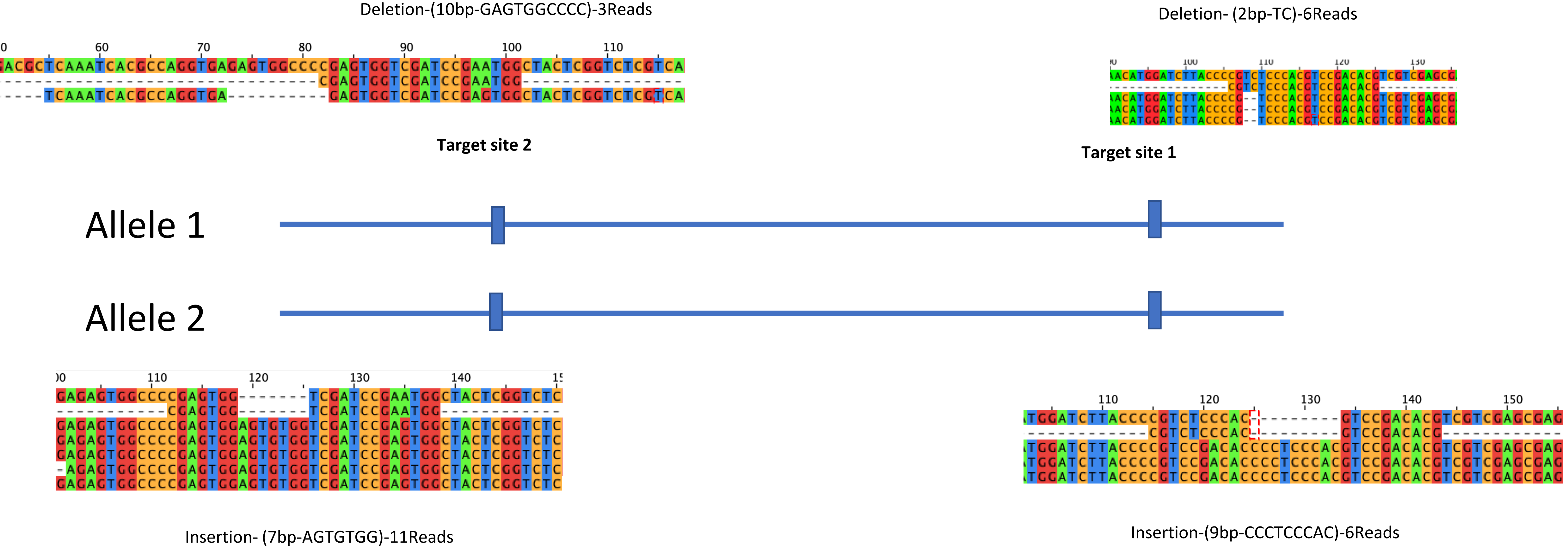

Deletion-(10bp-GAGTGGCCCC)-3Reads
Deletion- (2bp-TC)-6Reads
Target site 2
Target site 1
Allele 1
Allele 2
Insertion- (7bp-AGTGTGG)-11Reads
Insertion-(9bp-CCCTCCCAC)-6Reads

# KO6.2.2

Deletion-(8bp-GTGGTCGA)-4Reads

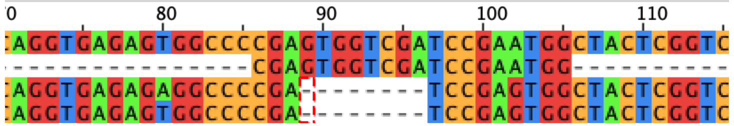

**Target site 2**

Deletion-(2bp-TC)-8Reads

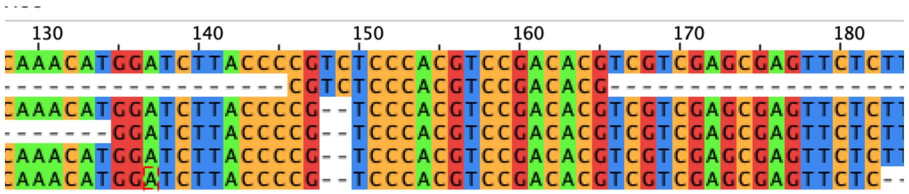

**Target site 1**

Allele 1

Allele 2

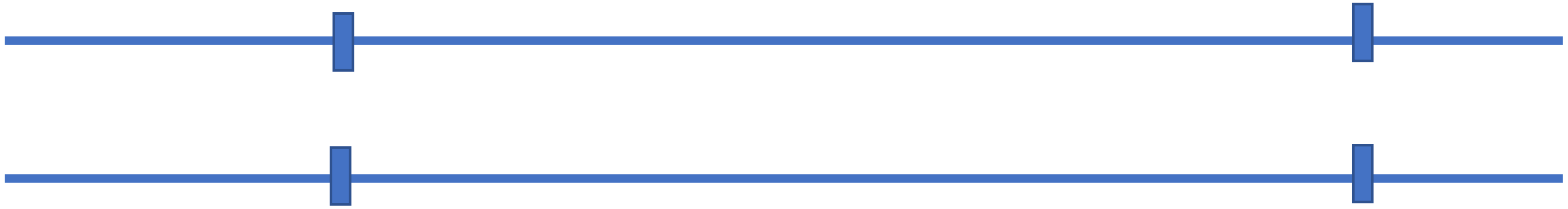

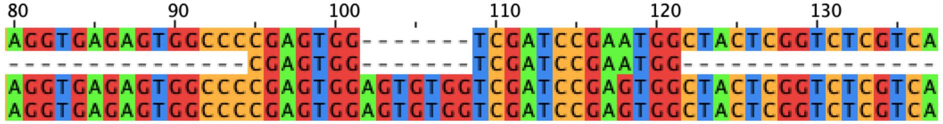

Insertion- (7bp-AGTGTGG)- 4Reads

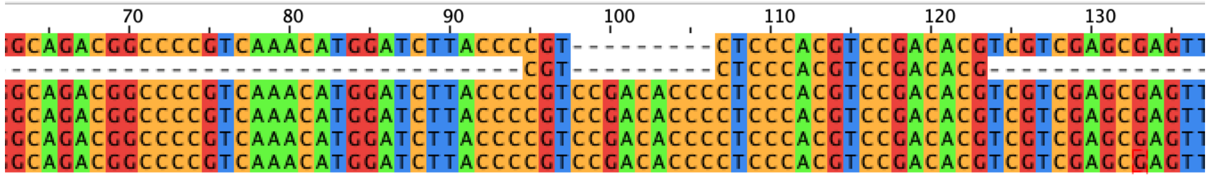

Insertion-(9bp-CCGACACCC)-4Reads

### KO6.3 broods

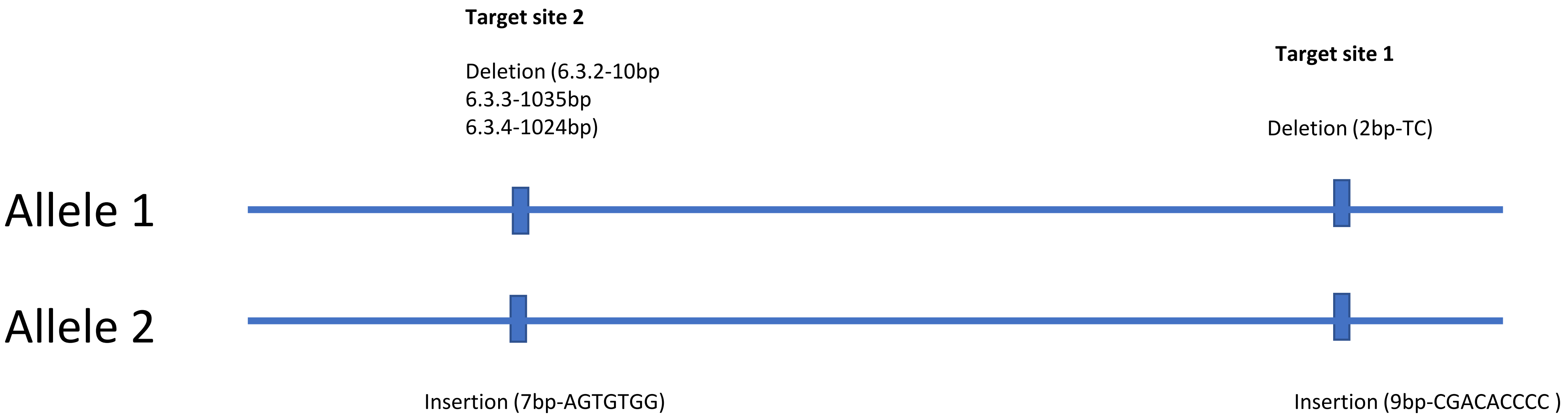
Target site 2
Deletion (6.3.2-10bp
6.3.3-1035bp
6.3.4-1024bp)
Target site 1
Deletion (2bp-TC)
Allele 1
Allele 2
Insertion (7bp-AGTGTGG)
Insertion (9bp-CGACACCCC )

# KO6.3.2

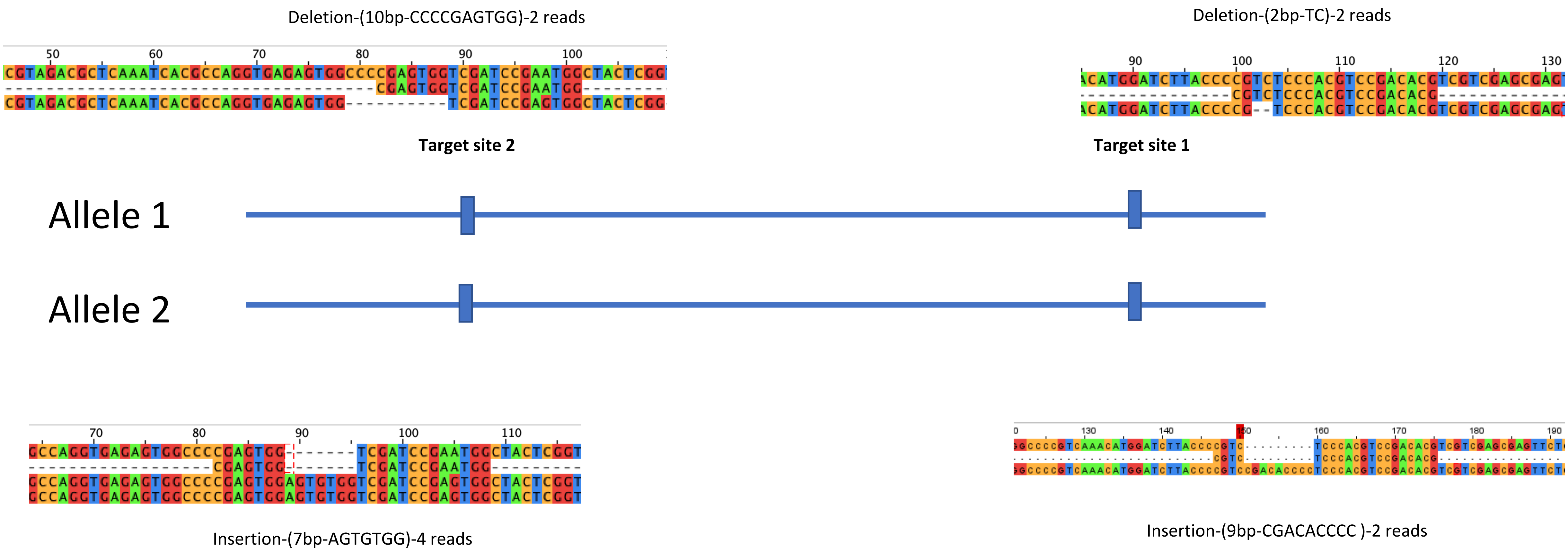
Deletion-(10bp-CCCCGAGTGG)-2 reads
Deletion-(2bp-TC)-2 reads
Target site 2
Target site 1
Allele 1
Allele 2
Insertion-(7bp-AGTGTGG)-4 reads
Insertion-(9bp-CGACACCCC )-2 reads

# KO6.3.3

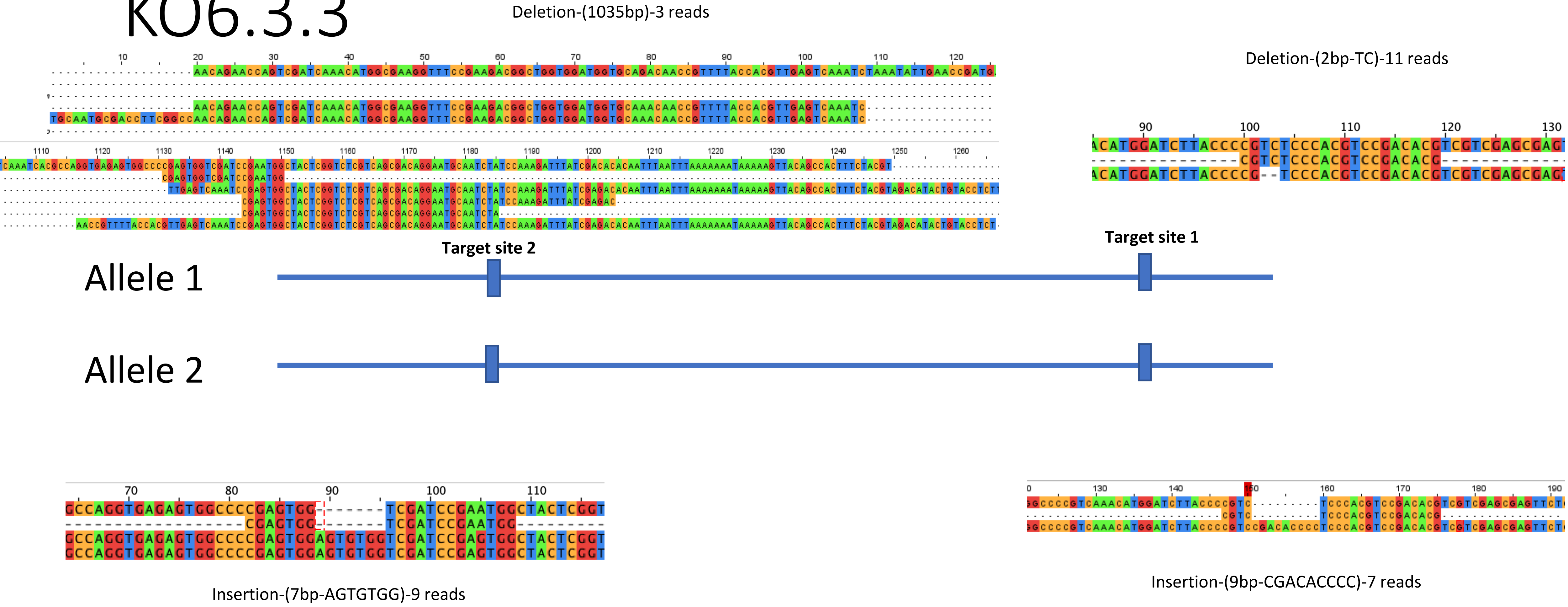

Deletion-(1035bp)-3 reads
Deletion-(2bp-TC)-11 reads
Target site 2
Target site 1
Allele 1
Allele 2
Insertion-(7bp-AGTGTGG)-9 reads
Insertion-(9bp-CGACACCCC)-7 reads

# KO6.3.3

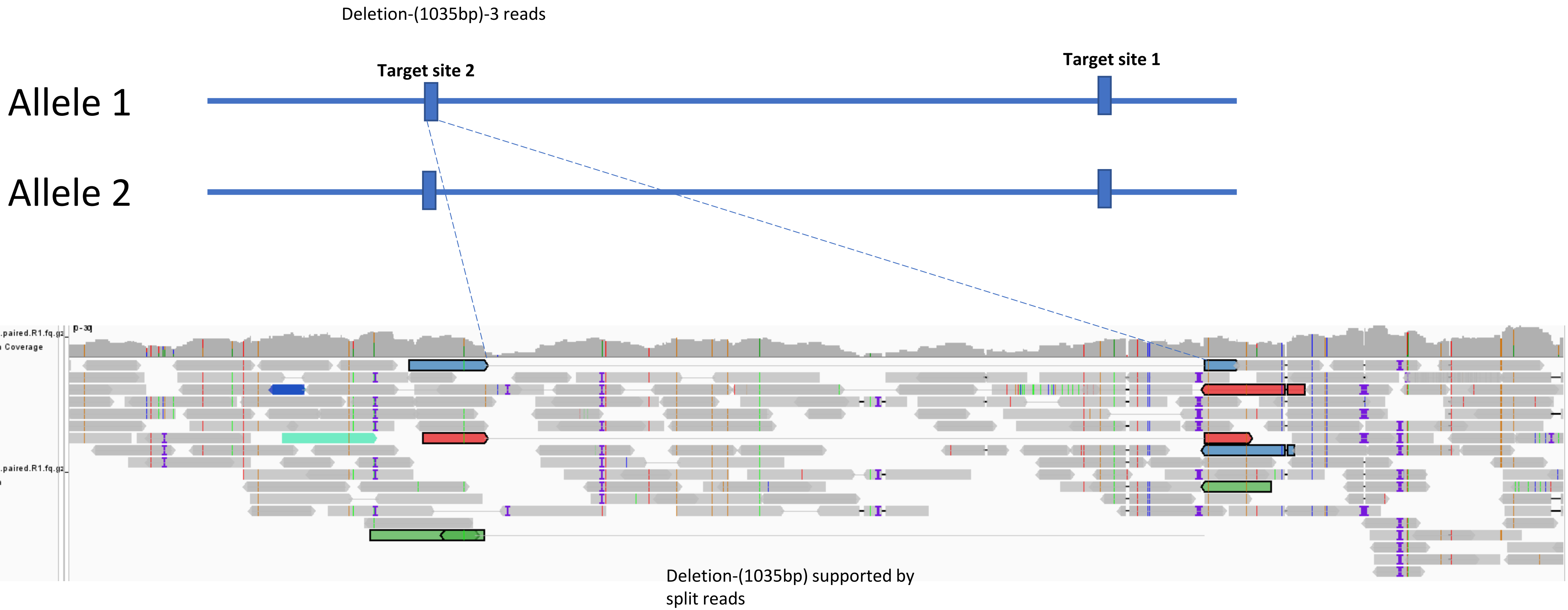
Deletion-(1035bp)-3 reads
Target site 2
Target site 1
Allele 1
Allele 2
KO6_3_3.trimmed.paired.R1.fq.gz_sorted.marked.bam Coverage
KO6_3_3.trimmed.paired.R1.fq.gz_sorted.marked.bam
Deletion-(1035bp) supported by split reads

# KO6.3.4

Deletion-(1024bp)-2 reads

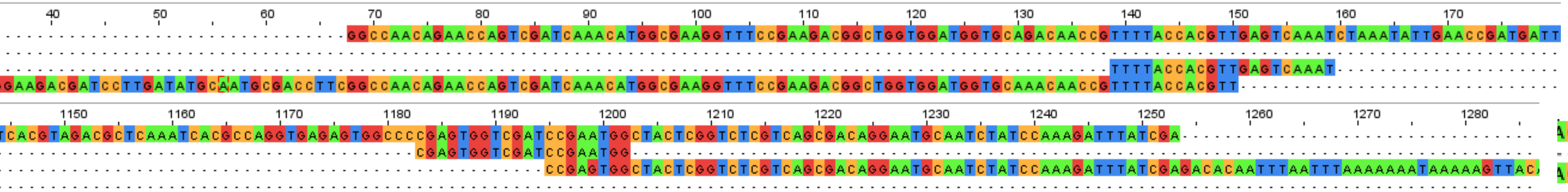

Deletion-(2bp-TC)-5 reads

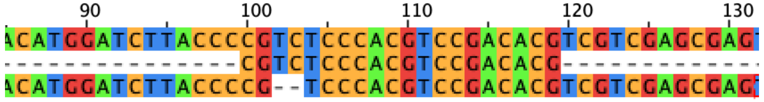

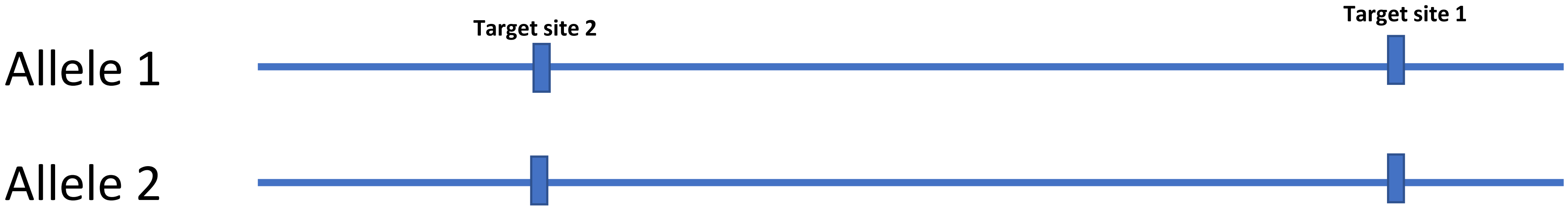


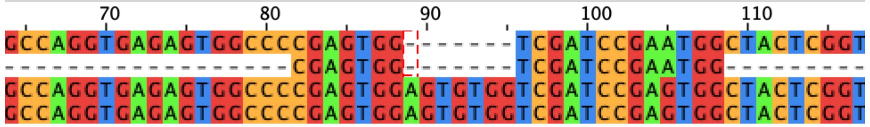

Insertion-(7bp-AGTGTGG)-4 reads

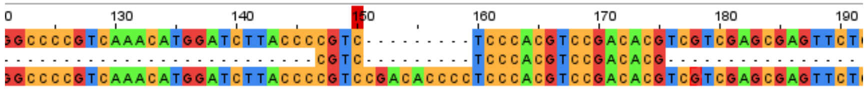

Insertion- (9bp-CGACACCCC)- 1 read

# KO6.3.4

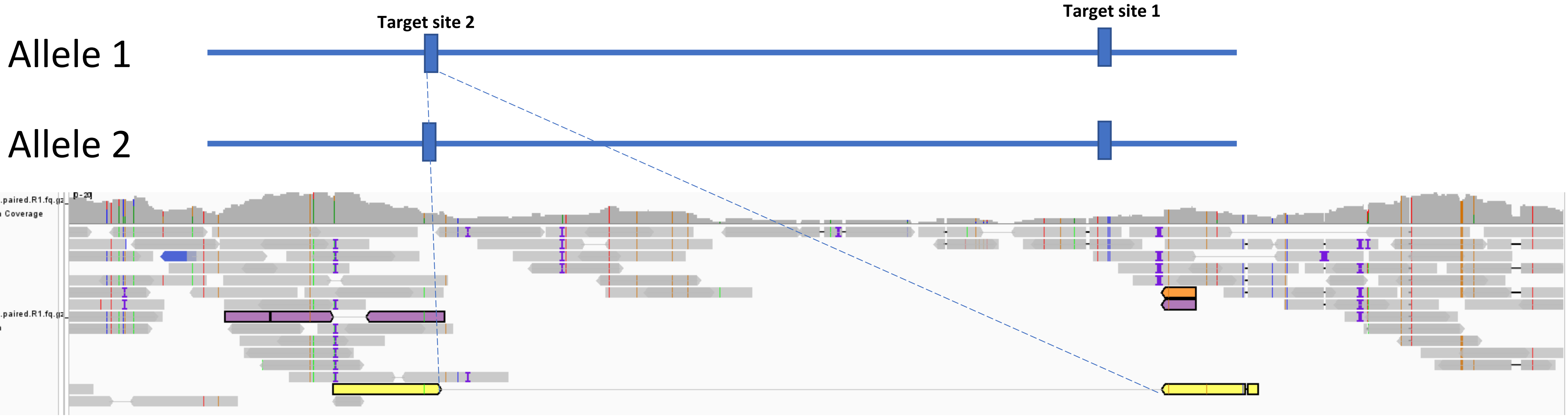


Deletion-1024bp supported by split reads

# KO6.4

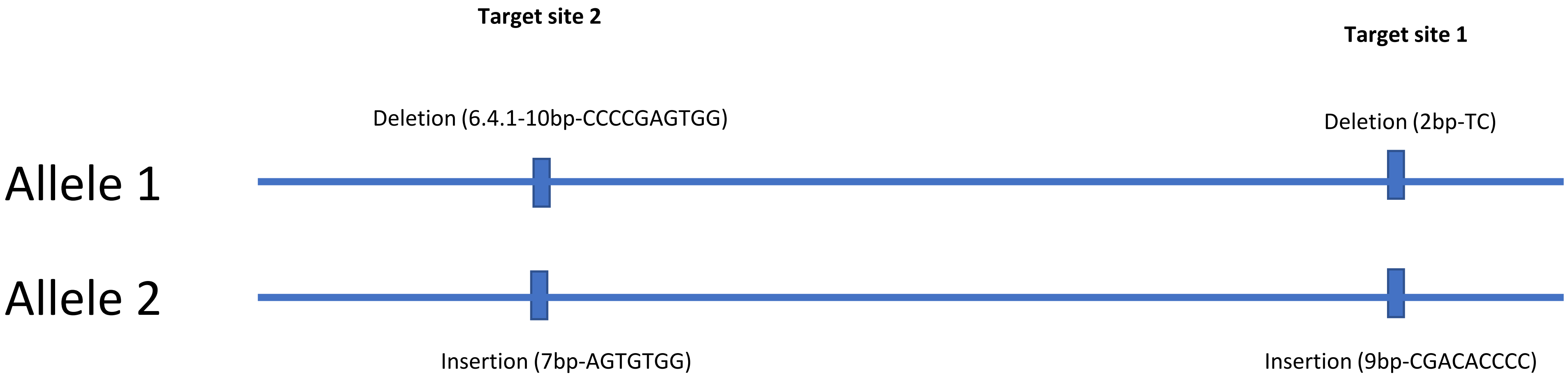
Target site 2
Target site 1
Deletion (6.4.1-10bp-CCCCGAGTGG)
Deletion (2bp-TC)
Allele 1
Allele 2
Insertion (7bp-AGTGTGG)
Insertion (9bp-CGACACCCC)

# KO6.4.1

Deletion-(10bp-CCCCGAGTGG)-4 reads

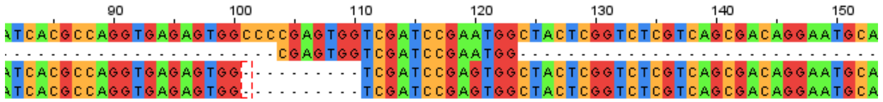

Deletion- (2bp-TC) -23 reads

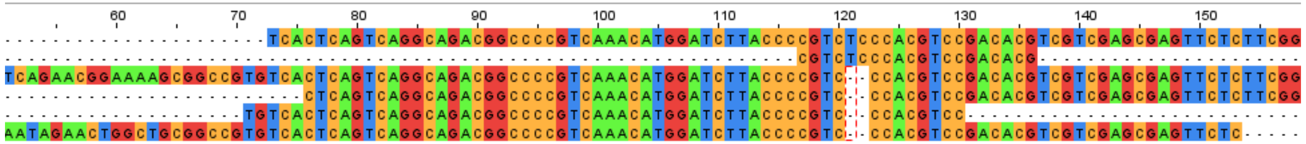

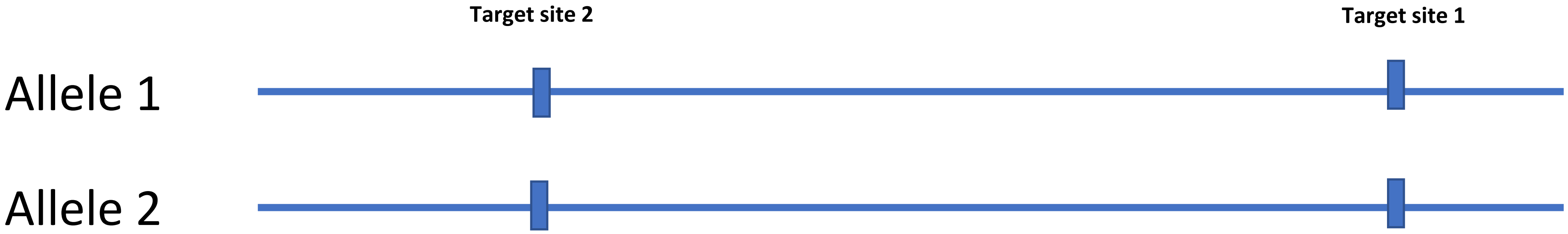


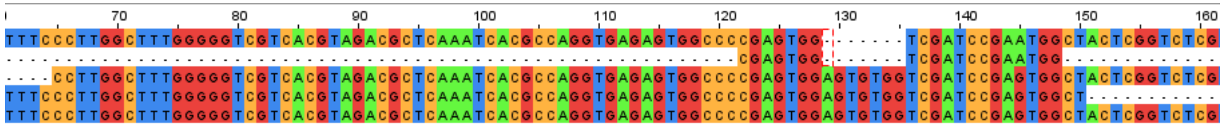

Insertion-(7bp-AGTGTGG)-12 reads

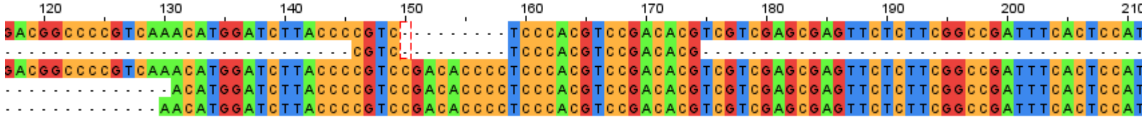

Insertion- (9bp-CGACACCCC)- 8 reads

# KO6.5

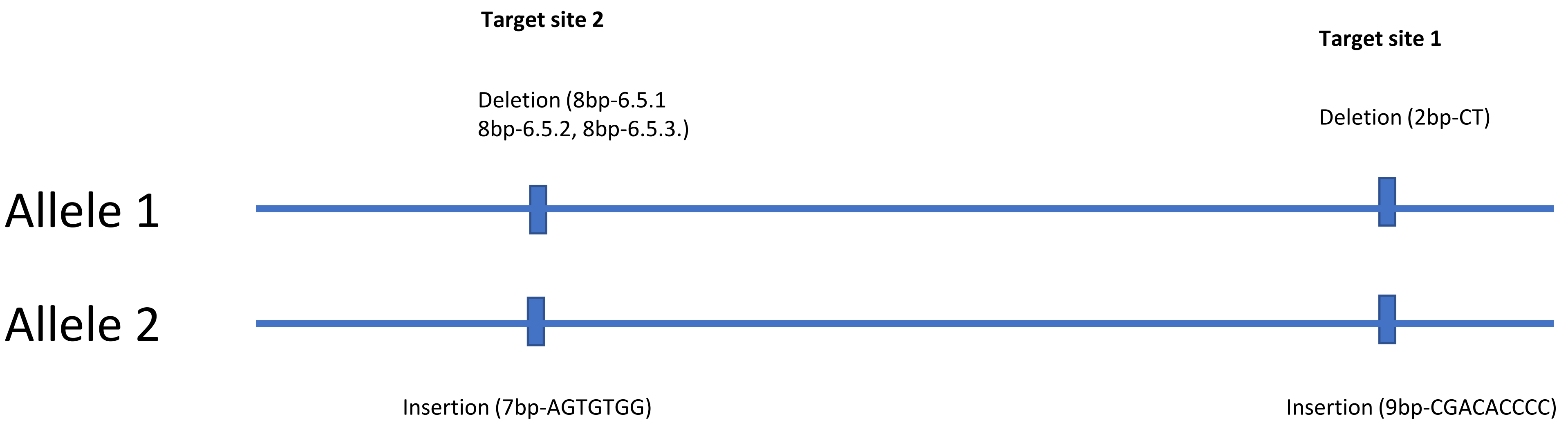

Target site 2
Deletion (8bp-6.5.1
8bp-6.5.2, 8bp-6.5.3.)
Target site 1
Deletion (2bp-CT)
Allele 1
Allele 2
Insertion (7bp-AGTGTGG)
Insertion (9bp-CGACACCCC)

# KO6.5.1

Deletion-(8bp-GTGGTCGA)-4 reads

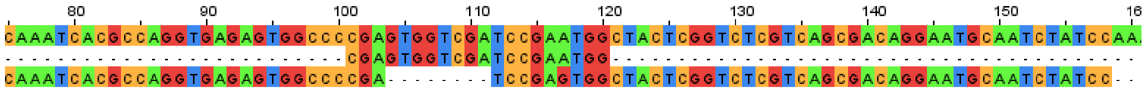

Deletion-(2bp-CT)-16 reads

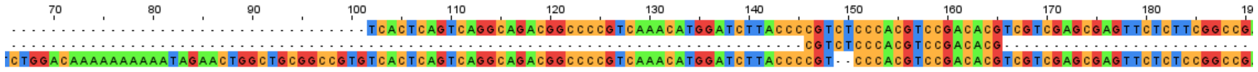

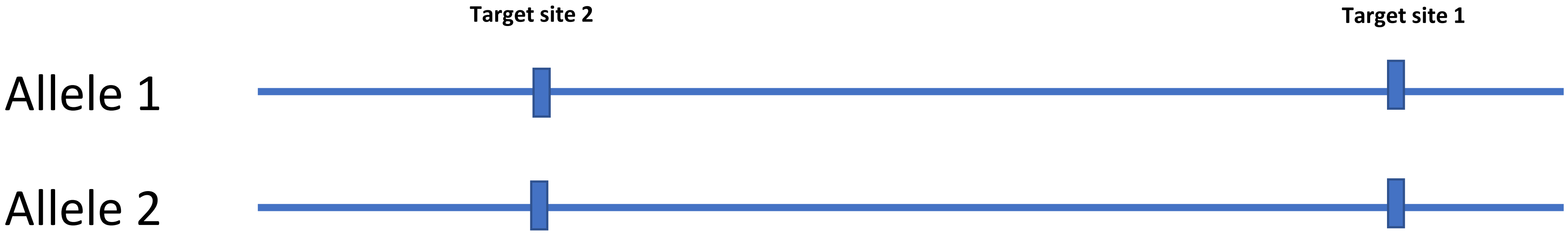


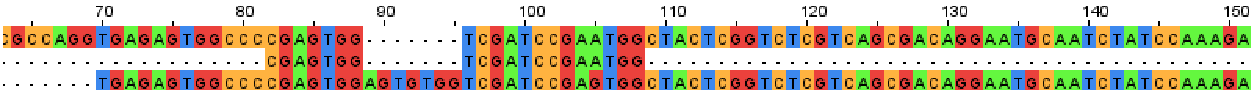

Insertion-(7bp-AGTGTGG)-9 reads

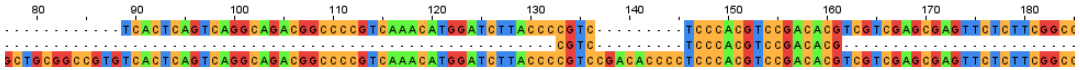

Insertion-(9bp-CGACACCCC)-7 reads

# KO6.5.2

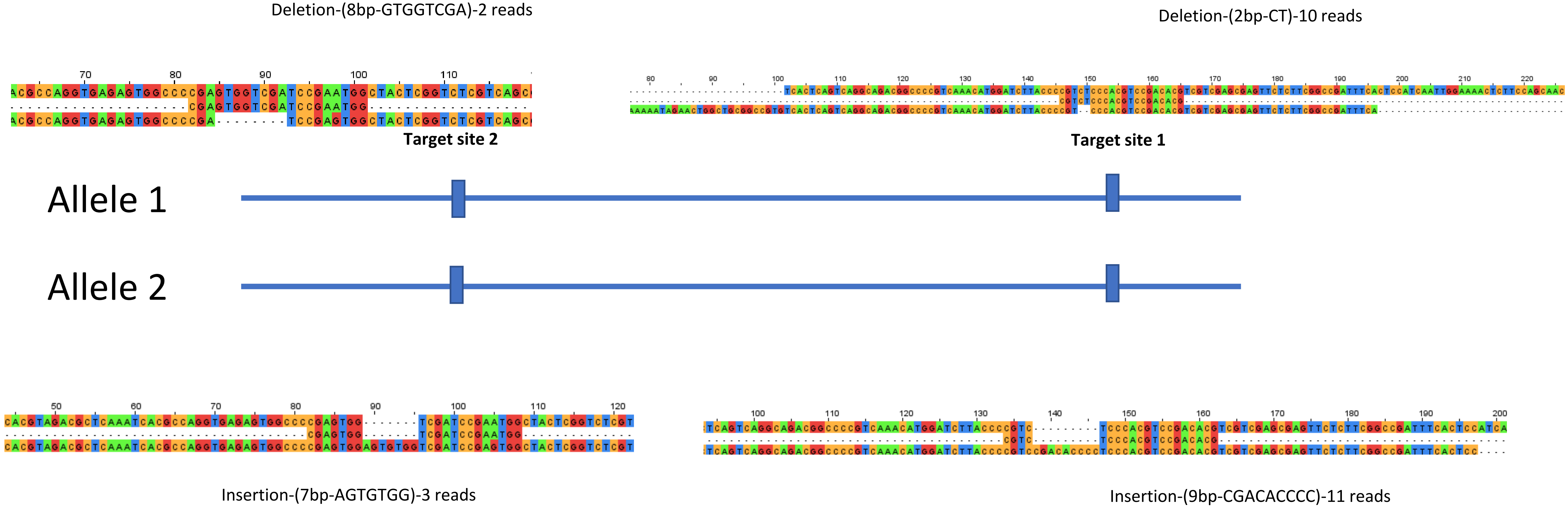
Deletion-(8bp-GTGGTCGA)-2 reads
Deletion-(2bp-CT)-10 reads
Target site 2
Target site 1
Allele 1
Allele 2
Insertion-(7bp-AGTGTGG)-3 reads
Insertion-(9bp-CGACACCCC)-11 reads

# KO6.5.3

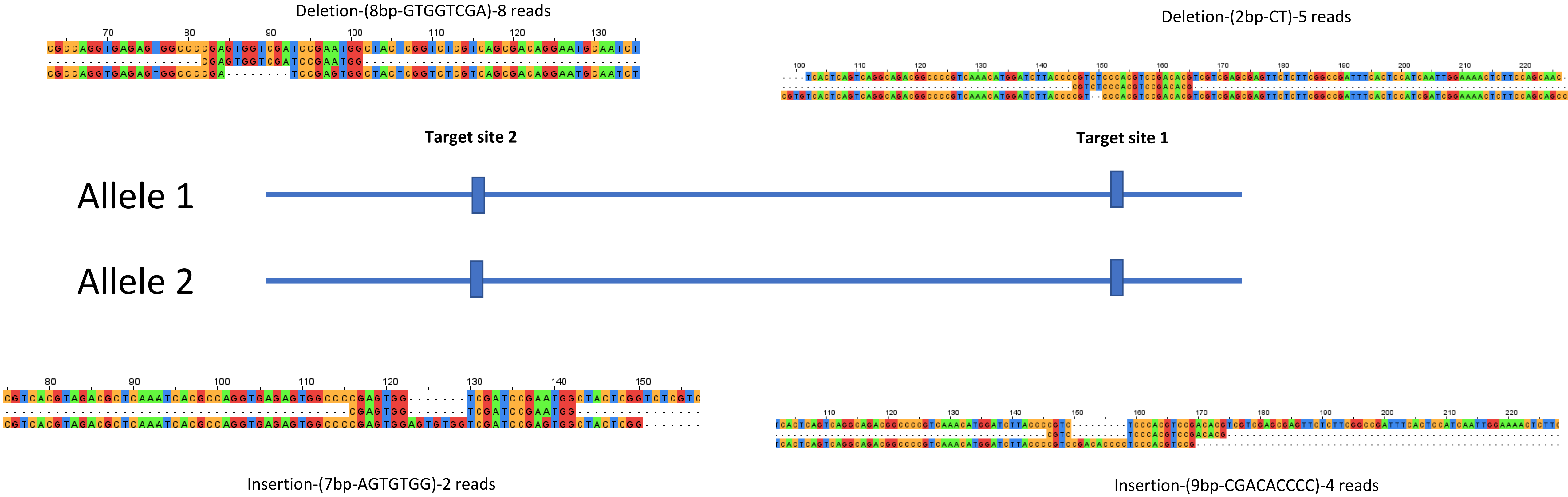
Deletion-(8bp-GTGGTCGA)-8 reads
Deletion-(2bp-CT)-5 reads
Target site 2
Target site 1
Allele 1
Allele 2
Insertion-(7bp-AGTGTGG)-2 reads
Insertion-(9bp-CGACACCCC)-4 reads

### KO7 (Cas12_we1)

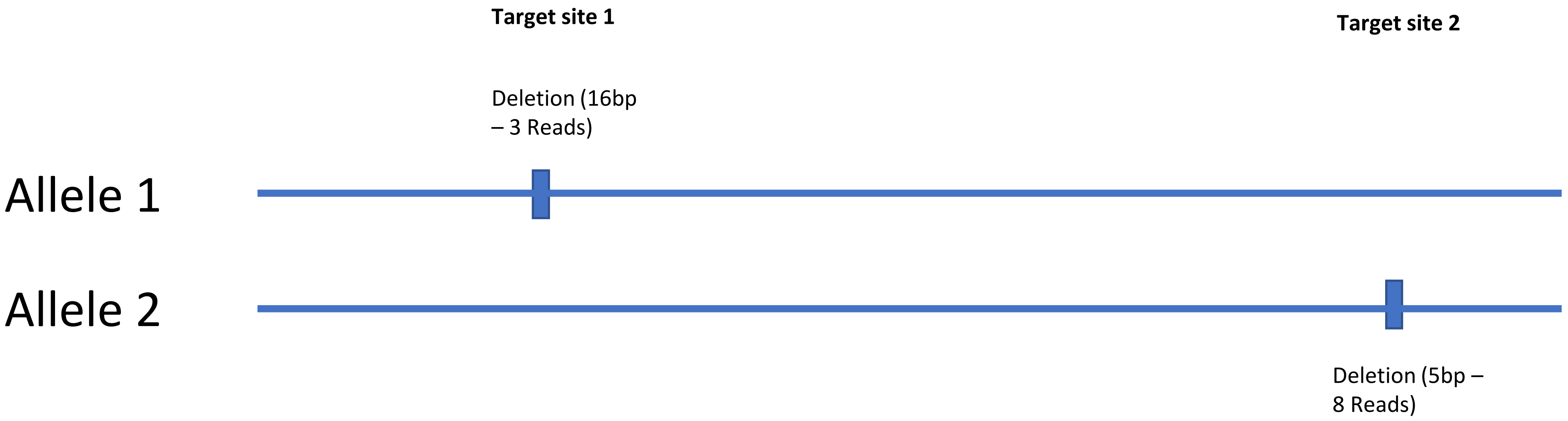

### KO7 (Cas12_we1)

Deletion (16bp-CGTGACGACCCCCAAA ) – 3 reads

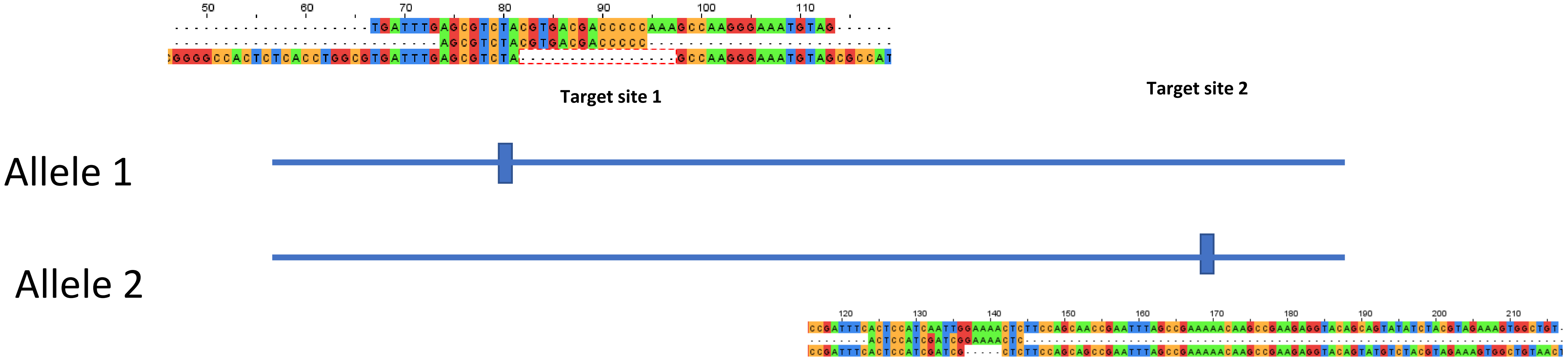


Deletion (5bp-GAAAA)-8 reads)

### KO8(Cas12_we2)

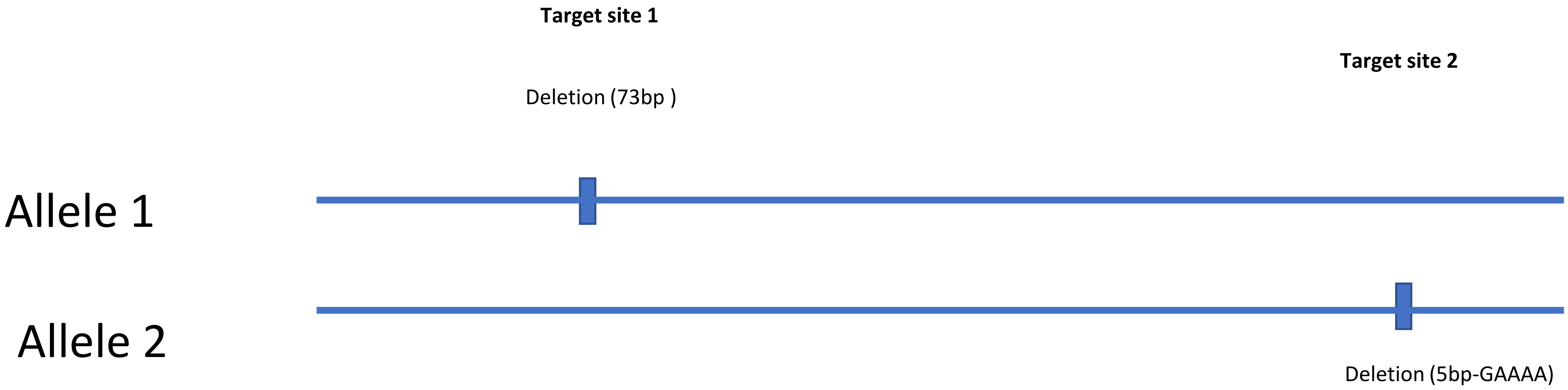

### KO8(Cas12_we2)

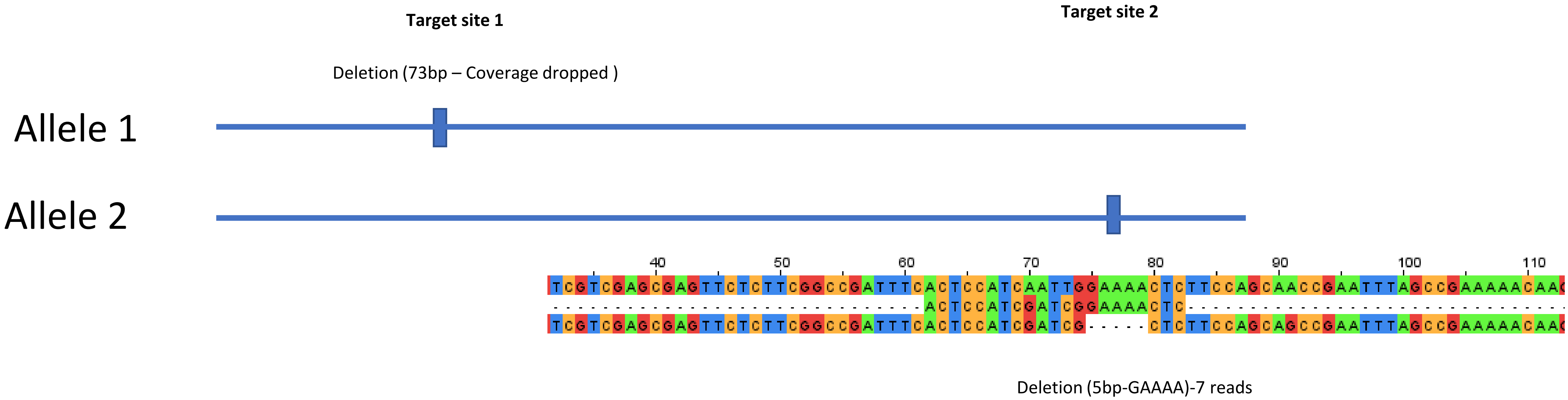

### KO8(Cas12_we2)

**Target site 1**

Deletion (73bp – Coverage dropped )

**Target site 2**

Allele 1

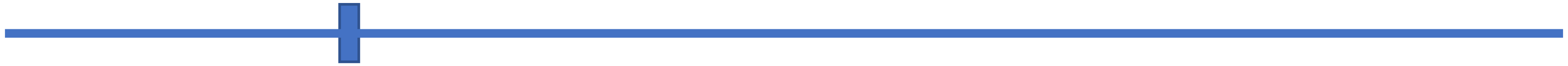

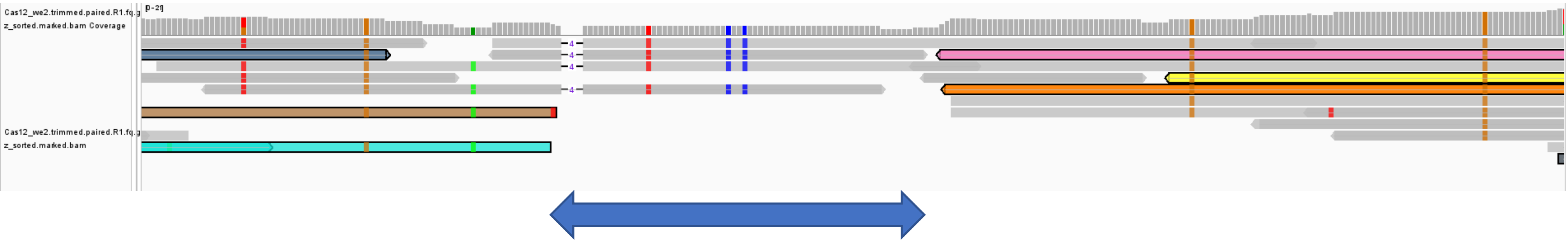


likely deletion region

### KO9(Cas12_we3)

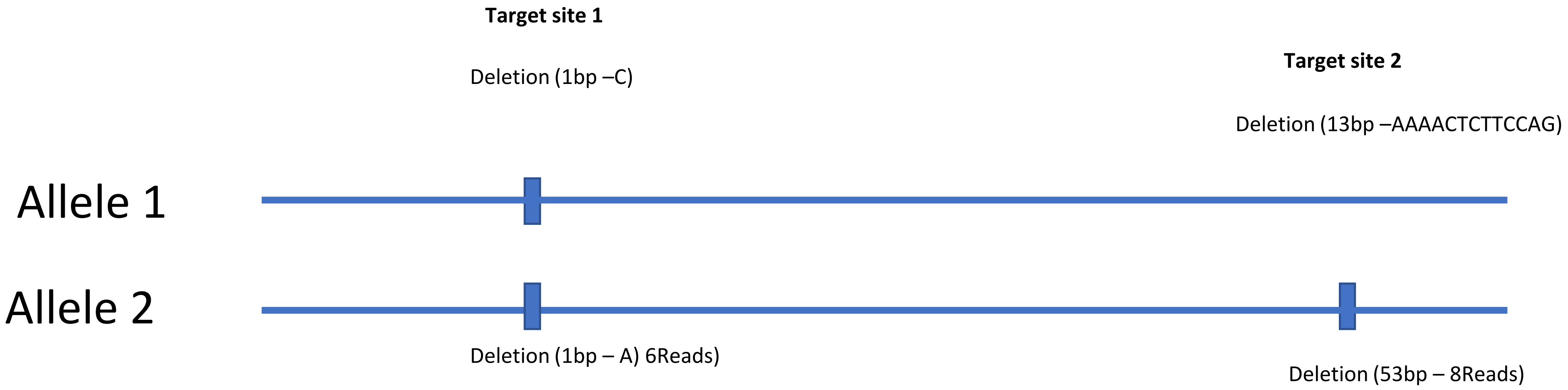

### KO9(Cas12_we3)

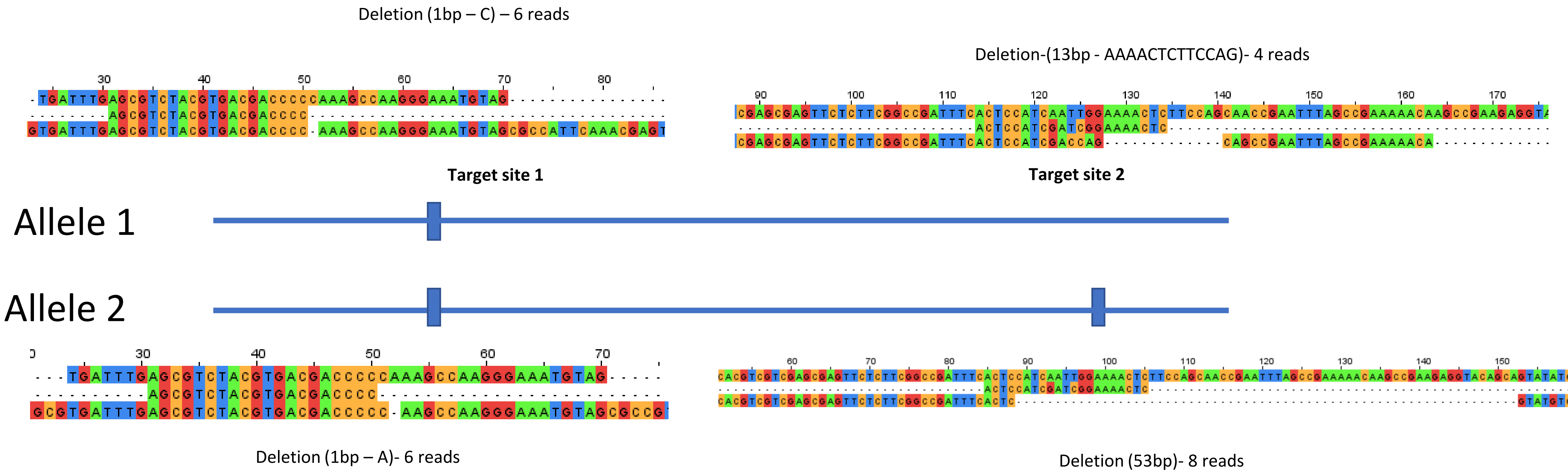

### KO10(Cas12_we4)

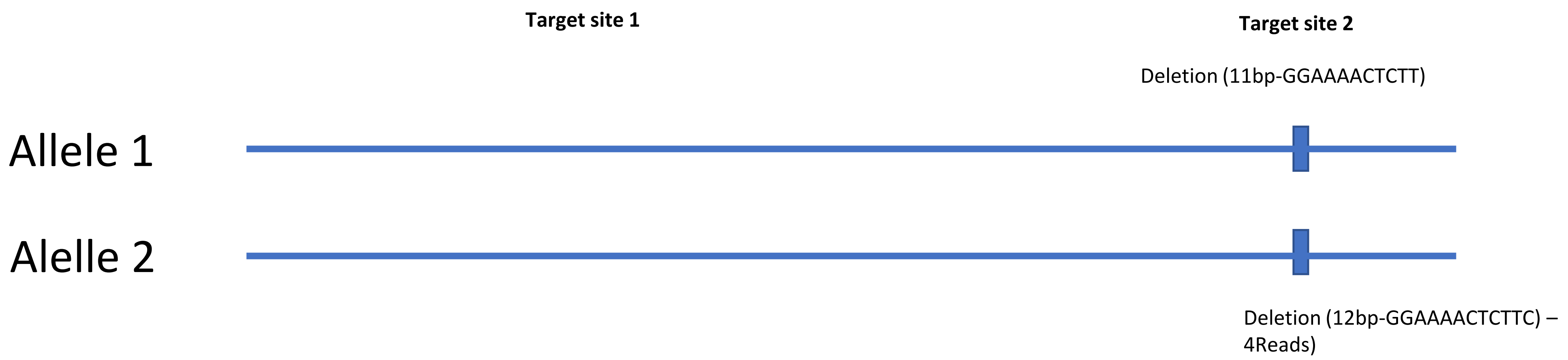

### KO10(Cas12_we4)

Deletion (11bp-GGAAAACTCTT) – 3 reads

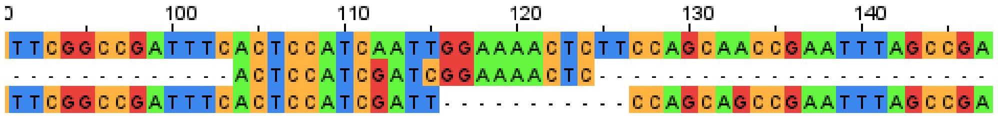


**Target site 1** **Target site 2**

Allele 1

Allele 2

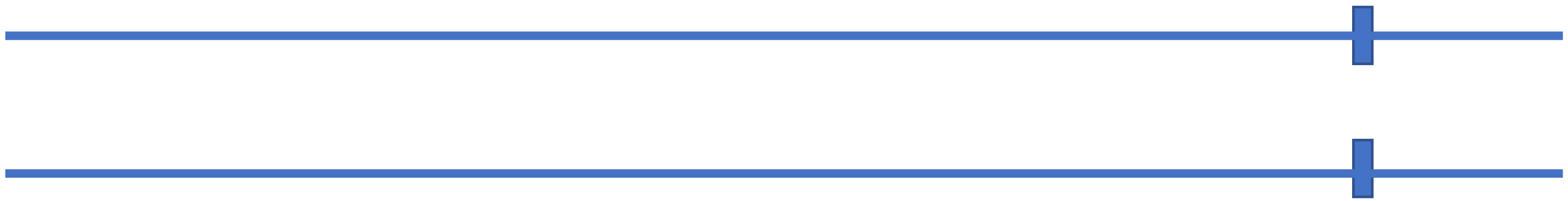

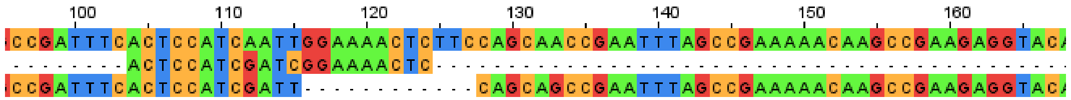


Deletion (12bp-GGAAAACTCTTC) – 4 reads
