## Supplementary materials for "Efficient CRISPR genome editing and integrative genomic analyses reveal the mosaicism of Cas-induced mutations and pleiotropic effects of *scarlet* gene in an emerging model system"

*Notes on the RNA-seq experiments and analysis*

Experimental animals were maintained in the same environmental conditions i.e., 18°C with a 16:8 light/dark cycle for two generations to minimize maternal effects. Three replicates of 2- or 3-day old neonates were collected for each of the seven mutant lines and the wild-type (i.e., control). RNA extractions were completed using the Promega SV Total RNA Isolation kit (Madison, WI, USA) following the manufacturer's instructions. RNA concentration was measured using a Qubit 4.0 Fluorometer (Thermo Scientific, Waltham, MA, USA) and RNA quality was assessed by electrophoresis on a 2% agarose gel. RNA sequencing libraries were prepared using the NEBNext Ultra II RNA Library Prep Kit for Illumina (Ipswich, MA, USA) and sequencing was done by Novogene Corporation Inc. (Sacramento, CA, USA) following standard Illumina sequencing protocol. Each library was sequenced on an Illumina NovaSeq 6000 platform with 150-bp paired-end reads.

Adapter trimming and quality filtering were completed using Trimmomatic v.0.39 (Bolger et al. 2014). with the following parameters: ILLUMINACLIP adapter.fasta:2:30:10:2:keepBothReads LEADING:3 TRAILING:3 MINLEN:36. The adapter.fasta file contained the following Illumina adapter sequences, read 1: AGATCGGAAGAGCACACGTCTGAACTCCAGTCA, read 2: AGATCGGAAGAGCGTCGTGTAGGGAAAGAGTGT. After trimming, the read quality was reassessed using FastQC to confirm the removal of adapter-derived and low-quality sequences.

Cut-off value for co-expression clusters

To select the soft cut-off threshold, we first generated a histogram and log-log plot to visualize the network connectivity. The approximate straight-line relationship shown by the high $R^2$ value

(0.89) in the log-log plot indicates when approximate scale free topology is reached. A soft cut-off threshold of 14 was selected since the power reached a scale-free topology fitting index ($R^2$) > 0.89 when plotting $R^2$ against the soft thresholds (power b). Modules were identified via hierarchical clustering using a dynamic tree cut algorithm and similar modules were merged at 0.25 height of the dendrogram. Modules can be defined as groups of genes with highly correlated expression profiles across mutant lines. The minimum number of genes forming a module was set to 30. Genes with low co-expression and no grouping into a module were added to the grey module.

**Table S1**. Guide RNA target sequence in the scarlet locus and DNA sequence for the HDR template. For the HDM template, green letters indicate stop codon cassette, red letters indicate the crRNA target sites. The homology arms are underlined.

| | **Sequence (5'-3')** |
|---|---|
| **Cas9 knock-out and knock-in** | |
| crRNA1 | CGTCTCCCACGTCCGACACG |
| crRNA2 | CCACTCGGATCGACCACTCG |
| HDR template | CTGTCGCTGACGAGACCGAGTAGCCACTCGGATCGACCACGTCATGGCGTTTAAACCTTAATTA AGCTGTTGTAGTCGGGGCCACTCTCACCTGGCGTGATTTGAGCGTCTACGT |
| **Cas12a knock-out** | |
| crRNA1-cas12 | AGCGTCTACGTGACGACCCCC |
| crRNA2-cas12 | ACTCCATCAATTGGCAAACTC |

**Table S2.** Phenotype of the offspring from two mosaic phenotype individuals

| Individual | Brood | No. of clear-eyed offspring | No. of black-eyed offspring |
|---|---|---|---|
| Mosaic 1 | 1st | 2 | 0 |
| | 2nd | 0 | 4 |
| | 3rd | 0 | 1 |
| | 4th | 1 | 7 |
| Mosaic 2 | 1st | 5 | 0 |
| | 2nd | 6 | 0 |
| | 3rd | 5 | 0 |
| | 4th | 7 | 0 |
| | 5th | 6 | 0 |
| | 6th | 2 | 0 |
| | 7th | 1 | 0 |
| Mosaic 3 | 1st | 3 | 1 |
| | 2nd | 4 | 0 |
| | 3rd | 3 | 2 |
| Mosaic 4 | 1st | 2 | 0 |
| | 2nd | 2 | 3 |
| | 3rd | 3 | 1 |
| | 4th | 4 | 1 |
| | 5th | 4 | 0 |
| Mosaic 5 | 1st | 2 | 2 |
| | 2nd | 3 | 1 |
| | 3rd | 3 | 4 |
| | 4th | 2 | 3 |

**Table S3.** Phenotype of the offspring from mosaic/clear-eye phenotype individuals in Cas12 knock-out experiments.

| Individual | Brood | No. of clear-eyed offspring | No. of black-eyed offspring |
|---|---|---|---|
| Clear-eye 1 | 1st | 0 | 7 |
| | 2nd | 3 | 3 |
| | 3rd | 3 | 3 |
| | 4th | 6 | 7 |
| | 5th | 9 | 7 |
| Mosaic 1 | 1st | 0 | 3 |
| | 2nd | 0 | 4 |
| | 3rd | 0 | 3 |
| | 4th | 0 | 10 |
| | 5th | 0 | 11 |
| Mosaic 2 | 1st | 0 | 2 |
| | 2nd | 0 | 3 |
| | 3rd | 0 | 7 |
| | 4th | 0 | 13 |
| Mosaic 3 | 1st | 4 | 0 |
| | 2nd | 8 | 0 |
| | 3rd | 9 | 0 |
| Mosaic 4 | 1st | 2 | 2 |
| | 2nd | 4 | 0 |
| | 3rd | 4 | 0 |
| | 4th | 4 | 1 |
| | 5th | 11 | 1 |
| Mosaic 5 | 1st | 0 | 4 |
| | 2nd | 0 | 10 |
| | 3rd | 0 | 8 |
| | 4th | 0 | 14 |
| | 5th | 0 | 17 |
| Mosaic 6 | 1st | 0 | 4 |
| | 2nd | 0 | 8 |
| Mosaic 7 | 1st | 4 | 1 |
| | 2nd | 4 | 0 |
| Mosaic 8 | 1st | 0 | 5 |
| | 2nd | 0 | 5 |
| | 3rd | 0 | 15 |

**Table S4.** Summary of Cas9-induced mutations at the scarlet locus. For mutant lines derived from neonates in different broods of the $G_0$ mutants, the second number in their names indicates the number of brood, whereas the third number in their names is their unique identifier. For example, KO5.1.1 means this mutant line is derived from the first neonate of the first brood of the $G_0$ mutant KO5.

| | Target site 2 | Target site 1 |
|---|---|---|
| KO1-allele 1 | Ins (5bp-TCTGA) | Del (9bp-TCCCACGTC) |
| KO1-allele 2 | Ins (13bp-TTGTGGTCGAATT) | None |
| KO2-allele 1 | Del (1569bp) | Del (7bp-GTCTCCC) |
| KO2-allele 2 | Segmental deletion (237bp) | Segmental deletion (237bp) |
| KO3-allele 1 | Ins (17bp-ACGATCCG ATCGAGTGG) | Del (9bp-TCCCACGTC) |
| KO3-allele 2 | Ins (29bp-AGCCCCGAGAGTGGCCCCAGGTGAGAGTG) | Ins (12bp-CGACATGGATCG) |
| KO4-allele 1 | Segmental deletion (234bp) | Segmental deletion (234bp) |
| KO4-allele 2 | Segmental deletion (234bp) | Segmental deletion (234bp) |
| KO5.1.1-allele 1 | Del (3bp-TGG) | Del (8bp-TACCCCGT) |
| KO5.1.1-allele 2 | Del (16bp-TGGTCGATCCGAATGG) | Del (3bp-GTC) |
| KO5.1.2-allele 1 | Del (3bp-TGG) | Del (8bp-TACCCCGT) |
| KO5.1.2-allele 2 | Del (1bp-G) | Del (3bp-GTC) |
| KO6.1.1-allele 1 | Del (8bp-GTGGTCGA) | Del (2bp-CT) |
| KO6.1.1-allele 2 | Ins (7bp-GTGGAGT) | Ins (9bp-CCGACACCC) |
| KO6.1.3-allel 1 | Del (1033bp) | Del (2bp-CT) |
| KO6.1.3-allel 2 | Ins (7bp-AGTGTGG) | Ins (5bp-CCGACACCC) |
| KO6.1.4-allele 1 | Del (10bp-CCCCGAGTGG) | Del (2bp-CT) |
| KO6.1.4-allele 2 | Ins (7bp-AGTGTGG) | Ins (9bp-CCGACACCC) |
| KO6.2.1-allele 1 | Del (10bp-GAGTGGCCCC) | Del (2bp-TC) |
| KO6.2.1-allele 2 | Ins (7bp-AGTGTGG) | Ins (9bp-CCCTCCCAC) |
| KO6.2.2-allele 1 | Del(8bp-GTGGTCGA) | Del (2bp-TC) |
| KO6.2.2-allele 2 | Ins (7bp-AGTGTGG) | Ins (9bp-CCGACACCC) |
| KO6.3.2-allele 1 | Del (10bp-CCCCGAGTGG) | Del (2bp-TC) |
| KO6.3.2-allale 2 | Ins (7bp-AGTGTGG) | Ins(9bp-CGACACCCC) |
| KO6.3.3-allele 1 | Del (1035bp) | Del (2bp-TC) |
| KO6.3.3-allele 2 | Ins (7bp-AGTGTGG) | Ins (9bp-CGACACCCC) |
| KO6.3.4-allele 1 | Del (1024bp) | Del (2bp-TC) |
| KO6.3.4-allele 2 | Ins (7bp- AGTGTGG) | Ins (9bp-CGACACCCC) |
| KO6.4.1-allele 1 | Del (10bp-CCCCGAGTGG) | Del (2bp-TC) |
| KO6.4.1-allele 2 | Ins (7bp-AGTGTGG) | Ins (9bp- CGACACCCC) |
| KO6.5.1-allele 1 | Del(8bp-GTGGTCGA) | Del (2bp-CT) |
| KO6.5.1-allele 2 | Ins (7bp-AGTGTGG) | Ins (9bp-CGACACCCC) |
| KO6.5.2-allele 1 | Del(8bp-GTGGTCGA) | Del (2bp-CT) |
| KO6.5.2-allele 2 | Ins (7bp-AGTGTGG) | Ins (9bp-CGACACCCC) |
| KO6.5.3-allele 1 | Del(8bp-GTGGTCGA) | Del (2bp-CT) |
| KO6.5.3-allele 2 | Ins (7bp-AGTGTGG) | Ins (9bp-CGACACCCC) |

**Table S5.** Summary of Cas12a-induced mutations at the scarlet locus.

| | **Target site 1** | **Target site 2** |
|---|---|---|
| KO7-allele 1 | Del (16bp-CGTGACGACCCCCAAA) | None |
| KO7-allele 2 | None | Del (5bp-GAAAA) |
| KO8-allele 1 | Del (73bp) | None |
| KO8-allele 2 | None | Del (5bp-GAAAA) |
| KO9-allele 1 | Del (1bp-C) | Del (13bp-AAAACTCTTCCAG) |
| KO9-allele 2 | Del (1bp-A) | Del (53bp) |
| KO10-allele 1 | None | Del (11bp-GGAAAACTCTT) |
| KO10-allele 2 | None | Del (12bp-GGAAAACTCTTC) |

**Table S6.** Summary of Cas9-induced mutations at target site 2 in the scarlet locus.

| | **Target site 2** |
|---|---|
| KI1-allele 1 | Segmental deletion (>1000bp) |
| KI1-allele 2 | Ins (8bp-CCAGGTGAC) |
| KI2-allele 1 | Ins (4bp-CGAT) |
| KI2-allele 2 | Ins (5bp-TCCGT) |
| KI3-allele 1 | Del (9bp-GTGGTCGAT) |
| KI3-allele 2 | Del (23bp-CCCCGAGTGGTCGATCCGAATGG) |
| KI4-allele 1 | See Figure 5 |
| KI4-allele 2 | Ins (7bp-CCCCGAC) |
| KI6-allele 1 | See Figure 5 |
| KI6-allele 2 | Ins (7bp-CCCCGAC) |

**Table S7**. Summary of base-substitution mutations identified in the scarlet Cas9/Cas12a mutant lines.

| **Mutant line** | **Total mutations** | **Transitions** | **Transversions** | **Mutation rate (/base/generation)** |
|---|---|---|---|---|
| KO1 | 1 | 1 | 0 | $2.1 \times 10^{-9}$ |
| KI1 | 2 | 1 | 1 | $4.2 \times 10^{-9}$ |
| KO6.1.1 | 1 | 0 | 1 | $2.1 \times 10^{-9}$ |
| KO6.1.3 | 3 | 2 | 1 | $6.2 \times 10^{-9}$ |
| KO6.1.4 | 2 | 0 | 2 | $4.2 \times 10^{-9}$ |
| KO6.4.1 | 1 | 1 | 0 | $2.1 \times 10^{-9}$ |
| KO9 | 1 | 0 | 1 | $2.1 \times 10^{-9}$ |

**Table S8.** Summary of detected SVs in *scarlet* mutants.

| Mutant | Insertion | Deletion | Duplications |
|---|---|---|---|
| KO1 | 0 | 1 (54bp) | 0 |
| KI3 | 0 | 2 (236bp, 98bp) | 0 |
| KI4 | 0 | 1 (100bp) | 0 |
| KI6 | 1 (113bp) | 0 | 0 |
| KO5.1.1 | 0 | 1 (552bp) | 0 |
| KO6.4.1 | 0 | 0 | 1 (297bp) |
| KO9 | 0 | 1 (23418bp) | 1 (1735bp) |

**Table S9.** Structural variation (SV) rate (/bp/generation) in the scarlet mutant lines.

| **Mutants** | KO1 | KI3 | KI4 | KI6 | KO5.1.1 | KO6.4.1 | KO9 |
|---|---|---|---|---|---|---|---|
| **Total deletion bases (bp)** | 54 | 334 | 100 | 0 | 552 | 0 | 23418 |
| **Total insertion bases (bp)** | 0 | 0 | 0 | 148 | 0 | 0 | 0 |
| **Total duplicated bases** | 0 | 0 | 0 | 0 | 0 | 297 | 1735 |
| **Deletion rate** | 1.12E-7 | 6.94E-7 | 2.08E-7 | 0 | 1.15E-6 | 0 | 4.86E-5 |
| **Insertion rate** | 0 | 0 | 0 | 3.07E-07 | 0 | 0 | 0 |
| **Duplication rate** | 0 | 0 | 0 | 0 | 0 | 6.17E-07 | 3.60E-6 |

**Table S10**. RNA-seq raw read information for the wildtype and mutants. Mapped reads refer to the reads mapped to the reference assembly, whereas the assigned reads represent those assigned to genes and counted towards transcript abundance.

| Sample | Total reads | Mapped reads | % Mapped | Assigned reads | % Assigned |
|---|---|---|---|---|---|
| EB1_1 | 21633708 | 19556862 | 90.40% | 15001364 | 76.70% |
| EB1_2 | 23697993 | 21425688 | 90.40% | 17467448 | 81.50% |
| EB1_3 | 18827901 | 17409826 | 92.50% | 14171239 | 81.40% |
| K01_1 | 22752251 | 16501793 | 72.50% | 10689811 | 64.80% |
| K01_2 | 19499336 | 18389992 | 94.30% | 14212611 | 77.30% |
| K01_3 | 18584770 | 17572245 | 94.60% | 13737910 | 78.20% |
| K02_1 | 22756640 | 21655509 | 95.20% | 17024560 | 78.60% |
| K02_2 | 20004832 | 18537508 | 92.70% | 14902569 | 80.40% |
| K02_3 | 17200229 | 16382627 | 95.20% | 13048123 | 79.60% |
| K03_1 | 23112912 | 21784315 | 94.30% | 17061204 | 78.30% |
| K03_2 | 28239190 | 25743251 | 91.20% | 20628262 | 80.10% |
| K03_3 | 20872801 | 19645596 | 94.10% | 15228723 | 77.50% |
| K04_1 | 16605506 | 15496413 | 93.30% | 12141580 | 78.40% |
| K04_2 | 17935652 | 16993776 | 94.70% | 13346137 | 78.50% |
| K04_3 | 20017175 | 18138361 | 90.60% | 14754930 | 81.30% |
| KI2_1 | 19331295 | 17487253 | 90.50% | 14275134 | 81.60% |
| KI2_2 | 28120289 | 25719637 | 91.50% | 20824239 | 81.00% |
| KI2_3 | 24716619 | 22870360 | 92.50% | 18508004 | 80.90% |
| KI3_1 | 20897809 | 18934374 | 90.60% | 15342953 | 81.00% |
| KI3_2 | 20897800 | 18929584 | 90.60% | 15346271 | 81.10% |
| KI3_3 | 18032118 | 13001724 | 72.10% | 8251995 | 63.50% |
| KI4_1 | 16142637 | 14810721 | 91.70% | 11566961 | 78.10% |
| KI4_2 | 15372255 | 14171167 | 92.20% | 11074284 | 78.10% |
| KI4_3 | 26837472 | 24776823 | 92.30% | 20235721 | 81.70% |

**Supplementary Table S11:** Number of upregulated and downregulated genes in each mutant line. Pooled represents the analysis where all mutants were pooled in comparison to the wildtype.

| Mutant | Total | Direction | Number |
|---|---|---|---|
| K01 | 1239 | Downregulated | 474 |
| | | Upregulated | 765 |
| K02 | 1291 | Downregulated | 410 |
| | | Upregulated | 881 |
| K03 | 703 | Downregulated | 250 |
| | | Upregulated | 453 |
| K04 | 940 | Downregulated | 394 |
| | | Upregulated | 546 |
| KI2 | 1393 | Downregulated | 616 |
| | | Upregulated | 777 |
| KI3 | 1822 | Downregulated | 711 |
| | | Upregulated | 1111 |
| KI4 | 938 | Downregulated | 329 |
| | | Upregulated | 609 |
| Pooled | | Downregulated | 112 |
| | | Upregulated | 328 |

**Supplementary Table S12.** Go term and KEGG enrichment analysis for the differentially expressed genes in the pooled analyses.

| GO.ID | Term | Annotated | Significant | Expected | weightFisher |
|---|---|---|---|---|---|
| GO:0006689 | Ganglioside catabolic process | 5 | 3 | 0.13 | 0.00015 |
| GO:0005975 | carbohydrate metabolic process | 272 | 18 | 6.91 | 0.00027 |
| GO:0009448 | gamma-aminobutyric acid metabolic process | 5 | 2 | 0.13 | 0.0061 |
| GO:0016998 | cell wall macromolecule catabolic process | 32 | 4 | 0.81 | 0.00829 |
| GO:0006693 | prostaglandin metabolic process | 8 | 2 | 0.2 | 0.01623 |
| GO:0008272 | sulfate transport | 22 | 3 | 0.56 | 0.01738 |
| GO:0006691 | leukotriene metabolic process | 9 | 2 | 0.23 | 0.02052 |
| GO:0019530 | taurine metabolic process | 9 | 2 | 0.23 | 0.02052 |
| GO:0019482 | beta-alanine metabolic process | 10 | 2 | 0.25 | 0.02523 |
| GO:0000079 | regulation of cyclin-dependent protein serine/threonin | 10 | 2 | 0.25 | 0.02523 |

| | | | | | |
|---|---|---|---|---|---|
| | e kinase activity | | | | |
| GO:0006270 | DNA replication initiation | 11 | 2 | 0.28 | 0.03033 |
| GO:0006730 | one-carbon metabolic process | 31 | 3 | 0.79 | 0.043 |
| GO:0006807 | nitrogen compound metabolic process | 3455 | 113 | 87.73 | 0.04577 |
| GO:1901990 | regulation of mitotic cell cycle phase transition | 14 | 2 | 0.36 | 0.04776 |
| GO:0023052 | signaling | 972 | 9 | 24.68 | 0.08246 |
| GO:0006522 | alanine metabolic process | 19 | 2 | 0.48 | 0.08268 |
| GO:0006531 | aspartate metabolic process | 19 | 2 | 0.48 | 0.08268 |
| GO:0030149 | sphingolipid catabolic process | 9 | 4 | 0.23 | 0.09621 |
| GO:0051301 | cell division | 21 | 2 | 0.53 | 0.09829 |
| GO:0006812 | monoatomic cation transport | 254 | 7 | 6.45 | 0.10036 |

| KEGG pathway | Significant genes | Total genes | p.adjust |
|---|---|---|---|
| 4974 Protein digestion and absorption | 32 | 207 | 0 |
| 4972 Pancreatic secretion | 30 | 207 | 0 |
| 5164 Influenza A | 17 | 146 | 0 |
| 4142 Lysosome | 15 | 174 | 0.0016 |
| 4080 Neuroactive ligand-receptor interaction | 13 | 149 | 0.0039 |
| 4514 Cell adhesion molecules | 10 | 107 | 0.0118 |
| 0500 Starch and sucrose metabolism | 6 | 46 | 0.0268 |
| 4614 Renin-angiotensin system | 5 | 41 | 0.0834 |
| 2020 Two-component system | 3 | 22 | 0.3536 |
| 0981 Insect hormone biosynthesis | 3 | 23 | 0.3592 |
| 4612 Antigen processing and presentation | 4 | 46 | 0.4803 |
| 4973 Carbohydrate digestion and absorption | 3 | 30 | 0.6023 |
| 0430 Taurine and hypotaurine metabolism | 2 | 14 | 0.617 |
| 0460 Cyanoamino acid metabolism | 2 | 14 | 0.617 |
| 0532 Glycosaminoglycan biosynthesis - chondroitin sulfate / dermatan sulfate | 2 | 15 | 0.6534 |
| 4977 Vitamin digestion and absorption | 3 | 36 | 0.7111 |
| 4210 Apoptosis | 11 | 263 | 0.7255 |
| 0770 Pantothenate and CoA biosynthesis | 2 | 18 | 0.7255 |

**Supplementary Table S13.** Summary of the consensus set of differentially expressed genes in comparing each scarlet mutant with the wildtype genotype.

| **GeneID** | **Direction** | **Gene symbol** | **Gene name** |
|---|---|---|---|
| gene10122 | Down | GGT1_5 | gamma-glutamyltranspeptidase |
| gene16728 | Down | CA | carbonic anhydrase |
| gene11677 | Down | Unknown | unknown |
| gene4054 | Down | SLIT2 | slit guidance ligand 2 |
| gene11607 | Down | KLKB1 | plasma kallikrein |
| gene15334 | Up | MAN | mannan endo-1,4-beta-mannosidase |
| gene13459 | Up | LCT | lactase-phlorizin hydrolase |
| gene17518 | Up | LIPA | lysosomal acid lipase |
| gene17448 | Up | ABAT | 4-Aminobutyrate Aminotransferase |
| gene6033 | Up | PNLIPRP1 | pancreatic lipase-related protein 1 |
| gene6036 | Up | PNLIPRP1 | pancreatic lipase-related protein 1 |
| gene9527 | Up | E3.2.1.4 | endoglucanase |
| gene14470 | Up | CTSD | cathepsin D |
| gene13917 | Up | PRSS1_2_3 | trypsin |
| gene14701 | Up | PRSS1_2_3 | trypsin |
| gene16855 | Up | PRSS1_2_3 | trypsin |
| gene16863 | Up | PRSS1_2_3 | trypsin |
| gene8308 | Up | PRSS1_2_3 | trypsin |
| gene8309 | Up | PRSS1_2_3 | trypsin |
| gene8228 | Up | CNTNAP2 | contactin associated protein-like 2 |
| gene9262 | Up | SLC12A2 | solute carrier family 12 |
| gene11745 | Up | CTRB | chymotrypsin |
| gene15133 | Up | CPA2 | carboxypeptidase A2 |
| gene15730 | Up | CPA2 | carboxypeptidase A2 |
| gene15732 | Up | CPA1 | carboxypeptidase A1 |
| gene2853 | Up | CTRL | chymotrypsin-like protease |
| gene2857 | Up | CTRL | chymotrypsin-like protease |
| gene2867 | Up | CELA2 | pancreatic elastase II |
| gene15316 | Up | EC 3.2.1.78 | endo-beta-1,4-mannanase |
| gene13658 | Up | TMPRSS11D | transmembrane protease serine 11D |
| gene8314 | Up | E3.4.21.59 | tryptase |

**Supplementary Table S14.** (A) KEGG pathway analysis for upregulated consensus genes in scarlet mutant lines. (B) KEGG pathway analysis for downregulated consensus genes in scarlet mutant lines

| (A) KEGG pathway | p.adjust |
|---|---|
| 4972 Pancreatic secretion | 0 |
| 4974 Protein digestion and absorption | 0 |
| 5164 Influenza A | 0 |
| 4080 Neuroactive ligand-receptor interaction | 3e-04 |
| 4975 Fat digestion and absorption | 0.0657 |
| 0561 Glycerolipid metabolism | 0.1394 |
| 2020 Two-component system | 0.3208 |
| 0650 Butanoate metabolism | 0.3208 |
| 0051 Fructose and mannose metabolism | 0.3208 |
| 4973 Carbohydrate digestion and absorption | 0.3208 |
| 0100 Steroid biosynthesis | 0.3208 |
| 0410 beta-Alanine metabolism | 0.3208 |
| 0250 Alanine, aspartate and glutamate metabolism | 0.3208 |
| 0640 Propanoate metabolism | 0.3208 |
| 4142 Lysosome | 0.3208 |

| (B) KEGG pathway | p.adjust |
|---|---|
| 0430 Taurine and hypotaurine metabolism | 0.0709 |
| 0460 Cyanoamino acid metabolism | 0.0709 |
| 0910 Nitrogen metabolism | 0.0966 |
| 4610 Complement and coagulation cascades | 0.0966 |
| 0590 Arachidonic acid metabolism | 0.1249 |
| 0480 Glutathione metabolism | 0.1754 |
| 3015 mRNA surveillance pathway | 0.1788 |
| 4361 Axon regeneration | 0.1788 |
| 4360 Axon guidance | 0.1832 |
| 3040 Spliceosome | 0.2013 |
| 5202 Transcriptional misregulation in cancer | 0.2013 |
| 3013 RNA transport | 0.2232 |
| 5014 Amyotrophic lateral sclerosis | 0.4323 |
| 5022 Pathways of neurodegeneration - multiple diseases | 0.4323 |
| 1100 Metabolic pathways | 0.4357 |

**Supplementary Figure S1.** Heatmap visualizing the relationship between module eigengenes of each module with the wildtype and scarlet mutant phenotypes. The Spearman correlation value and p-value (in parenthesis) for each module-phenotype association are shown. The color scale (red-blue) represents the strength of the correlation between the modules and phenotypes.

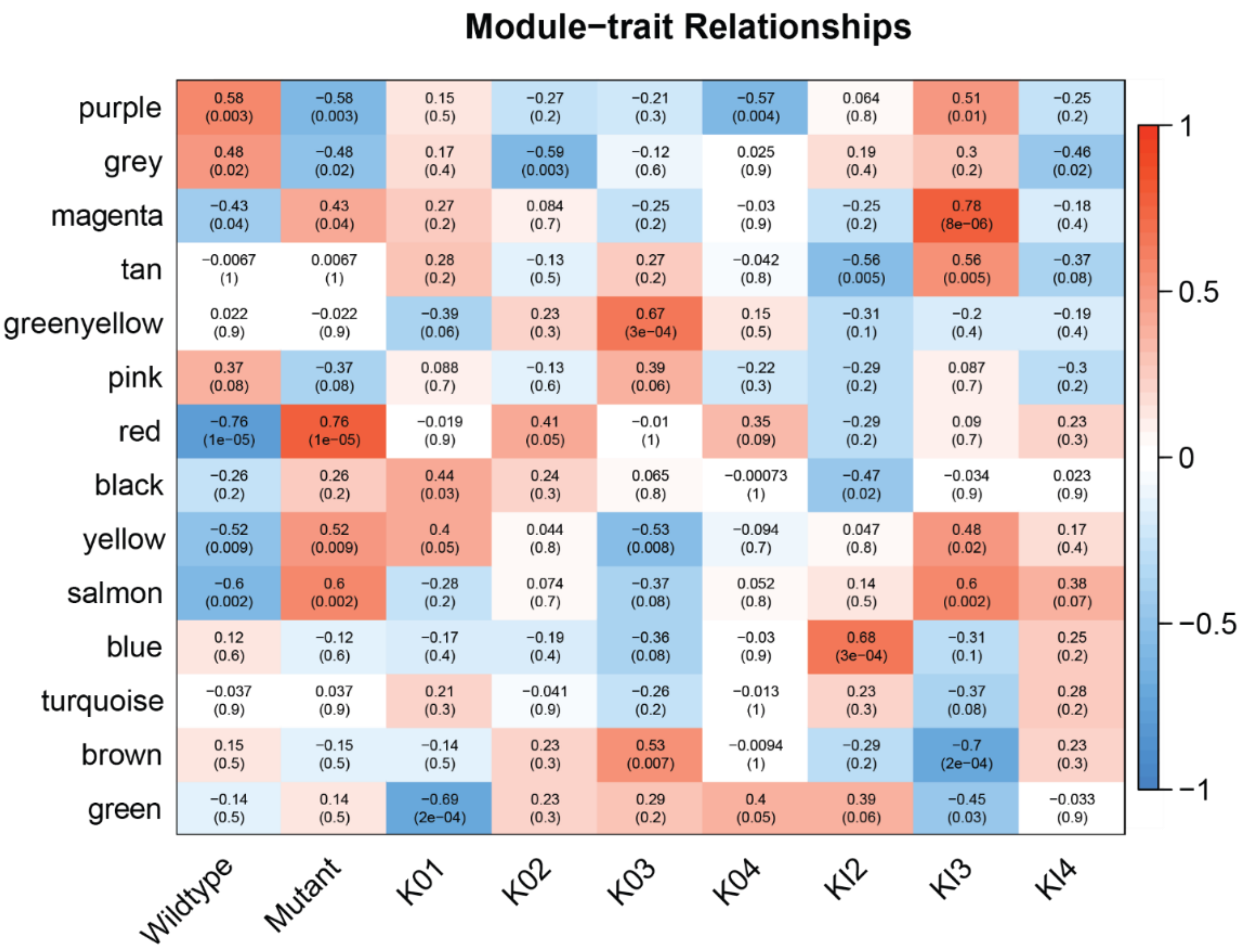

**Supplementary Figure S2**. The Red co-expression module

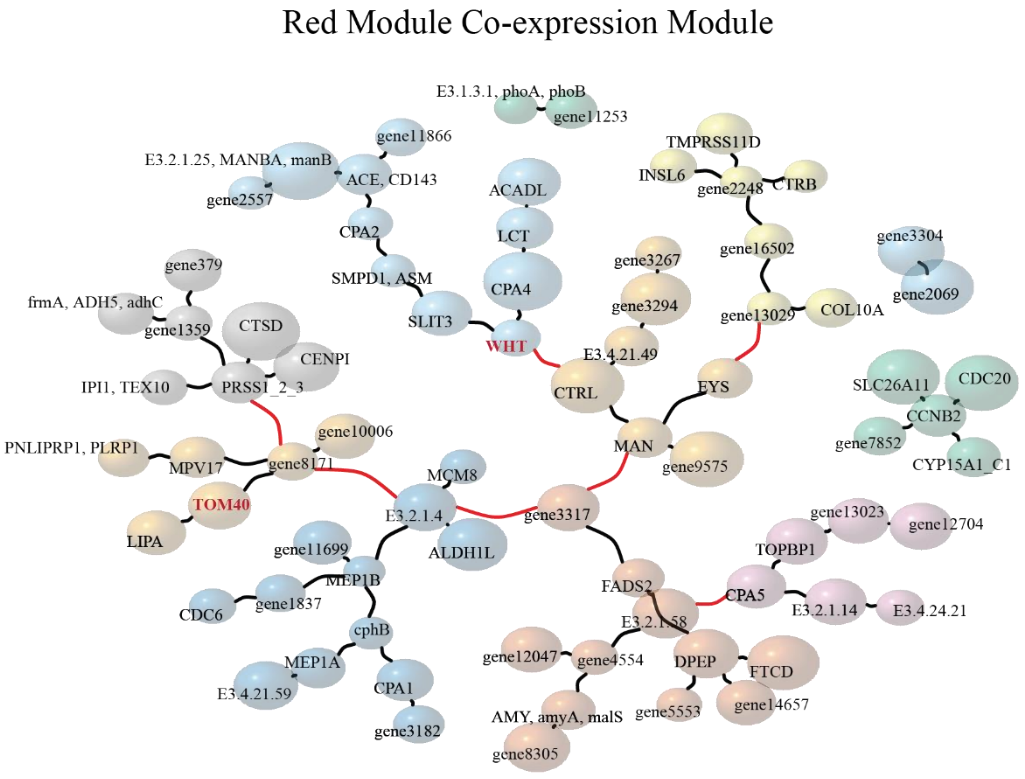

**Supplementary Figure S3.** The magenta co-expression module.

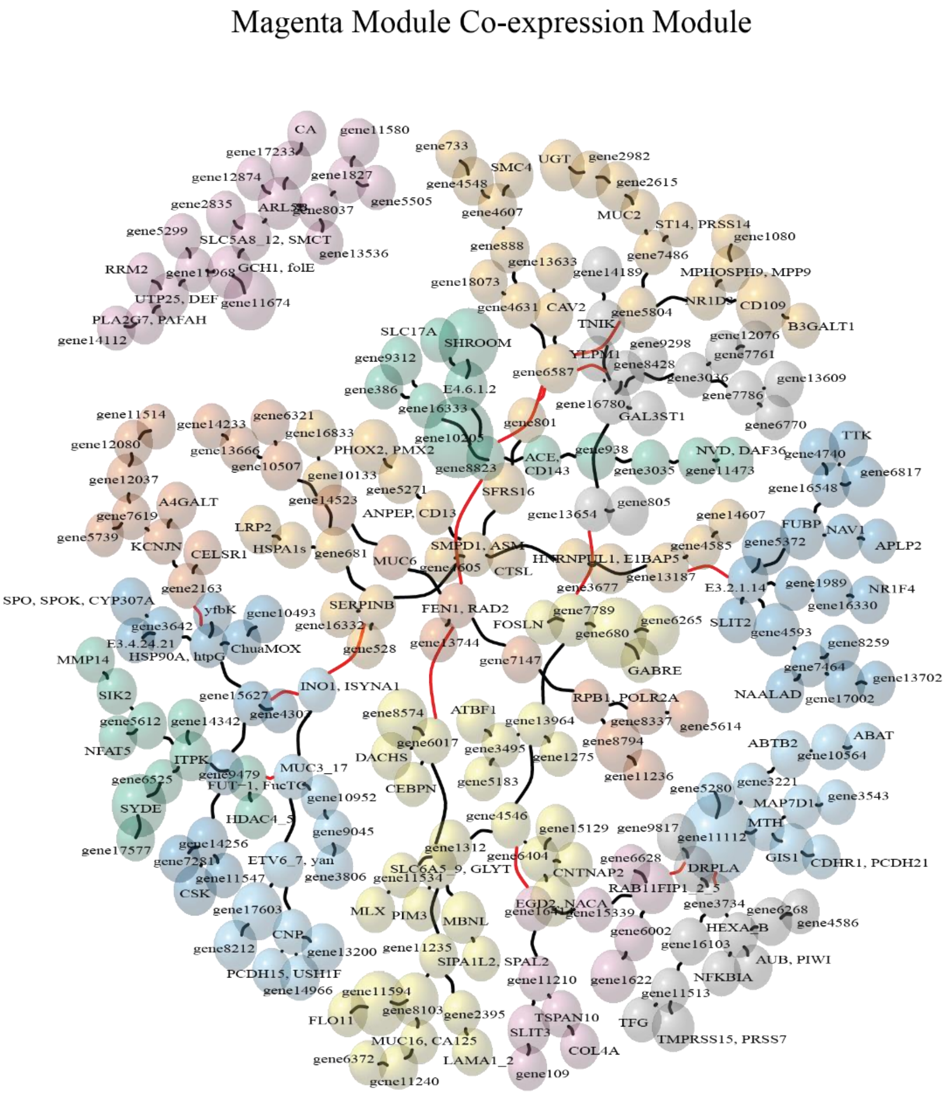

**Supplementary Figure S4.** The yellow co-expression module.

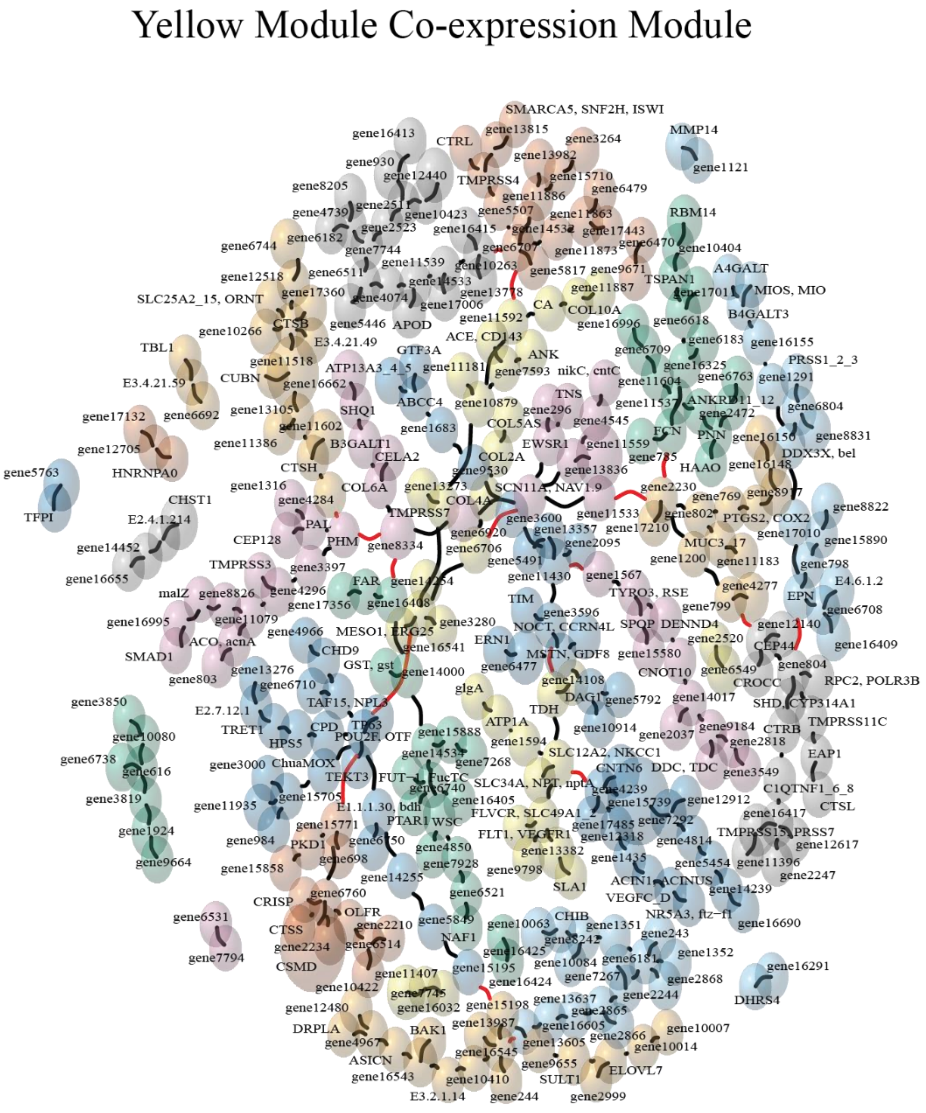

**Supplementary Figure S5.** The salmon co-expression module.

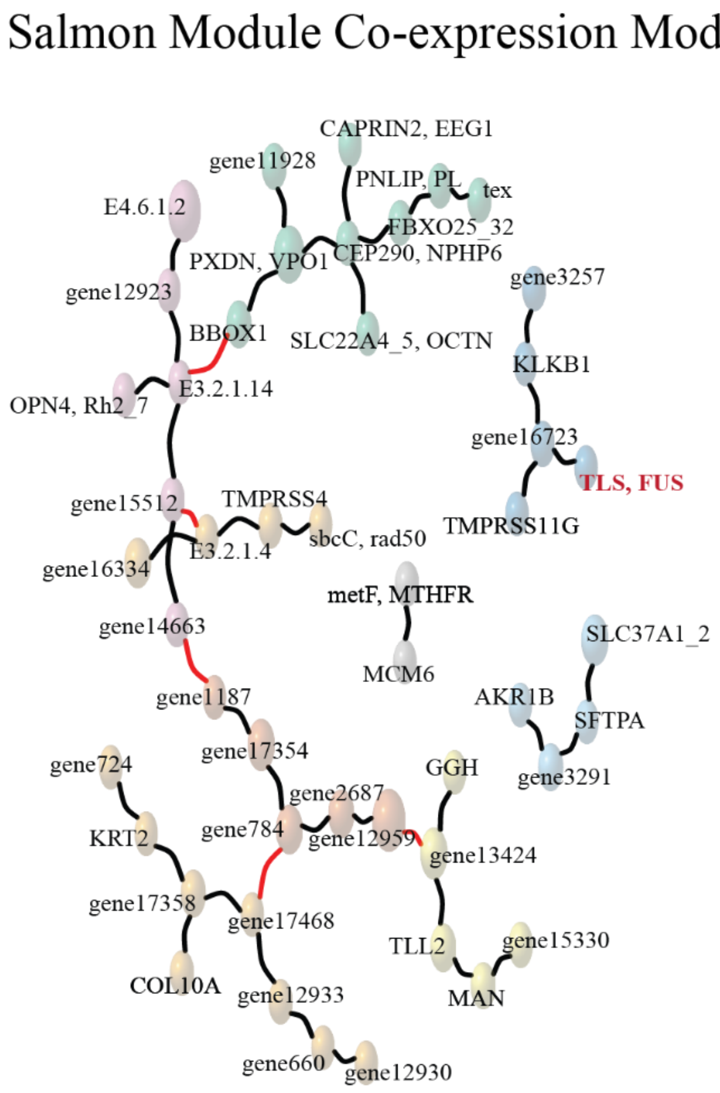

**Supplementary Figure S6.** The purple co-expression module.

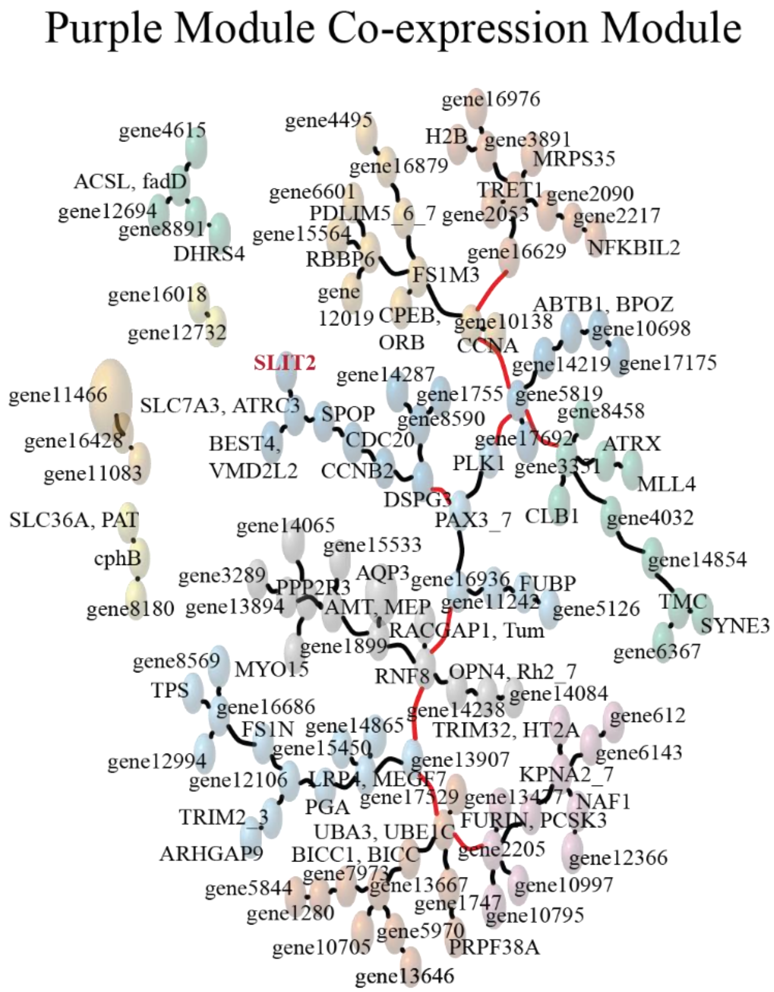
