## Supplementary file 2 for "Efficient CRISPR genome editing and integrative genomic analyses reveal the mosaicism of Cas-induced mutations and pleiotropic effects of *scarlet* gene in an emerging model system"

### Scarlet Knock-in Lines

# KI1

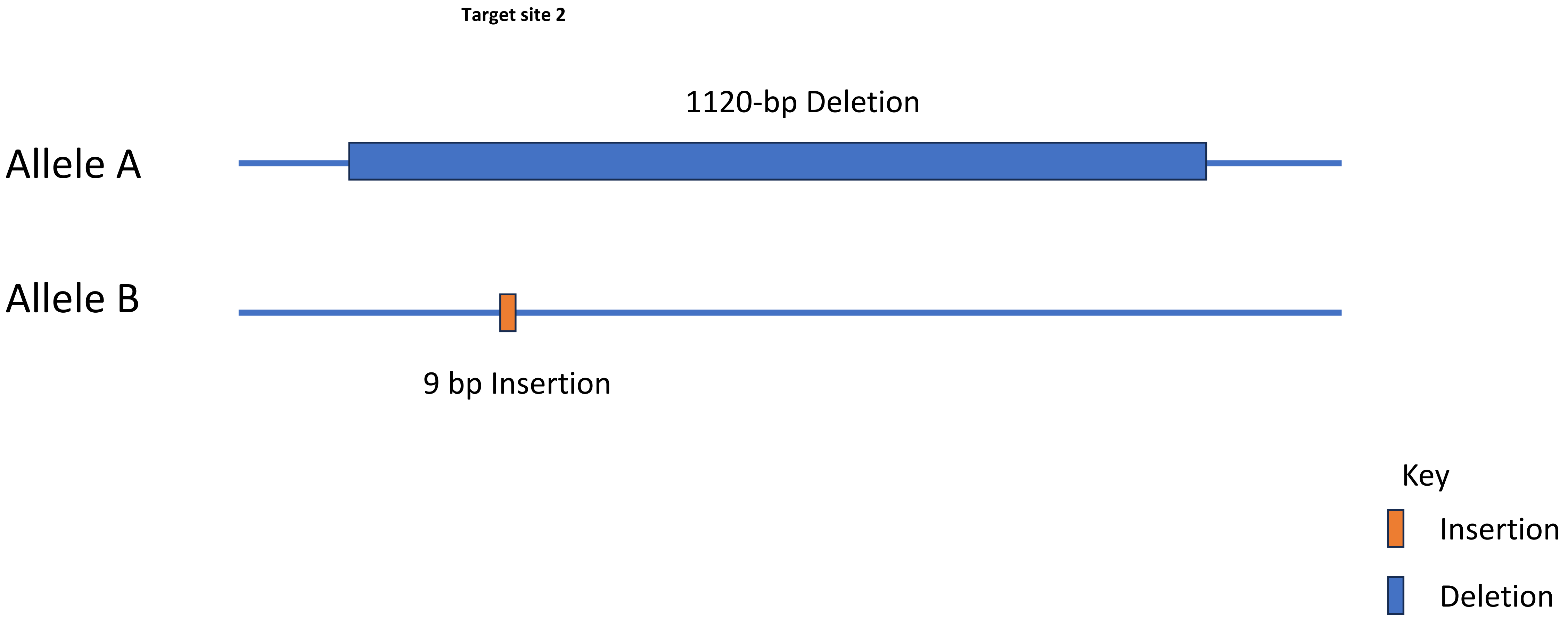
Target site 2
1120-bp Deletion
Allele A
Allele B
9 bp Insertion
Key
Insertion
Deletion

# KI1

**Target site 2**

Supported by 5 Reads
1120-bp Deletion
(635,615-636,735)

Allele 1

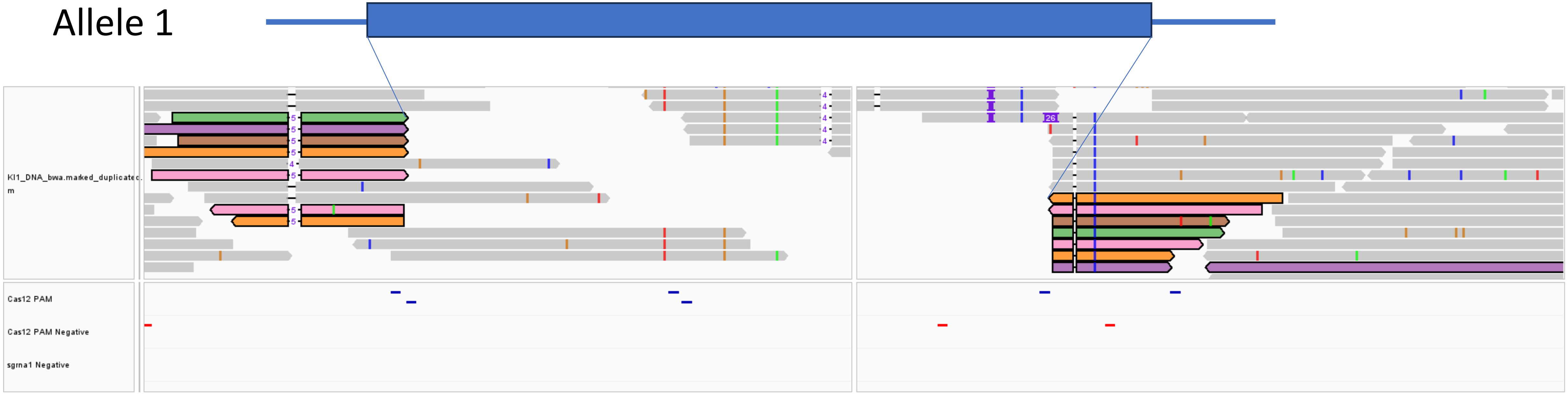

# KI1

**Target site 2**

Supported by 5 Reads
>1000 bp Deletion
(635,615-636,735)

Allele 1

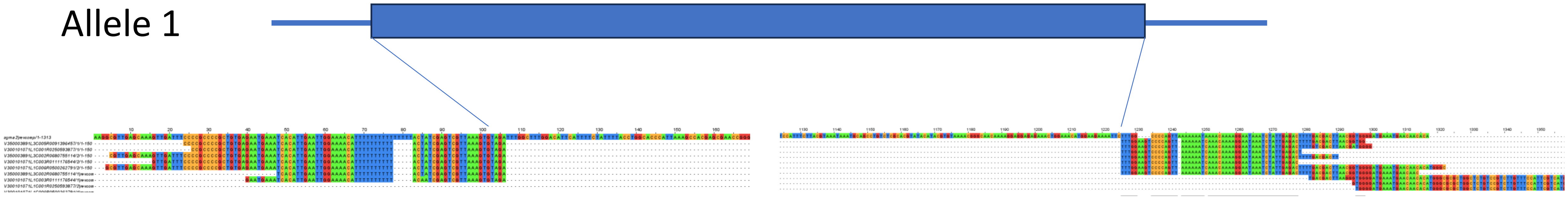

# KI1

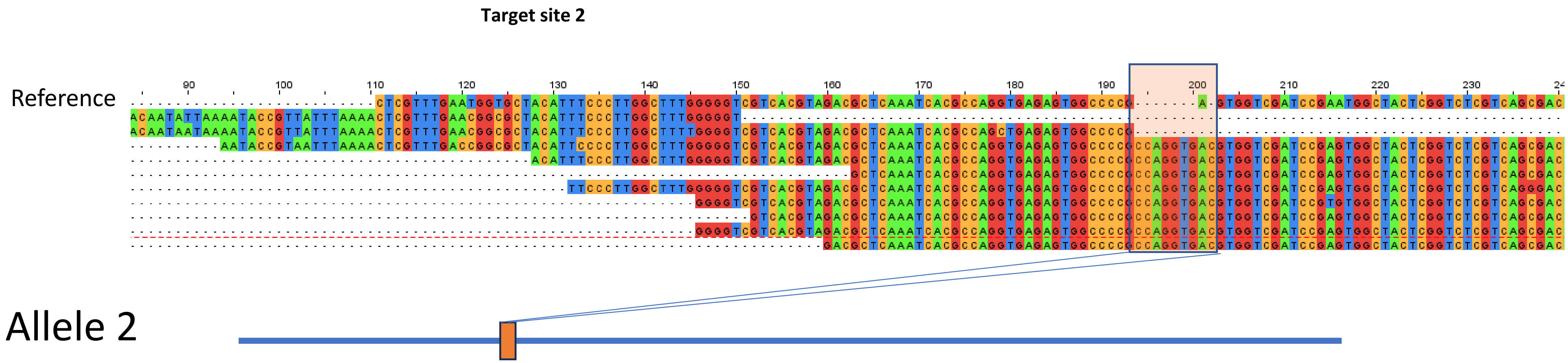


Insertion (9bp-CCAGGTGAC)-10 reads

Key

Insertion

Deletion

# KI2

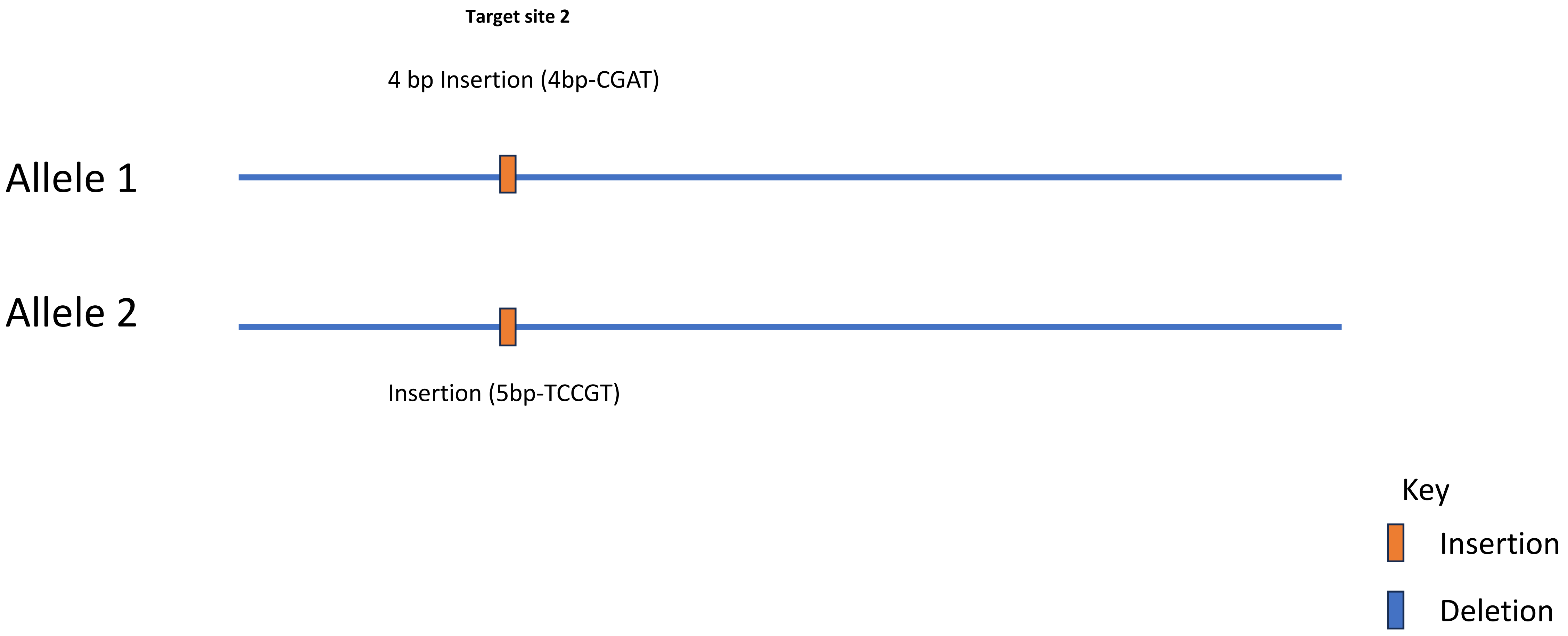

Target site 2
4 bp Insertion (4bp-CGAT)
Allele 1
Allele 2
Insertion (5bp-TCCGT)
Key
Insertion
Deletion

# KI2

**Target site 2**

4 bp Insertion (CGAT)-10 reads

Allele 1

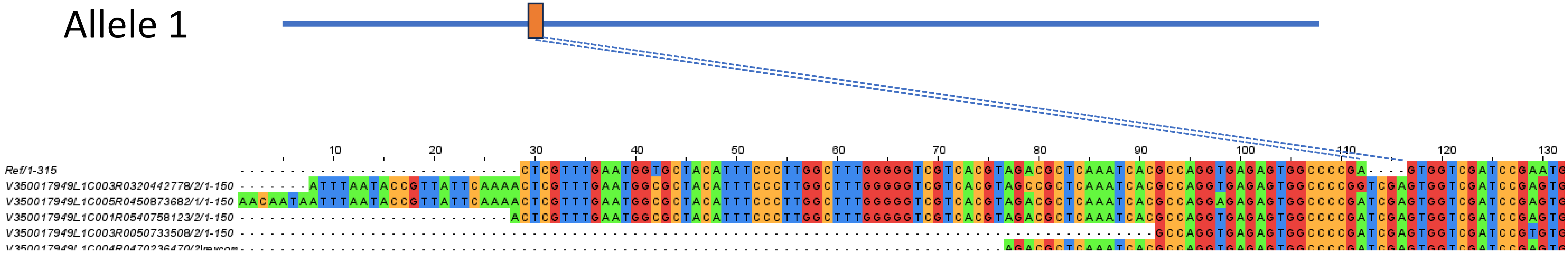

# KI2

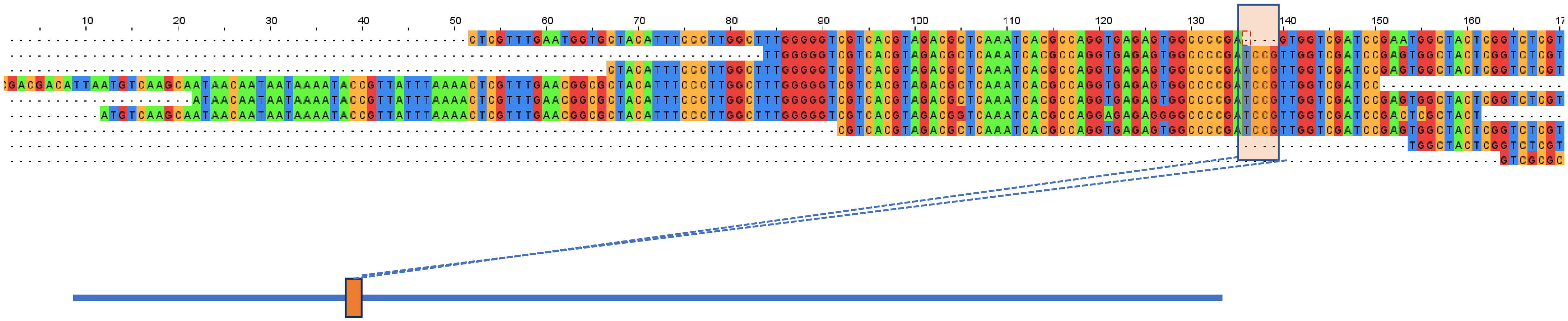


Allele 2

Insertion (5bp-TCCGT)-15 reads

Key

Insertion

Deletion

# Kl3

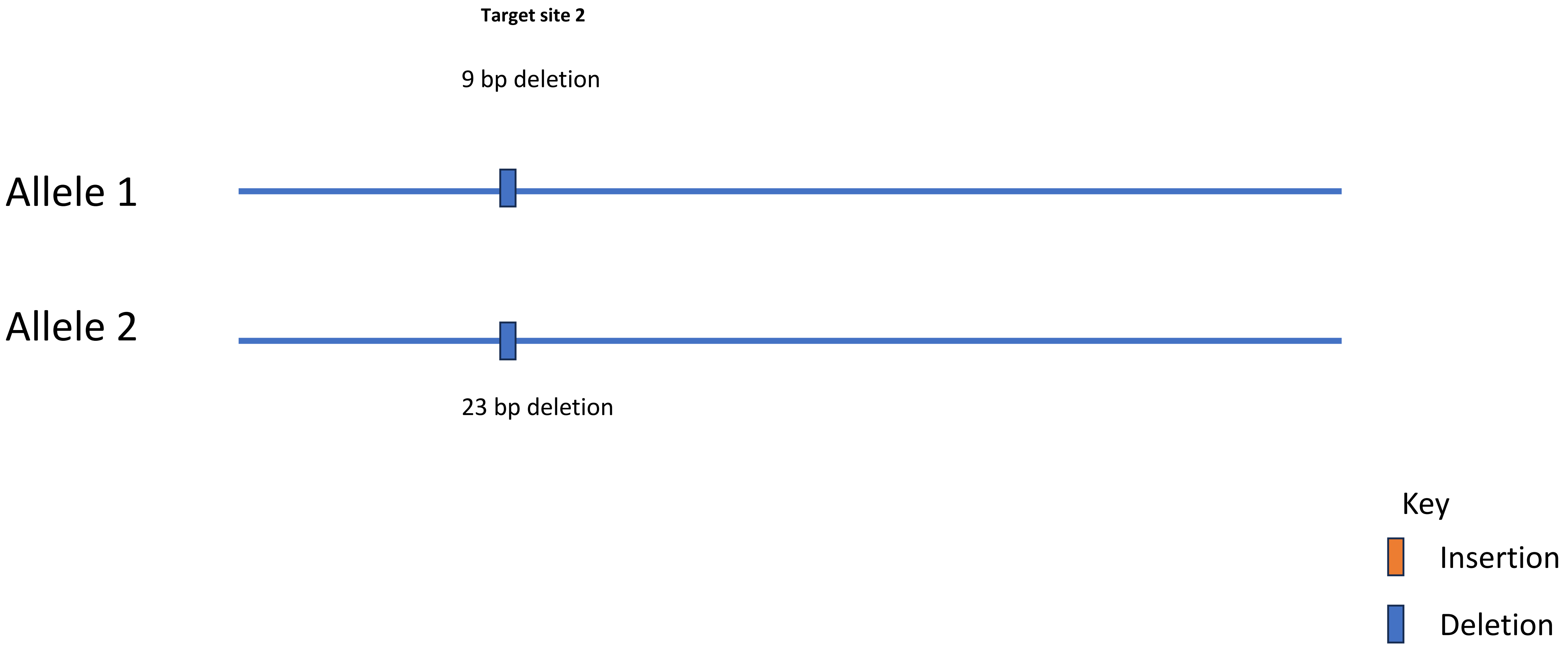
Target site 2
9 bp deletion
Allele 1
Allele 2
23 bp deletion
Key
Insertion
Deletion

# KI3

**Target site 2**

Deletion (9bp-GTGGTCGAT)-17reads

Allele 1

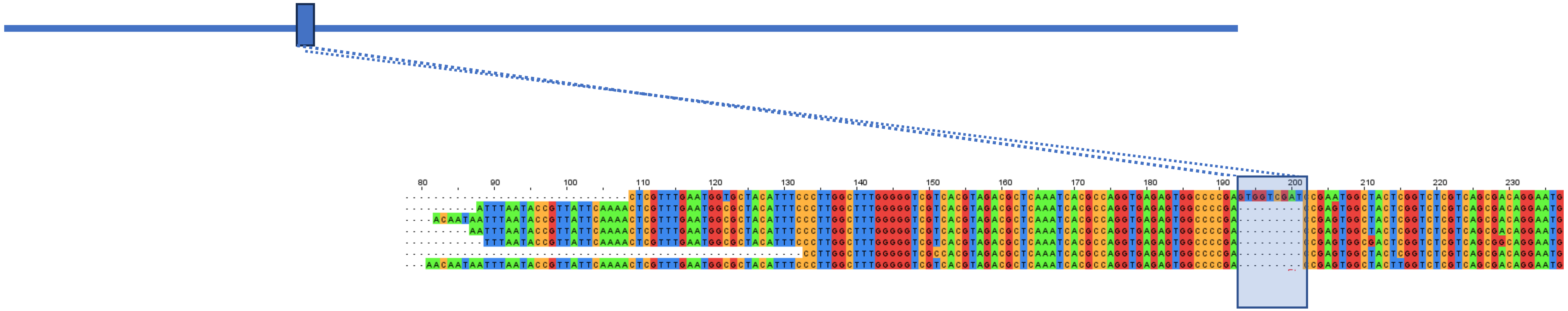

Key

Insertion

Deletion

# KI3

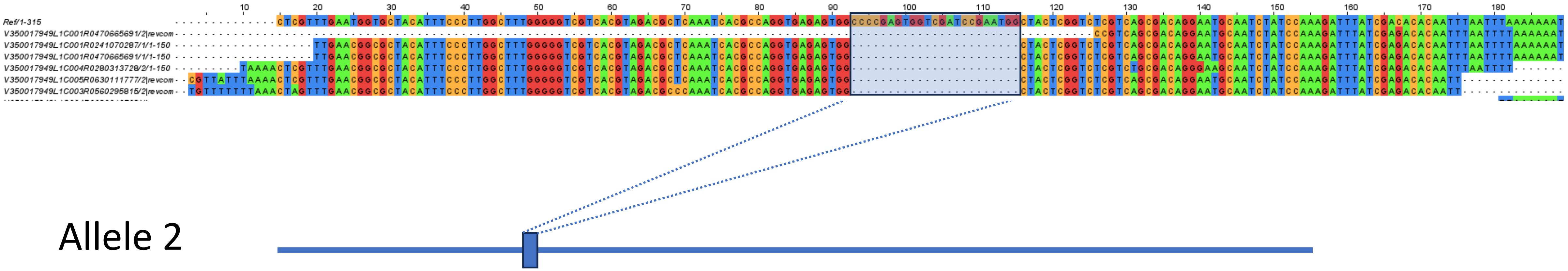

Allele 2

Deletion (23bp-CCCCGAGTGGTCGATCCGAATGG)-17 reads

Key

Insertion

Deletion
